# Gastrulation-stage gene expression in *Nipbl*^+/-^ mouse embryos foreshadows the development of syndromic birth defects

**DOI:** 10.1101/2023.10.16.558465

**Authors:** Stephenson Chea, Jesse Kreger, Martha E. Lopez-Burks, Adam L. MacLean, Arthur D. Lander, Anne L. Calof

**Affiliations:** Department of Developmental and Cell Biology, School of Biological Sciences, University of California Irvine, Irvine, CA 92697; Center for Complex Biological Systems, University of California Irvine, Irvine, CA 92697; Department of Quantitative and Computational Biology, Dornsife College of Letters, Arts, and Sciences, University of Southern California, Los Angeles, CA; Department of Anatomy and Neurobiology, School of Medicine, University of California Irvine, Irvine, CA 92697

## Abstract

In animal models, *Nipbl*-deficiency phenocopies gene expression changes and birth defects seen in Cornelia de Lange Syndrome (CdLS), the most common cause of which is *Nipbl*-haploinsufficiency. Previous studies in *Nipbl^+/-^* mice suggested that heart development is abnormal as soon as cardiogenic tissue is formed. To investigate this, we performed single-cell RNA-sequencing on wildtype (WT) and *Nipbl^+/-^* mouse embryos at gastrulation and early cardiac crescent stages. *Nipbl^+/-^*embryos had fewer mesoderm cells than WT and altered proportions of mesodermal cell subpopulations. These findings were associated with underexpression of genes implicated in driving specific mesodermal lineages. In addition, *Nanog* was found to be overexpressed in all germ layers, and many gene expression changes observed in *Nipbl^+/-^* embryos could be attributed to *Nanog* overexpression. These findings establish a link between *Nipbl*-deficiency, *Nanog* overexpression, and gene expression dysregulation/lineage misallocation, which ultimately manifest as birth defects in *Nipbl^+/-^* animals and CdLS.

**Teaser:** Gene expression changes during gastrulation of *Nipbl*-deficient mice shed light on early origins of structural birth defects.

## Introduction

Each year, 1 of every 33 babies in the United States is born with a birth defect (*1*), the most prevalent of which are congenital heart defects (CHDs), neural tube defects, and cleft lip/palate (*2*). Given the major impact that birth defects have on infant mortality and morbidity (*3, 4*), there is a need to elucidate their origins, but their diversity and sporadic nature pose challenges for identifying causal mechanisms. A promising approach is to study genetic syndromes that present multiple, concurrent defects in various body parts, many of which mirror common isolated birth defects. Studying them might thus provide insights into the causes and development of isolated birth defects.

Cornelia de Lange Syndrome (CdLS) affects an estimated 1 in 10,000 to 1 in 30,000 live births (*5*), and is characterized by craniofacial anomalies, delays in growth and maturation, intellectual disability, neurological impairments, abnormalities of limbs, especially arms and hands, coupled with issues in the visual, auditory, gastrointestinal, genitourinary, and cardiopulmonary systems (*6*).

The majority of CdLS cases – more than 55% – are caused by heterozygous mutations in the gene *Nipped-B-like* (*NIPBL*) (*7*), named for its homology to the Drosophila gene *Nipped-B*. These mutations often produce a non-functional protein, suggesting CdLS arises from haploinsufficiency (*8*). Remarkably, even a subtle 15% reduction in *NIPBL* gene expression can produce a mild yet recognizable CdLS phenotype (*9*). Such observations highlight the importance of precise *NIPBL* gene dosage in human development.

The *NIPBL* gene encodes a universally conserved protein that plays a role in loading cohesin onto chromosomes (*10*). Cohesin, similarly conserved and ubiquitous, is a four-subunit protein complex (Smc1, Smc3, Rad21, and either SA1 or SA2) essential for chromosome organization and genome stability (*11*). Mutations in the cohesin subunits *SMC1* and *SMC3* account for a small proportion of clinically mild CdLS (∼5% and <1%, respectively) (*12–14*). Additionally, mutations in *HDAC8*, which catalyzes cohesin release from chromatin during mitosis, are found in a distinct subset of CdLS patients (*15*). Mutations in *RAD21* have also surfaced in individuals exhibiting a CdLS-like phenotype but with substantially milder cognitive impairment (*16*). Collectively, these findings reinforce the idea that impairment of cohesin function contributes to CdLS.

Cohesin was initially identified for its role in sister chromatid cohesion during mitosis (*11*). However, pronounced defects in sister chromatid cohesion or irregularities in mitosis have not been observed in either CdLS patients or *Nipbl*-haploinsufficient (*Nipbl^+/-^*) mice (*17, 18*), suggesting that cohesin has additional functions. Studies in CdLS patient cells and *Nipbl*-haploinsufficiency animal models indicate that cohesin is involved in transcriptional regulation (*19*). Specifically, *Nipbl*-haploinsufficiency leads to alterations in the expression of many hundreds to thousands of genes (*18*). Interestingly, many of the affected genes are regulated by long-distance enhancers (*20*), aligning with the emerging concept of NIPBL and cohesin as critical determinants in DNA-looping (*21*).

Interestingly, most gene expression changes in *Nipbl*-deficient animals are small, usually less than 2-fold. Though likely to be inconsequential individually, such small changes can act collectively to produce structural and functional defects. For example, in zebrafish, joint depletion of two developmental genes downregulated by *nipbl*-deficiency produced a CdLS-like phenotype (*22*), suggesting that altered gene expression is the ultimate cause of developmental and physiological abnormalities. Thus, CdLS exemplifies a class of genetic disorders known as “transcriptomopathies” (*23*), in which the cumulative or synergistic effects of minor disturbances to gene expression lead to developmental abnormalities.

Previously, we reported that *Nipbl*-haploinsufficient mice displayed birth defects phenocopying those in CdLS (*18*). These included CHDs (primarily atrial septal defects), in about 30% of *Nipbl^+/-^* mice. Subsequently, we used a conditional *Nipbl* allelic series to investigate the role of *Nipbl* expression in the production of CHDs (*24*). That study showed that *Nipbl^+/-^* mice exhibit heart abnormalities early in development: At embryonic day 13.5 (E13.5), 70% of *Nipbl*^+/-^ mice displayed delays in ventricular septal fusion, whereas three days earlier, at E10.5, 100% showed right ventricle hypoplasia (*24*). In situ hybridization experiments showed reduced expression of two transcription factors crucial for early heart progenitor cell growth and differentiation: *Nkx2-5* and *Mesp1* (*24*). That structural abnormalities may begin at the earliest stages of heart development was suggested by results in in *nipbl*-morphant zebrafish, in which defects in the initial migration of cardiogenic mesoderm were detected as early as 18 hours post-fertilization (*22*). These results suggested that, at least for CHDs, causal events may occur as early as gastrulation, when the three primary germ layers form and the earliest progenitor cells of major tissues and organs begin to differentiate (*25*).

In this study, we sought to identify developmental alterations during gastrulation that might account for birth defects in *Nipbl*^+/-^ mice. We used single-cell RNA sequencing (scRNAseq) to compare the cellular compositions, lineage trajectories, and transcriptional landscapes of *Nipbl*^+/-^ mouse embryos to their WT counterparts at two stages spanning the end of gastrulation: the late bud-(LB) stage (E7.5) and cardiac crescent-(CC) stage (E7.75). Our findings reveal that *Nipbl*^+/-^ embryos have the same cell populations as WT embryos but display subtle misallocation of specific mesodermal cell populations. Our evidence suggests this occurs as a result of alterations in specific cell fate decisions, including the choice by mesoderm cells to progress toward a non-cardiac versus cardiac fate. Our observations strongly suggest these events cannot be attributed to changes in apoptosis, cell proliferation, global developmental delay, or the structure of cell lineages. As in earlier research, we observed that most gene expression changes in *Nipbl*-deficient tissues at these stages were small (*18*), but we also identified several key developmental genes that are more markedly misexpressed in *Nipbl*^+/-^ embryos, the most notable of which was *Nanog*. *Nipbl*^+/-^ embryos failed to downregulate *Nanog* at the end of gastrulation, which normally occurs in all but germ cells. As a result, the misexpression of many *Nanog* target genes was observed in all germ layers. We also saw substantial underexpression of *Hox* genes and overexpression of Nodal signaling genes, which play roles in anterior-posterior and left-right patterning, respectively. As a result, we propose a model in which birth defects in CdLS arise from the prolonged overexpression of *Nanog* and dysregulation of developmental pathways governing axial patterning, resulting in the misdirection of cell fate decisions, and misallocation of specific progenitor cell populations.

## Results

### Gastrula-stage *Nipbl*^+/-^ mice display the same cell populations as are found in WT mice

To investigate early factors influencing birth defects in CdLS, we generated WT and *Nipbl*^+/-^ littermate embryos by crossing *Nanog^Cre/+^* mice (*26*) with *Nipbl^Flox/Flox^* mice (*24*) as described in Methods and shown in Fig. 1A. To ensure an accurate accounting of cell populations and proportions at well-defined stages of early embryonic development, we generated a large excess of embryos and sorted them into groups of narrowly defined stage based on morphological criteria (*27*). Using the 10X Genomics Chromium Single Cell Expression platform (*28*), we performed scRNAseq on 5 WT and 6 *Nipbl^+/-^* samples of late bud stage (LB) embryos (E7.5), and 8 WT and 8 *Nipbl^+/-^* samples of cardiac crescent stage (CC) embryos (E7.75) (Fig. 1B and fig. S1). Given the low cell numbers in individual LB-stage embryos, and the 10X Genomics Chromium Single Cell Expression system’s capture efficiency, samples at the LB stage consisted of pairs of embryos, whereas at the CC-stage they consisted of single embryos. Pairing LB-stage embryos facilitated the capture of rare cell groups, such as primordial germ cells, which typically comprise about 50 cells per embryo at this stage (*29*).

**Fig. 1.**
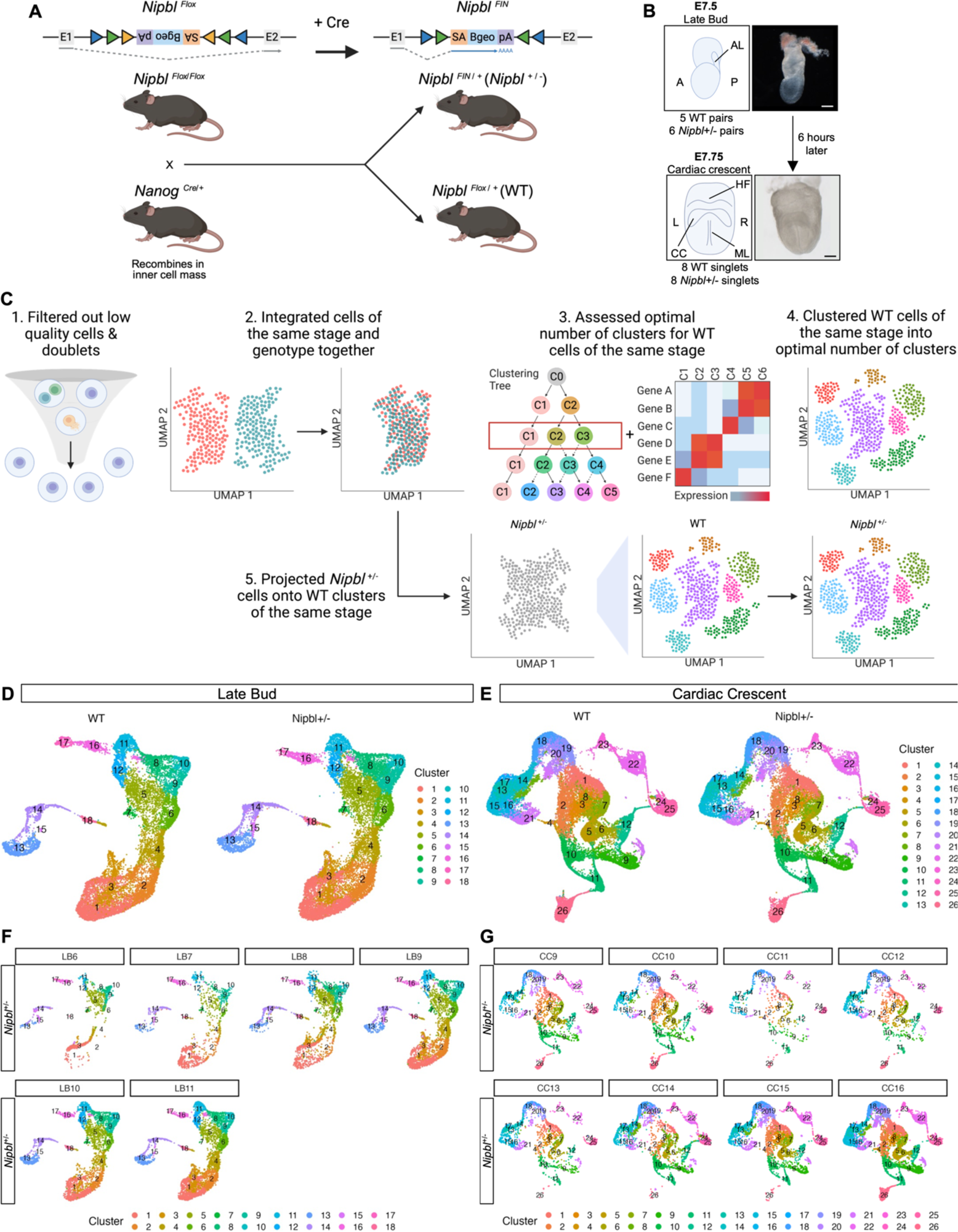
*Nipbl*^+/-^ mice do not lack any cell populations found in WT mice. (**A**) *Nipbl* alleles used in this study. *Nipbl*^Flox^ contains an inverted gene trap cassette encoding *β-geo* flanked by *Cre* recombinase target sites in intron 1 of *Nipbl* gene alleles (*24*). In this inverted orientation, there is no trapping of the *Nipbl* gene and *Nipbl* is expressed normally. However, when this cassette is exposed to Cre recombinase, the gene trap cassette gets inverted producing the *Nipbl*^FIN^ allele. In this orientation, trapping of the *Nipbl* gene occurs and *β-geo* is expressed as a reporter of successful gene trapping. When *Nanog*^Cre/+^ mice are mated with *Nipbl*^Flox/Flox^ mice, the resulting littermates are either entirely *Nipbl*^Flox/+^ or entirely *Nipbl*^FIN/+^, as *Nanog*^Cre/+^ mice carry a transgene that expresses *Cre* recombinase in the earliest cells of the developing embryo (*26*). (**B**) Lateral view of LB-stage and anterior view of CC-stage embryos subjected to scRNAseq. A = anterior, P = posterior, AL = allantois, L = left, R = right, HF = head fold, CC = cardiac crescent, ML = midline. Dashed line represents where embryonic tissue was separated from extraembryonic tissue. Scale bar = 100 microns. (**C**) Workflow used to filter out low quality cells and doublets, cluster WT cells into optimal number of clusters at each stage, and project *Nipbl^+/-^* cells onto WT clusters of the same stage. Uniform manifold approximation and projection (UMAP) of clusters in WT and *Nipbl^+/-^* embryos at (**D**) LB- and (**E**) CC-stage. UMAP of clusters in each *Nipbl*^+/-^ embryo at (**F**) LB- and (**G**) CC-stage.

At LB-stage, we captured a median of 2,537 cells per sample, with a median of 17,301 RNA transcripts per cell (fig. S2 and table S1). At CC-stage, we captured a median of 4,116 cells per sample, with a median of 14,997 RNA transcripts per cell (table S1). As expected, all *Nipbl*^+/-^ embryos expressed both *Cre* and *Betageo*, which report an inactivated *Nipbl* allele (*24*), while WT embryos did not (fig. S3, A and B). *Nipbl*^+/-^ embryos across both stages expressed *Nipbl* at levels that were approximately 50% lower than WT counterparts (fig. S3C).

We corrected for batch effects among embryos of identical stage and genotype using Seurat’s integration protocol (*30*) (Fig. 1C and fig. S4), which identifies anchors (cells of similar gene expression across samples of the same biological condition) and uses them to align cells into a space shared by all samples. Recognizing that all cells in *Nipbl*^+/-^ tissues exhibit substantial gene expression changes (*18*), and considering that clustering algorithms rely on differences in gene expression among cells (*30*), we implemented measures to prevent our cell clustering from being skewed by gene expression alterations attributable to *Nipbl*-haploinsufficiency. We did this by first clustering cells from WT samples (Fig. 1C), using a robust iterative clustering method that considered both intra-cluster stability and inter-cluster variation (fig. S5) to determine the optimal number of clusters for cells at each developmental stage. WT cells were clustered into 18 cell populations at LB-stage (Fig. 1D) and 26 cell populations at CC-stage (Fig. 1E). We then projected *Nipbl*^+/-^ cells onto these WT cell populations at corresponding stages (Fig. 1C). We observed that all *Nipbl*^+/-^ cells could be projected to WT cell populations, with each WT population receiving some *Nipbl*^+/-^ cells (Fig. 1, D and E). This pattern held true across individual *Nipbl*^+/-^ samples, as well as the total *Nipbl*^+/-^ cells in aggregate at each stage (Fig. 1, F and G). To ensure that any differences observed in cell populations were not the result of technical artifacts introduced by the projection algorithm, we performed additional studies in which WT cells were projected onto *Nipbl*^+/-^ clusters (fig. S6 and fig. S15). In both cases, *Nipbl*^+/-^ cells contributed to all clusters, but as we describe later, in varying proportions (Fig. 3).

### scRNAseq captured cells from all germ layers and progenitors of major tissues

To assign biological identities to clustered cell populations, we used WT cells as a reference. For each WT cell population, we performed differential gene expression analysis (DGEA) by comparing it to all other cells within embryos of identical stage (Mann-Whitney U test). We detected markers indicative of germ layer identity in every cluster (data S1, data S2, and table S2), enabling us to associate each cluster with a specific germ layer at each developmental stage (Fig. 2A). Figure 2B displays all cells from all samples, categorizing them into germ layers at both LB- and CC-stages of development. Figures 2C and 2D further depict the assignment of cells based on the expression of germ layer markers, at LB- and CC-stages, respectively. These markers include those for ectoderm, such as *Utf1, Sox2,* and *Pou3f1* (*31–33*), mesoderm, including *T, Hand1, Vim, Twist1,* and *Prrx2* (*34–38*), and endoderm, *Ttr* (*39*).

**Fig. 2.**
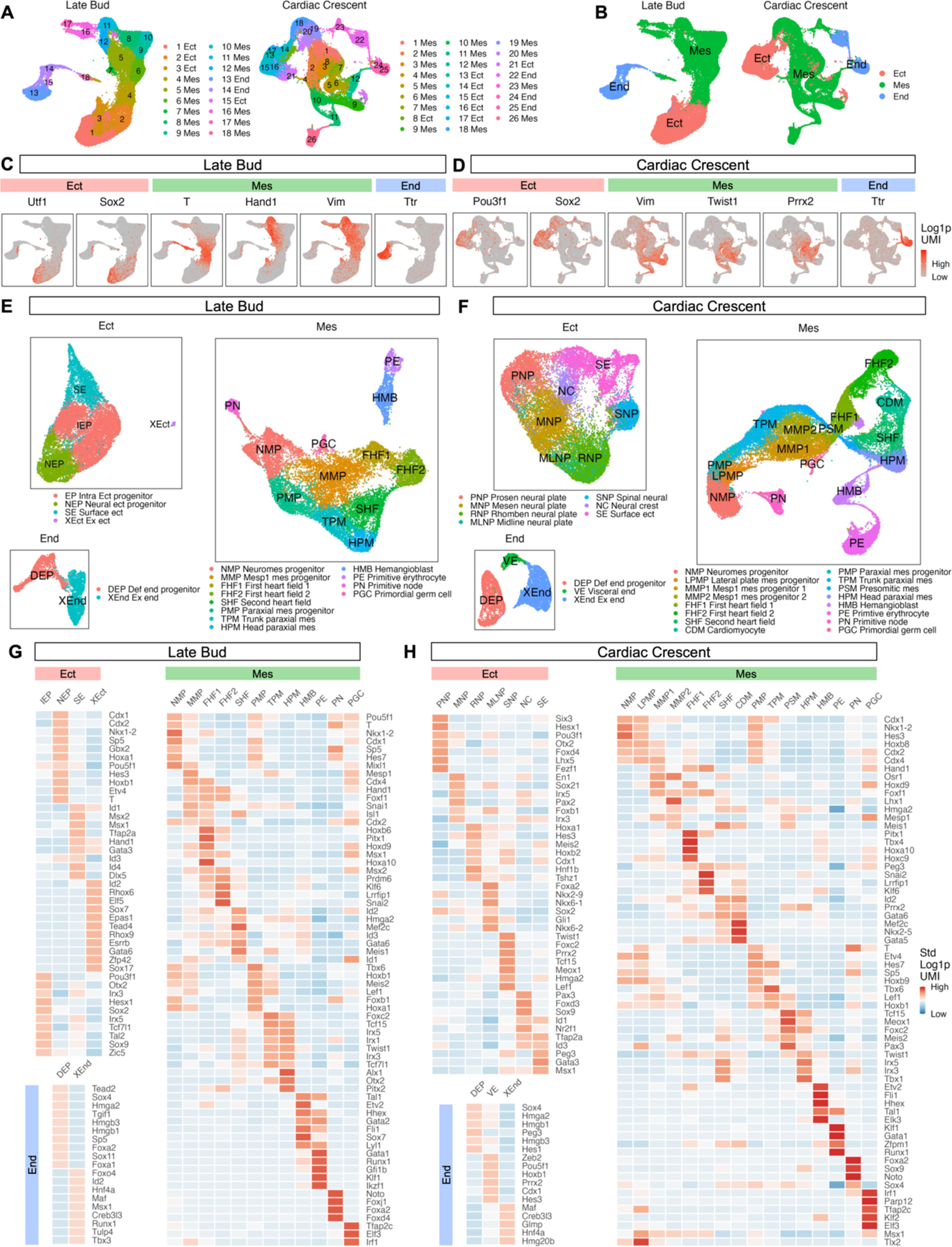
scRNAseq captured cells from all germ layers, as well as progenitors of major tissues. **(A)** UMAP of clusters assigned to germ layers in LB- and CC-stage embryos. (**B**) UMAP of germ layers in LB- and CC-stage embryos. Expression of genes marking germ layers of (**C**) LB- and (**D**) CC-stage embryos in UMAP. UMAP of cell populations in germ layers of (**E**) LB- and (**F**) CC-stage embryos. Heatmap of fold change in expression of the most differentially expressed transcription factor genes (lowest *Q*-values from Mann-Whitney U Test) (*101*) of cell populations from all other cell populations in germ layers of (**G**) LB- and (**H**) CC-stage embryos.

To identify cell types within germ layers, we re-applied DGEA after clustering individual germ layers. For the endoderm, this enabled us to identify definitive endoderm progenitors (DEP) (*Sox4* and *Foxa2*) (*40, 41*), as well as extraembryonic endoderm (XEnd) (*Hnf4a* and *Tbx3)* (*42, 43*) at LB-stage (Fig. 2E, data S3, and table S3). In CC-stage embryos, we captured identical cell types in the endoderm, and additionally, visceral endoderm (VE) (*Hnf4a* and *Esx1*) (*42, 44*)(Fig. 2F, data S4, and table S3). DEPs and XEnd give rise to the gut and placenta, respectively.

Within the ectoderm of LB-stage embryos, we identified early progenitor populations including intraembryonic ectoderm progenitors (IEPs) (*Pou3f1* and *Otx2*) (*33, 45*), neural ectoderm progenitors (NEPs) (*Cdx1* and *Nkx1-2)* (*46, 47*), and surface ectoderm (SE) (*Id1, Msx2,* and *Msx1*) (*48*) (Fig. 2, E and G, data S5, and table S4). Additionally, we detected a small portion of extraembryonic ectoderm (XEct) (*Elf5* and *Tead4*) (*49, 50*), which was expected, as a small quantity of XEct cells was included in our dissection. By CC-stage, IEPs, NEPs, and XEct were no longer detectable, a phenomenon previously noted (*51*). Rather, we continued to detect surface ectoderm (SE). We also detected six distinct neural populations (Fig. 2, F and H, data S6, and table S4). These neural populations included: prosencephalic neural plate (*Six3* and *Hesx1*) (*52, 53*), mesencephalic neural plate (*En1* and *Pax2*) (*54, 55*), rhombencephalic neural plate (*Hoxa1* and *Hes3*) (*56, 57*), midline neural plate (*Foxa2* and *Nkx2-9*) (*58, 59*), spinal neural plate (*Sox2* and *Ezr*) (*60, 61*), and neural crest (*Pax3* and *Foxd3*) (*62, 63*) (data S6). Collectively, these populations constitute the nascent progenitors of the brain, spinal cord, and peripheral nervous system.

Within the mesoderm of LB-stage embryos, we identified six distinct groups of cells, detailed in Fig. 2, E and G, data S7, and table S5. These consisted of: 1) neuromesodermal progenitors (NMPs): expressing markers *T,* and *Nkx1-2* (*34, 47*); 2) derivatives of NMPs: including *Mesp1*-expressing mesoderm progenitors (MMPs) (*Mesp1*) (*64*), two types of first heart field cells (FHF1 and FHF2) marked by *Hand1* (*35*); cells of the second heart field (SHF) with *Id2* and *Mef2c* (*65, 66*); and three kinds of paraxial mesoderm cells, further categorized into paraxial mesoderm progenitors (PMPs) (*Tbx6* and *Meis2)* (*67, 68*), trunk paraxial mesoderm (TPM) (*Foxc2* and *Tcf15*) (*69, 70*), and head paraxial mesoderm (HPM) (*Alx1* and *Tcf15*) (*70, 71*); 3) hematopoietic cells: including hemangioblasts (HMBs) (*Tal1* and *Etv2*) (*72, 73*) and primitive erythrocytes (PEs) (*Gata1* and *Tal1*) (*72, 74*); and 4) two rare populations: comprising cells of the primitive node (PN) (*Noto* and *Foxj1*) (*75, 76*), and primordial germ cells (PGCs) (*Tfap2c* and *Msx1*) (*77, 78*). At CC-stage, we detected all mesodermal populations found earlier, along with several that emerge later in development. These comprised lateral plate mesoderm progenitors (LPMP) (*Tlx2* and *T*) (*34, 79*), additional populations of *Mesp1*-expressing mesoderm progenitors (MMP1 and MMP2) (*Mesp1*) (*64*), cardiomyocytes (CDMs) (*Mef2c* and *Nkx2-5*) (*80, 81*), and presomitic mesoderm (PSM) (*Tcf15* and *Meox1*) (*70, 82*) (Fig. 2, F and H, data S8, and table S5). Collectively, these progenitors are responsible for the formation of structures including skeletal muscle, bone, blood, and heart.

### *Nipbl^+/-^* mice have fewer mesoderm cells, primitive erythrocytes, and first heart field cells, and more paraxial mesoderm cells

We analyzed the cellular composition of embryos at both LB- and CC-stage, quantifying the proportion of cells within each germ layer that contributed to the total cell count. LB-stage mutants exhibited 13% fewer mesoderm cells (Fig. 3, A and B, data S9), while concurrently displaying a greater proportion of endoderm and ectoderm cells. We sought to ascertain if this decrease in mesoderm was generalized or restricted to specific subpopulations. Focusing on LB-stage embryos, we merged mesodermal cell subpopulations of similar biological identity together (FHF1 + FHF2 into FHF, and PSM + CPM into PM) (Fig. 3C) and calculated the percentage of cells across these subpopulations, relative to the overall mesoderm. The findings (Fig. 3D) revealed a 77% reduction in PEs *Nipbl*^+/-^ embryos (*P*=0.06, T-Test, data S10). Additionally, differences were observed in two mesodermal derivatives of the neuromesoderm: 1) Mutants had 24% fewer first heart field cells (FHF) (Fig. 3D); and 2) *Nipbl*^+/-^ embryos showed a contrasting pattern in paraxial mesoderm (PM), a 33% increase in PM cells compared to WT (Fig. 3D).

**Fig. 3.**
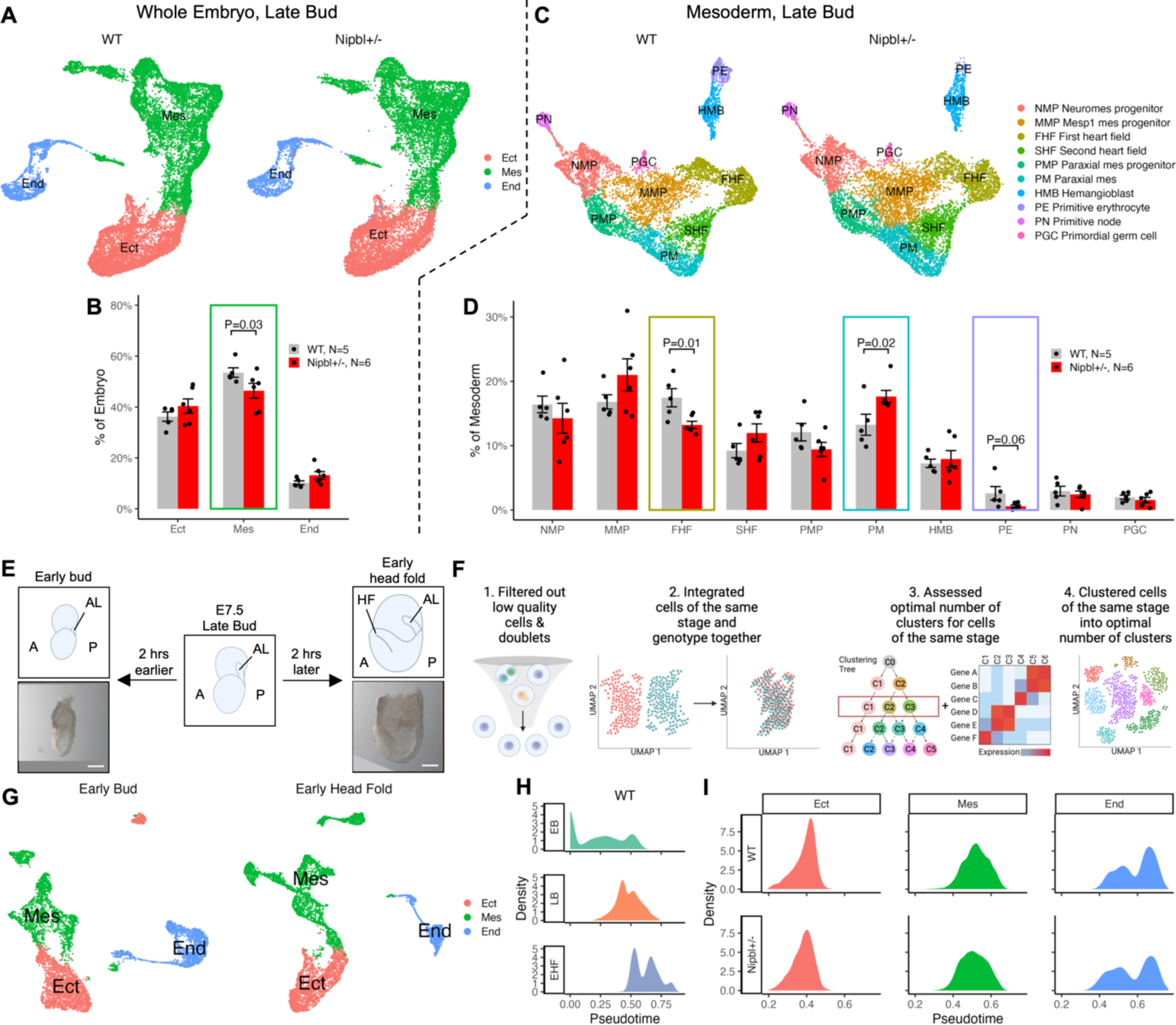
*Nipbl^+/-^*embryos exhibit changes in the sizes of mesodermal subpopulations that are not accompanied by changes in developmental timing. (**A**) UMAP of germ layers in LB-stage WT and *Nipbl*^+/-^ embryos. (**B**) Percentage of cells in germ layers from all cells in LB-stage WT and *Nipbl^+/-^* embryos. Error bars show standard error of the mean. *P*-values from T-Test. (**C**) UMAP of cell populations in mesoderm of LB-stage WT and *Nipbl^+/-^* embryos. (**D**) Percentage of cells in mesodermal cell populations from all cells in LB-stage WT and *Nipbl^+/-^* embryos (bottom). Error bars show SEM. *P*-values from T-Test. (**E**) Lateral view of WT EB- and EHF-stage embryos subjected to scRNAseq. A = anterior, P = posterior, AL = allantois, L = left, R = right, HF = head fold. Dashed line represents where embryonic tissue was separated from extraembryonic tissue. Scale bar = 100 um. (**F**) Workflow used to filter out low quality cells and doublets and cluster cells at EB- and EHF-stage into optimal number of clusters. (**G**) UMAP of clusters assigned to germ layers in EB- and EHF-stage embryos. (**H**) Density of cells from WT EB-, LB-, and EHF-stage embryos along pseudotime (calculated using URD). (**I**) Density of cells in germ layers of LB-stage WT and *Nipbl*^+/-^ embryos along pseudotime.

Since *Nipbl^+/-^* cells were projected onto WT cell populations, we wondered whether any observed differences between *Nipbl^+/-^* and WT embryos might have occurred as a result of misprojecting cells, i.e. assigning cells to incorrect clusters. To investigate this, we independently clustered *Nipbl^+/-^* cells and projected WT cells onto those clusters; we refer to this as reverse projection (fig. S6). If observed differences in the allocation of *Nipbl^+/-^* cells to particular clusters had been a result of misclassification due to technical limitations of projecting cells, then the same differences should not appear when reverse projection was done. Indeed, if misclassified cells were simply distributed at random one should expect reverse projection to produce differences opposite to those produced by the original, forward projection. This was not found to be the case, however. Cluster assignments resulting from reverse projection quantitatively resembled those of forward projection (fig. S6, A to D). Germ layer identities, which were based on the expression of specific markers (table S6 and data S11), were generally the same in the reverse projection analysis as well (fig. S6, E to H).

Finally, we quantified the proportion of cells within each germ layer and mesodermal population following reverse projection. Consistent with the findings from forward projection (Fig. 3B), *Nipbl^+/-^* embryos showed 13% fewer mesoderm cells than WT embryos, while concurrently showing more endoderm and ectoderm cells (fig. S6M and data S13). Likewise, the mesoderm of *Nipbl^+/-^* embryos showed fewer FHF cells (24% fewer) and more PM cells (22% more) than WT embryos (Fig. 3D, fig. S6N, and data S14). (Because very few PEs are present in *Nipbl^+/-^* embryos (Fig. 3D), it was not possible to cluster these cells into a distinct cell population during reverse projection (fig. S6, K and L).) Altogether, these data indicate that differences in cell population sizes observed in *Nipbl^+/-^* embryos represent true biological changes and are not a consequence of technical issues, such as misprojection.

### *Nipbl^+/-^* embryos do not show changes in developmental timing

Cellular composition changes very quickly during and after gastrulation; one possible explanation for differences in proportions of cell types could be a change in the overall pace of development, such that embryos of one genotype were slightly delayed or accelerated relative to the other. To investigate whether this was the case for *Nipbl^+/-^* embryos, we employed pseudotemporal ordering. To ensure that pseudotime accurately mirrored real developmental time, we augmented our analysis with scRNAseq data from an additional two pairs of WT embryos at the early bud (EB) stage and two pairs of WT embryos at the early head fold (EHF) stage (Fig. 3E and fig. S7). EB-stage embryos are two hours younger and EHF-stage embryos are two hours older than LB-stage embryos, so these supplementary samples provided the temporal resolution necessary to discern even minor changes (less than 2 hours) in developmental timing (*27*). We eliminated low-quality cells and doublets (Fig. 3F, fig. S8 and table S8) before integrating cells of the same stage (fig. S9). We then clustered EB-stage cells into 8 distinct populations and EHF-stage cells into 13 (fig. S10, A and B). Each population was subsequently annotated as ectoderm (Ect), mesoderm (Mes), or endoderm (End) (Fig. 3G, and fig. S10, C and D) based on the expression of specific germ layer markers (data S15 and S16, and table S9).

We ordered cells from each embryo at every developmental stage (EB, LB, and EHF) using URD (*83*), which constructs a diffusion map of transition probabilities and, starting with an assigned group of root cells, performs a probabilistic breadth-first graph search using the transition probabilities. When we visualized the arrangement of cells from each stage based on their density in pseudotime, we found that cells from WT embryos ordered in accordance to their developmental stage: EB came first, followed by LB, and finally EHF (Fig. 3H, data S17), while showing partial overlap between cells of different stages. This confirmed that the pseudotime values we acquired were an accurate representation of actual developmental timing. When the pseudotime orderings of WT and *Nipbl^+/-^*LB-stage embryos were compared with each other, we found no statistically significant deviations (Kolmogorov-Smirnov test) (Fig. 3I). These findings argue that the observed reduction in mesoderm cells in *Nipbl^+/-^* mice at LB-stage cannot be attributed to overall developmental delay or acceleration. The fact that *Nipbl*-haploinsufficiency does not result in global changes in developmental timing strongly suggests that birth defects in CdLS result from specific cellular or molecular irregularities within individual developmental pathways, and not a globally-altered developmental timeline.

### *Nipbl^+/-^* embryos do not show changes in cell lineage trajectory

To discern whether alterations in the sizes of cell population in *Nipbl^+/-^* embryos were attributable to changes in the structures of cell lineages, we used scVelo (*84*) to compute the RNA velocity for all mesoderm cells from LB-stage embryos. RNA velocity provides a predictive metric for a cell’s future transcriptional state, gauging the equilibrium between the synthesis of spliced mRNA from unspliced mRNA and mRNA degradation. scVelo was used to build cell lineage trajectories separately for WT and *Nipbl^+/-^* embryos at LB-stage (Fig. 4A). In WT mesoderm, five distinct lineages were discernible (Fig. 4B, data S18), with neuromesodermal progenitors (NMPs) giving rise to three terminal fates via four specific pathways. These pathways encompassed the differentiation of NMPs into FHF, SHF, and PM. It’s intriguing to note that *Mesp1*-expressing cells have the versatility to differentiate into both cardiac and paraxial mesoderm. While *Mesp1* is typically recognized as a cardinal factor for cardiac specification (*85*), our findings suggest its influence may extend beyond this role into paraxial specification as well. As a result, the paraxial mesoderm (PM) has dual progenitors, PMPs and MMPs. We also observed a fifth lineage, independent of NMP lineages, where HMBs differentiate into PEs. Notably, PGCs and PN cells did not form part of any identified lineages. Examination of the mesoderm in *Nipbl^+/-^* embryos revealed lineage trajectories identical to those in WT (Fig. 4A, data S19). These observations suggest that variations in the overall structures of cell lineage pathways do not explain differences in the number of FHF and PM cells in *Nipbl^+/-^* embryos (Fig. 3, B and D).

**Fig. 4.**
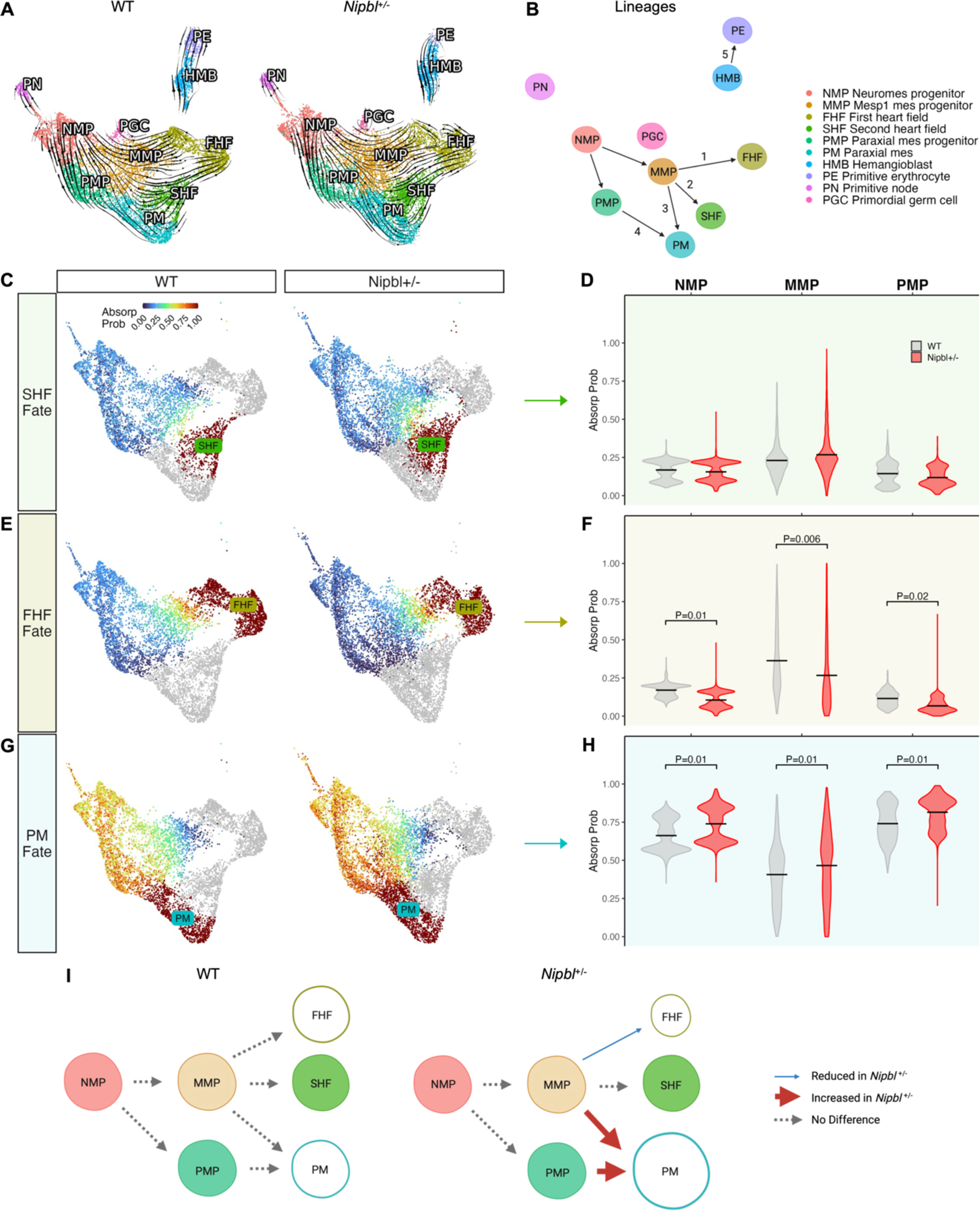
*Nipbl^+/-^*mice misdirect mesoderm cells into paraxial mesoderm at the expense of the first heart field. (**A**) RNA velocities, calculated by scVelo of mesoderm cells from LB-stage WT and *Nipbl^+/-^* embryos in UMAP. (**B**) Cell lineage trajectories in mesoderm of LB-stage WT and *Nipbl^+/-^* embryos. Probability of mesoderm cells from LB-stage WT and *Nipbl^+/-^* embryos terminally transitioning (absorption probabilities calculated using CellRank) into (**C**) SHF, (**E**) FHF, or (**G**) PM fates in UMAP. Violin plot of probability of NMPs, MMPs, and PMPs from LB-stage WT and *Nipbl^+/-^* embryos terminally transitioning (absorption probabilities) into (**D**) SHF, (**F**) FHF, or (**H**) PM fates. Lines show means. *P*-values from T-Test. (**I)** Schematic illustrating how mesoderm cells in *Nipbl^+/-^* embryos are misdirected into PM fate at the expense of FHF fate.

### *Nipbl^+/-^* embryos misdirect mesoderm cells into paraxial mesoderm at the expense of the first heart field

To explore whether variations in cell fate decisions might be responsible for the observed differences in the numbers of FHF and PM cells in *Nipbl^+/-^* embryos, we employed CellRank (*86*) to compute the likelihood of cells from the mesoderm (excluding HMBs, PEs, PGCs, and PN cells) transitioning terminally into cells of SHF, FHF, or PM fate (data S20 and data S21). HMBs, PEs, PGCs, and PN were omitted from the calculation because our trajectories derived from RNA velocity indicated that these populations do not ultimately transition into SHF, FHF, or PM. In CellRank, the absorption probability defines the likelihood that a specific cell type will make a transition to a terminal state, derived using RNA velocity-directed, random walks from initial to terminal cell states (*86*). We represented these probabilities in UMAP space, with shades of blue to signify low probabilities and red for high probabilities. Fig. 4, C and D illustrates that the probabilities of NMPs, MMPs, and PMPs making a terminal transition solely into SHF do not vary significantly between *Nipbl^+/-^*and WT embryos, as quantified in the adjacent violin plots. However, the transition probabilities into FHF are 27-48% lower in *Nipbl^+/-^* embryos (Fig. 4, E and F), while the probabilities for transitions into PM are 9-11% higher (Fig. 4, G and H). Taken together, these results support the hypothesis that disparities in cell numbers in *Nipbl^+/-^* embryos stem from a misdirection of cells from FHF pathway towards a PM fate (Fig. 4I).

### *Nipbl^+/-^* embryos do not show evidence of altered apoptotic activity or cell proliferation

Altered cell numbers in different mesodermal subpopulations of *Nipbl^+/-^*embryos could conceivably also stem from modifications in rates of cell death or proliferation. Focusing on apoptotic activity, we carried out differential gene expression analysis (DGEA) on WT and *Nipbl^+/-^* embryos at LB-stage, using genes associated with apoptosis from two well-known gene sets: Reactome Apoptosis and Hallmark Apoptosis (*87, 88*). We ranked each gene by minimum Q-value, arranging them from lowest to highest, and visualized their expression across all germ layers in *Nipbl^+/-^* embryos as heatmaps of the fold change relative to their WT counterparts (fig. S11, A and B, and data S22). For both gene sets, we saw no visible pattern of differences in the expression of genes with even the lowest Q-values between any of the germ layers in both WT and *Nipbl^+/-^* embryos. Although genes from these gene sets tend to mark cells with the capacity for apoptosis, their expression levels are not necessarily indicative of cells actively undergoing apoptosis. We therefore also turned to a gene set identified in a study that compared 180 apoptosis-associated genes in hematopoietic cells that were either healthy or undergoing apoptosis (*89*), and identified 93 apoptosis-associated genes that were differentially expressed. When we examined these genes in the germ layers of *Nipbl^+/-^*embryos, we also saw no visible difference with WT embryos (fig. S11C and data S22). These data suggest that changes in apoptotic activity are unlikely to be a major contributor to differences in cell type proportions.

To examine cell proliferation, we again employed DGEA across all germ layers of both WT and *Nipbl^+/-^* embryos at LB-stage, focusing on the meta-PCNA gene set, an ensemble of genes that exhibit the strongest positive correlation with PCNA expression, a recognized biomarker of cell proliferation (*90*). When we visualized the expression of these genes in the same manner as the gene sets above, we found no discernible variations in the expression levels of the meta-PCNA genes between the germ layers of WT and *Nipbl^+/-^* embryos (fig. S12A and data S23). As another measure of cell proliferation, we used Seurat to assign cells from LB-stage WT and *Nipbl^+/-^* embryos into phases (G1, S, and G2/M) of the cell cycle (*91*). Seurat does this by calculating a cell cycle phase score based on the expression of canonical S phase and G2/M phase markers. Seurat considers these marker sets to be anticorrelated in their expression, so when cells express neither, they are considered to be in G1 phase. As expected, in all germ layers, cells from LB-stage WT and *Nipbl^+/-^*embryos assigned into all phases of the cell cycle (fig. S12B and data S24). In all germ layers, there was no statistically significant difference in the proportions of cells in each cell cycle between WT and *Nipbl^+/-^* embryos (fig. S12C). We therefore concluded that extensive alterations in cell proliferation are not likely responsible for disparities in cell subpopulations.

### *Nipbl*^+/-^ mice underexpress genes predicted to drive the transition of mesoderm cells into first heart field

Next, we sought to investigate whether the misdirection of mesoderm cells in *Nipbl^+/-^* embryos at the LB-stage could be attributed to the misexpression of genes that drive the transition of mesoderm cells into either FHF or PM. To answer this question, we also employed CellRank to identify potential driver genes of the FHF and PM fates, using those mesoderm cells that specifically contribute to these developmental pathways (Fig. 5, A and D). CellRank achieves this by calculating a correlation coefficient between the likelihood (absorption probabilities) of cells progressing towards a particular lineage fate and the expression level of individual genes (*86*). In this context, genes with positive correlation coefficients are considered drivers, as their expression elevates alongside an increase in absorption probabilities. Conversely, genes with negative correlation coefficients are deemed anti-drivers, as their expression diminishes with increasing absorption probabilities. To minimize the risk of false discoveries, we set a threshold, accepting genes with correlation coefficients greater than 0.25 as drivers and those less than -0.25 as anti-drivers.

**Fig. 5.**
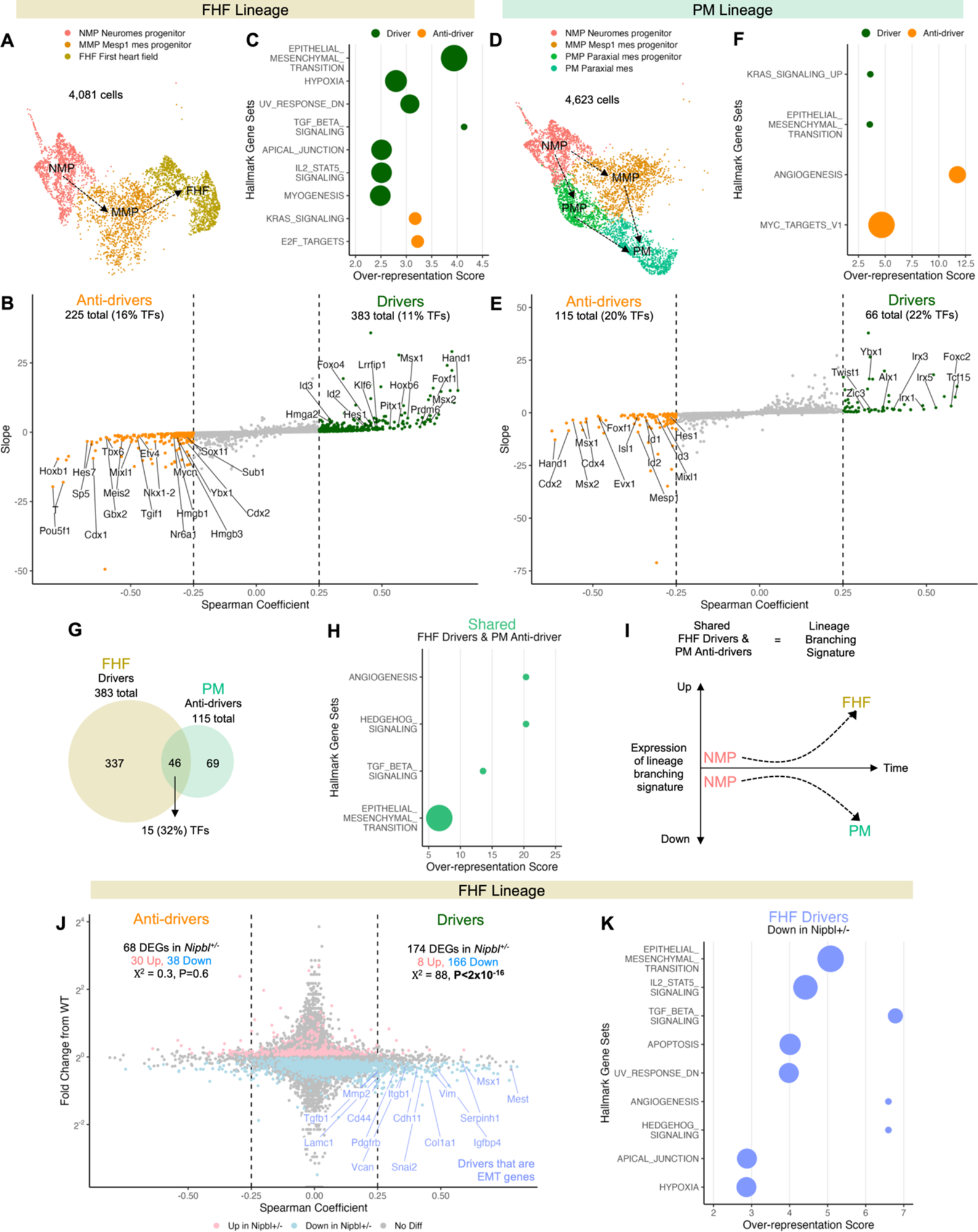
*Nipbl*^+/-^ mice underexpress genes predicted to drive the transition of mesoderm cells into first heart field. UMAP of mesoderm cells in (**A**) FHF and (**D**) PM lineages of WT LB-stage embryos. Genes whose expression are positively correlated (drivers) or negatively correlated (anti-drivers) with the transition (absorption probabilities from Fig. 3J) of mesoderm cells from WT LB-stage embryos into (**B**) FHF or (**E**) PM fates. Correlation coefficients calculated using Spearman’s Rank Correlation. Genes with correlation coefficient greater than 0.25 or less than -0.25 were considered drivers and anti-drivers, respectively. Slope refers to change in gene expression along absorption probability. Over-representation score of MSigDB’s Hallmark gene sets among drivers and anti-drivers of (**C**) FHF and (**F**) PM fate. (**G**) Venn diagram of genes shared between PM anti-drivers and FHF drivers. (**H**) Over-representation score of MSigDB’s Hallmark gene sets among shared FHF drivers and PM anti-drivers. (**I**) Expression of lineage branching signature as NMPs transition into FHF or PM fates. (**J**) Fold change in expression of genes in mesoderm cells of FHF lineage of LB-stage *Nipbl*^+/-^ embryos differentially expressed (*Q* < 0.05, Mann-Whitney U Test) from that of WT embryos along Spearman’s Rank Correlation coefficient from Fig. 5B. Genes associated with EMT are labeled. (**K**) Over-representation score of MSigDB’s Hallmark gene sets among shared FHF drivers downregulated in FHF lineage of *Nipbl^+/-^* mice.

CellRank identified 383 genes as drivers of FHF fate (Fig. 5B and data S25). 11% of them were transcription factors. Among these were *Hand1* and *Tbx20*, transcription factors previously recognized for their roles in promoting FHF development (*35, 92*). Several transcription factors that have not been previously linked to FHF development were also predicted to be FHF drivers, including *Msx1, Msx2, Foxf1, Hoxb6,* and *Pitx1*. Of these, *Hoxb6* and *Pitx1* are known to play important roles in segmentation and patterning. Gene set over-representation analysis using MSigDB’s Hallmark gene sets (*87*), which characterize well-defined biological states and processes, revealed that FHF drivers were significantly enriched in genes associated with TGF-beta signaling (Fig. 5C), which has been previously described as important in cardiomyocyte proliferation (*93*). Genes associated with epithelial to mesenchymal transition (EMT) were also significantly enriched (see below). Conversely, CellRank predicted 225 genes to be anti-drivers of FHF fate, with 16% being transcription factors. Among these were genes known to be involved in pluripotency (*Pou5f1*), mesoderm specification/expansion (*T, Cdx1, Sp5,* and *Zic3*), anterior-posterior patterning (*Hoxb1* and *Hoxa1*), and paraxial mesoderm development (*Hes7, Meis2, Tbx6, Gbx2,* and *Foxb1*).

CellRank predicted 66 genes to be drivers and 115 genes to be anti-drivers of PM fate (Fig. 5E and data S26). A substantial proportion (22%) of PM drivers were transcription factors. These included *Tcf15* and *Foxc2*, transcription factors that have been previously recognized for their role in regulating PM development (Fig. 5E) (*69, 94*). The *Irx* transcription factors *Irx5, Irx1,* and *Irx3* – known for their roles in segmentation during development (*95*) – were also identified as PM drivers. PM drivers did not exhibit significant over-representation of any Hallmark gene sets (Fig. 5F). Of all PM anti-drivers, 20% were identified as transcription factors, including *Hand1, Cdx2, Msx2, Msx1,* and *Cdx4*. Intriguingly, *Hand1, Msx2, and Msx1* had also been predicted as drivers for FHF. This overlap prompted us to conduct a comparative analysis between PM anti-drivers and FHF drivers. Our examination revealed that 40% of all PM anti-drivers were concurrently FHF drivers (Fig. 5, G and H).

The confluence of expressed genes between FHF drivers and PM anti-drivers seems to indicate that during normal development, increased expression of these shared genes by mesoderm cells predisposes them to transition into FHF, while decreased expression of these same genes steers them towards PM (Fig. 5I). This set of shared genes may thus be considered to be a “lineage branching signature”.

How does *Nipbl*-haploinsufficiency impact the expression of driver and anti-driver genes? Examination of *Nipbl*^+/-^ embryos revealed significant misexpression in NMP, MMP, and PMP cells of numerous driver and anti-driver genes associated with both FHF (first heart field) and PM (paraxial mesoderm) (Fig. 5J, fig. S13, data S27, and data S28). Interestingly, most affected driver and anti-driver genes were downregulated. To ascertain whether this was more than coincidental, we employed a Chi-square analysis, which revealed a striking pattern: *Nipbl^+/-^*embryos underexpressed FHF drivers at a higher frequency than they overexpressed them (166 out of 174 differentially expressed genes) (Fig. 5J). Similarly, they commonly underexpressed PM anti-drivers (40 out of 51) (fig. S13), aligning with expectations since many PM anti-drivers serve dual roles as FHF drivers.

Strikingly, genes associated with EMT were highly enriched among FHF drivers that were underexpressed in *Nipbl^+/-^*embryos (Fig. 5K). A total of 15 EMT genes was found to be underexpressed, including the EMT transcription factors *Msx1* and *Snai2*, and signaling genes such as *Igfbp4*, *Pdgfrb*, and *Tgfb1* (Fig. 5J). These results suggest that, in the context of *Nipbl*-haploinsufficiency, skewed differentiation of mesoderm cells towards the PM lineage at the expense of FHF lineage may be attributable to the underexpression of genes driving FHF fate. These results are consistent with the idea that genes associated with EMT play a role in this lineage misdirection.

### *Nipbl*^+/-^ mice show large changes in the expression of major developmental regulators in all germ layers

Using DGEA to compare the germ layers of LB-stage WT and *Nipbl^+/-^*embryos (data S29, data S30, and data S31), we found that *Nipbl^+/-^*embryos misexpressed hundreds of genes across all germ layers. Underexpression was more common than overexpression in all cases (Fig. 6A), supporting a broad role for *Nipbl* in enhancing gene expression. The majority of gene expression changes were subtle, i.e., less than 2-fold (Fig. 6A). This pattern of gene expression changes agrees with earlier studies (*18*). Gene set enrichment analysis (GSEA) for MSigDB’s Hallmark gene sets found that *Nipbl^+/-^* embryos showed enrichment, and no de-enrichment, for 4 out of 40 gene sets: oxidative phosphorylation in the mesoderm; Mtorc1 signaling in the ectoderm; and G2M checkpoint, Myc targets, and E2F targets in the endoderm (Fig. 6B). In these cases, however, enrichment was driven by a relatively small number of the genes in these sets; the vast majority were either not differentially expressed in *Nipbl^+/-^* embryos or showed changes much smaller than 1.5-fold (Fig. 6C).

**Fig. 6.**
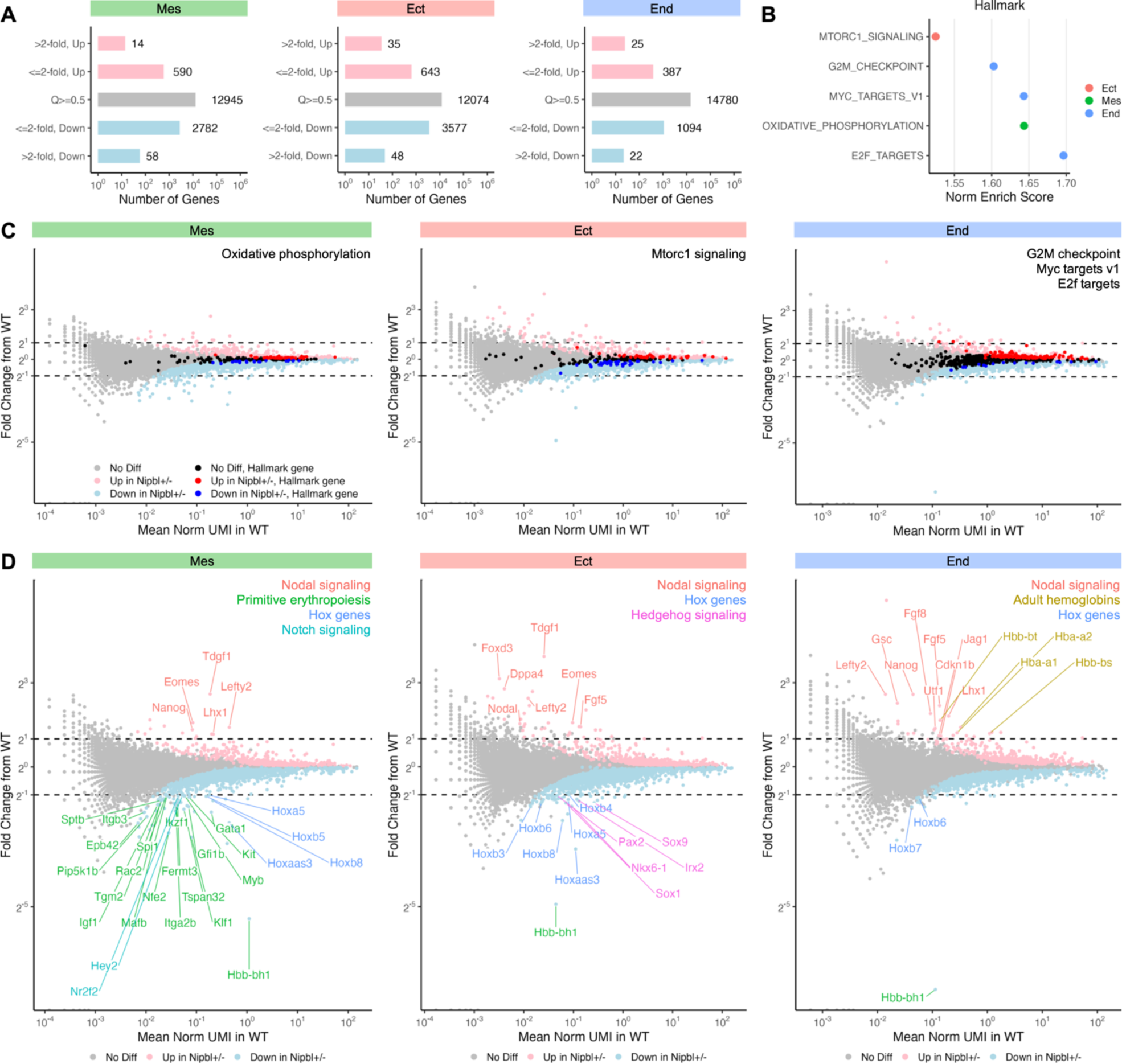
*Nipbl*^+/-^ mice show large changes in the expression of major developmental regulators in all germ layers. (**A**) Number of differentially expressed genes (*Q* < 0.05, Mann-Whitney U Test) in germ layers of LB-stage *Nipbl*^+/-^ embryos showing small (≤ 2-fold) or large (> 2-fold) changes in expression from that of WT embryos. (**B**) Normalized enrichment score of statistically significant (*Q* < 0.05, FGSEA) Hallmark genes sets in germ layers of LB-stage *Nipbl*^+/-^ embryos from that of WT embryos. (**C**) Fold change in expression of Hallmark gene sets from Fig. 5B in germ layers of LB-stage *Nipbl^+/-^* embryos from that of WT embryos along their average expression in WT embryos. (**D**) Fold change in expression of differentially expressed genes (*Q* < 0.05, Mann-Whitney U Test) in germ layers of LB-stage *Nipbl^+/-^* embryos showing large changes in expression (> 2-fold) from that of WT embryos along their average expression in WT embryos.

Genes exhibiting substantial changes in expression – greater than 2-fold upregulated or downregulated – were nevertheless observed across every germ layer of *Nipbl^+/-^* embryos (Fig. 6A). Individual curation of these genes unveiled associations with developmental processes (Fig. 6D). Within the mesoderm, two- to six-fold upregulation of genes involved in Nodal signaling, including *Tdgf1, Eomes, Lefty2, Lhx1*, was observed (*96–99*). At the same time, three clusters of genes were robustly downregulated: those associated with primitive erythropoiesis, including *Gata1* and *Klf1* (2-3 fold) (*74, 100*); *Hox* genes such as *Hoxb8, Hoxb5,* and *Hoxa5* (2-fold); and genes associated with Notch signaling, including *Hey2* and *Nr2f2* (2-fold) (*101, 102*) (Fig. 6D).

In the ectoderm, five transcription factor genes tied to the hedgehog signaling pathway were downregulated more than 2-fold, including *Pax2, Sox9, Nkx6-1, Sox1,* and *Irx2* (*103–107*) (Fig. 6D). In the endoderm, there was a distinct 2-3 fold upregulation of several adult hemoglobin genes such as *Hbb-bt, Hba-a2, Hba-a1,* and *Hbb-bs* (Fig. 6D). Intriguingly, consistent patterns were evident in both the ectoderm and endoderm, mirroring the mesoderm’s trends. Genes related to Nodal signaling were upregulated, and *Hox* genes were noticeably downregulated, across all germ layers. The magnitude of changes in Nodal signaling genes and *Hox* genes, being greater than 2-fold in both directions, and their critical roles in various germ layers, strongly suggest that disruptions in these pathways could be major contributors to phenotypes observed in *Nipbl^+/-^*mice.

When we compared the sets of differentially expressed genes (DEGs) obtained from forward projection (Fig. 6) with those obtained from reverse projection (fig. S14), we found greater than 94% of DEGs in mesoderm and ectoderm were identical (fig. S14, A and B). For the endoderm, reverse projection showed more than 75% of the same DEGs (fig. S14C). As was the case for changes in the relative sizes of cell populations, we conclude that DEGs observed in *Nipbl^+/-^*embryos represent true biological differences and are not a consequence of technical issues in cell classification.

### *Nipbl*^+/-^ mice overexpress *Nanog* during and after gastrulation

To uncover developmental processes that might be the most broadly impacted in *Nipbl^+/-^* embryos, we used the large, significant, changes in gene expression (greater than 2-fold up or down) as input data for STRING (*108*), a database and algorithm that constructs a network of potential gene interactions (data S35), in this case limiting predictions to those informed by experiments demonstrating co-expression or protein-protein interactions. The outcome of this analysis (Figs. 7, A and B) revealed that many genes with substantial expression changes in *Nipbl^+/-^* embryos are predicted to interact either directly or indirectly with *Nanog*. Indeed, *Nanog* itself was overexpressed across all three germ layers in *Nipbl^+/-^* embryos, with a more pronounced overexpression, exceeding 2-fold, in the mesoderm and endoderm (Fig. 7B).

**Fig. 7.**
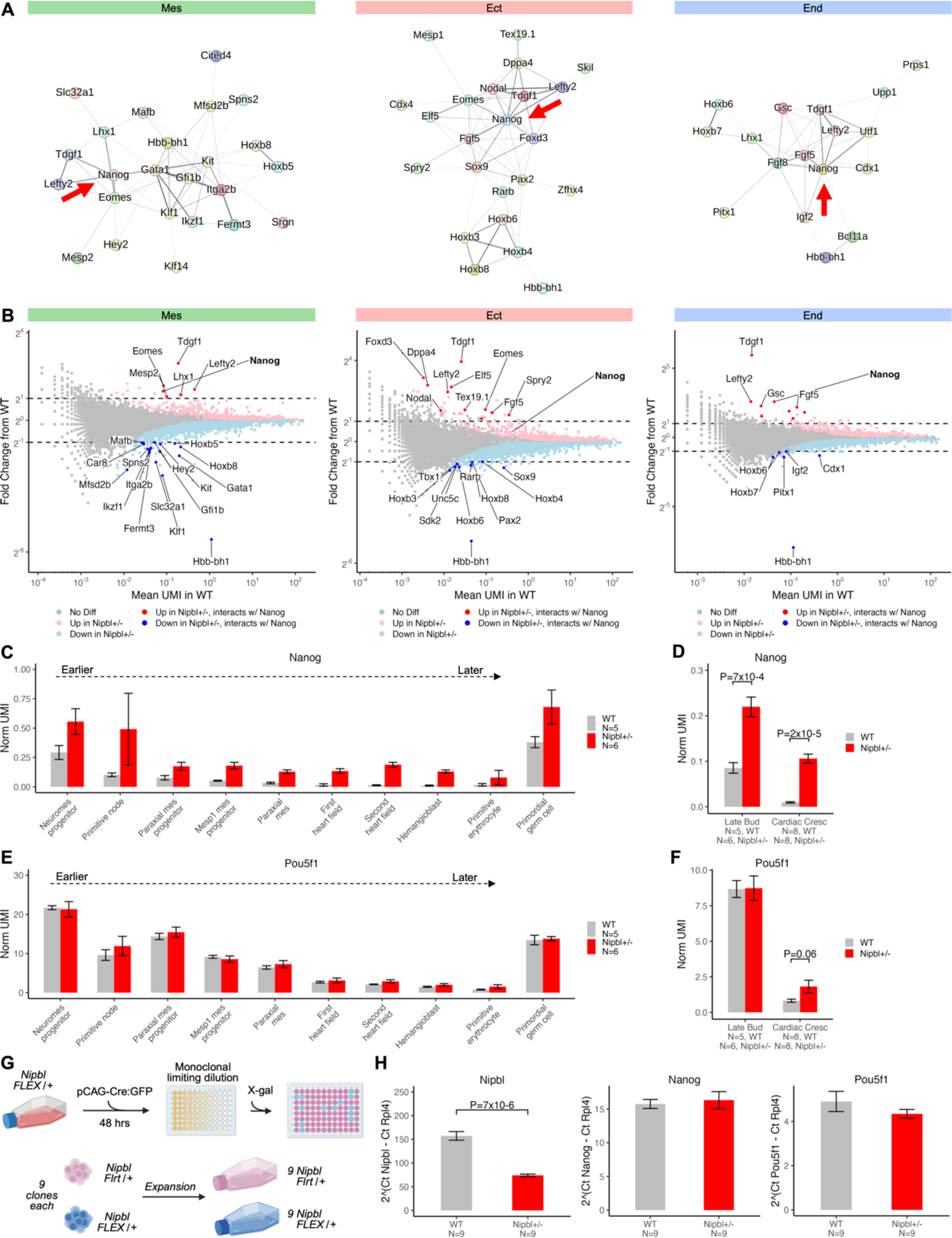
*Nipbl*^+/-^ mice overexpress *Nanog* during and after gastrulation. (**A**) Differentially expressed genes (greater than 2-fold upregulated or downregulated) between the germ layers of WT and *Nipbl*^+/-^ embryos predicted by STRING to interact with each other. (**B**) Fold change in expression of differentially expressed genes (*Q* < 0.05, Mann-Whitney U Test) in germ layers of LB-stage *Nipbl^+/-^*embryos showing large changes in expression (> 2-fold) from that of WT embryos along their average expression in WT embryos that are predicted by STRING to interact with *Nanog*. (**C**) Expression of *Nanog* in mesodermal cell populations of LB-stage WT and *Nipbl^+/-^* embryos ordered from left (earlier) to right (later) by their RNA velocity positions in Fig. 4A. Error bars show SEM. (**D**) Expression of *Nanog* in WT and *Nipbl^+/-^* embryos from LB- to CC-stage. Error bars show SEM. *P*-values from T-Test. € Expression of *Pou5f1* in mesodermal cell populations of LB-stage WT and *Nipbl^+/-^* embryos ordered from left (earlier) to right (later) by their RNA velocity positions in Fig. 4A. Error bars show SEM. (**F**) Expression of *Pou5f1* in WT and *Nipbl^+/-^*embryos from LB- to CC-stage. Error bars show SEM. *P*-values from T-Test. (**G**) Monoclonal generation of *Nipbl*^Flrt/+^ (WT) and *Nipbl*^Flex/+^ (*Nipbl^+/-^*) ESCs. (**H**) Expression of *Nipbl*, *Nanog*, and *Pou5f1* in WT and *Nipbl*^+/-^ embryonic stem cells as measured by RT-qPCR and normalized to the housekeeping gene, *Rpl4*. Error bars show SEM. *P*-values from T-Test.

*Nanog*, which encodes a transcription factor, is notably expressed at two junctures during mouse embryonic development. Initially, it appears at the blastocyst stage, where it plays an essential role in sustaining the pluripotency of the inner cell mass cells (*109*). Subsequently, during gastrulation, *Nanog* expression temporarily surges, only to be silenced as cells transition out of the primitive streak (*110*). This expression pattern could be observed directly in our data: In Fig. 7C, we ordered mesodermal cell populations from LB-stage embryos by their stage in gastrulation, as inferred from their RNA velocity positions (Fig. 4A). In WT NMPs, *Nanog* is expressed at a relatively high level. As cells move through successive stages of gastrulation within WT embryos, *Nanog’s* expression exhibits a consistent decline (Fig. 7C and data S36), reaching near zero in cells of the first and second heart field (FHF and SHF). An exception to this trend occurs in primordial germ cells, which are known to maintain elevated *Nanog* expression (*111*). In contrast, in *Nipbl^+/-^* embryos (Fig. 7C), *Nanog* expression remained elevated across all cell populations. Although expression still falls after the NMP stage in *Nipbl^+/-^* cells, *Nanog* never falls to WT levels. This is evident even in primordial germ cells. Moreover, the overexpression of *Nanog* in *Nipbl^+/-^* embryos is sustained up to the CC-stage (Fig. 7D and table S10), where it reaches a 10-fold increase over WT levels. These observations indicate that *Nipbl*-haploinsufficiency leads to a marked failure of *Nanog* downregulation.

Another transcription factor that plays a role in maintaining pluripotency is *Pou5f1* (Oct4). From studies in embryonic stem cells, it is known that *Pou5f1* promotes the expression of *Nanog*, and conversely, *Nanog* promotes the expression of *Pou5f1* (*112*). Given this, we were curious about whether *Nipbl*^+/-^ embryos might also overexpress of *Pou5f1*. Despite the fact that *Pou5f1* is more highly expressed than *Nanog*, we did not observe any statistically significant difference in *Pou5f1* expression across mesodermal cell types or throughout LB-stage *Nipbl^+/-^* embryos (Fig. 7E, Fig. 7F, data S37, and table S11). However, a more than 2-fold elevation in *Pou5f1* expression was detected in *Nipbl^+/-^* embryos at CC-stage (Fig. 7F).

We sought to determine whether elevated *Nanog* expression in *Nipbl^+/-^* embryos during gastrulation might reflect some sort of non-specific overactivity of the *Nanog* gene. To investigate this hypothesis, we looked at *Nanog* expression in WT and *Nipbl^+/-^* embryonic stem cells (ESCs), as ESCs are known to express *Nanog* (as do to the blastocyst inner cell mass cells from which ESCs are derived). We generated WT and *Nipbl^+/-^* ESCs by treating *Nipbl*^Flex/+^ (*Nipbl^+/-^*) ESCs with Flp recombinase (which inverts a gene trap in *Nipbl*-Flex ESCs in such a way that reverses gene trapping) to produce *Nipbl*^Flrt/+^ (WT) ESCs (Fig. 7G) (*24*). We assessed *Nanog* expression in nine independent clones from each genotype, using quantitative reverse transcription polymerase chain reaction (RT-qPCR). We observed no discernible difference in *Nanog* expression between *Nipbl^+/-^* and WT ESCs, a pattern that was also observed for *Pou5f1* (Fig. 7H and data S38). As a control, we showed that *Nipbl^+/-^* ESCs exhibit the expected reduced expression of *Nipbl*, as also seen in *Nipbl^+/-^* embryos. Collectively, these findings support the conclusion that the overexpression of *Nanog* in *Nipbl^+/-^* LB-stage and CC-stage embryos is result of a specific failure to appropriately suppress *Nanog* following gastrulation, rather than unusually elevated *Nanog* expression persisting from early embryogenesis.

### *Nanog* overexpression accounts for many of the gene expression changes in *Nipbl^+/-^* mice

As a critical regulator of pluripotency in early embryonic development, *Nanog* directly influences the expression of a multitude of other genes. This relationship prompted us to ask how many of the gene expression differences in *Nipbl^+/-^* embryos might be attributable to the overexpression of *Nanog*. To address this question, we took advantage of data from a recent study, by Tiana et al., 2022 (*113*), in which mice were engineered to express *Nanog* under doxycycline-inducible (Dox) control. In that study, bulk RNA sequencing was performed on untreated (*Nanog* Dox-) and Dox-treated (*Nanog* Dox+) embryos at stages E7.5 and E9.5, with E7.5 corresponding closely to the LB-stage embryos analyzed here. The E7.5 *Nanog* Dox+ embryos that were sequenced had been treated with doxycycline from E4.5 to E7.5, while E9.5 *Nanog* Dox+ embryos were treated with doxycycline from E6.5 to E9.5. Differential gene expression analysis, performed in that study, found hundreds of gene expression changes between *Nanog* Dox- and *Nanog* Dox+ embryos at both stages (*113*). We subsequently analyzed the results of their analysis in four ways.

First, we plotted the fold change in expression of genes in *Nipbl^+/-^*embryos, as a whole, from that of WT embryos, concentrating on LB-stage (Fig. 8A data S39). (CC-stage comparisons are shown in fig. S15A, fig. S15B, and data S40.) In Fig. 8A, genes significantly overexpressed in *Nipbl*^+/-^ embryos are colored pink and those significantly underexpressed, light blue. DEGs in *Nipbl^+/-^* embryos that are also significantly overexpressed or underexpressed in E7.5 *Nanog* Dox+ embryos, are colored red and blue, respectively. The results are summarized in Fig. 8B. At LB-stage, 68.8% of overexpressed and 61.5% of underexpressed genes in *Nipbl^+/-^* embryos were also overexpressed or underexpressed, respectively, in E7.5 *Nanog* Dox+ embryos. A binomial test confirmed that *Nanog* Dox+ DEGs are highly over-represented in *Nipbl^+/-^* embryos (Fig. 8B).

**Fig. 8.**
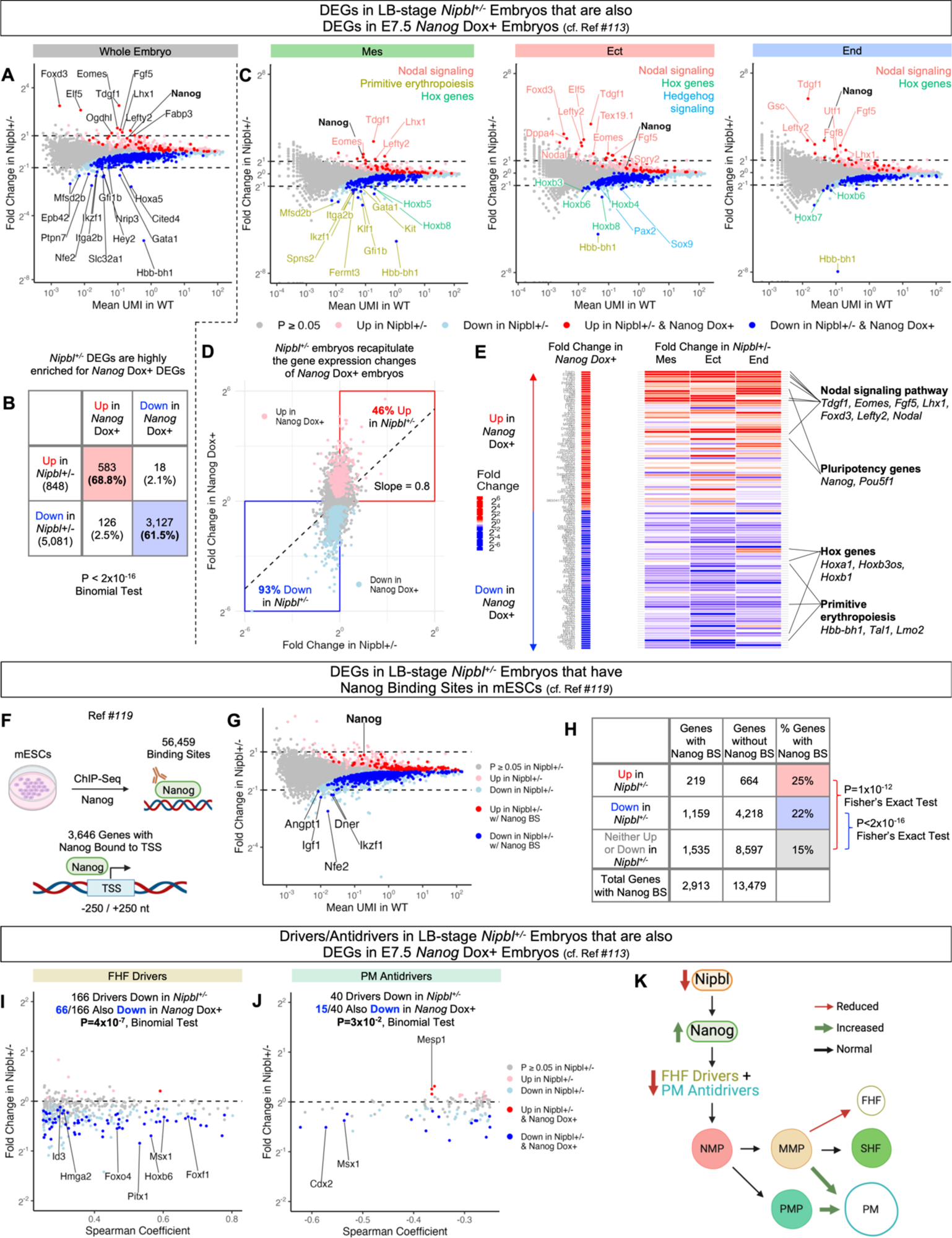
LB-stage *Nipbl^+/-^* mice replicate the gene expression changes of *Nanog* overexpression. (**A**) Fold change in expression of DEGs in LB-stage *Nipbl*^+/-^ embryos (*Q* < 0.05, Mann-Whitney U Test) that are also DEGs in E7.5 *Nanog* Dox+ embryos (*Q* < 0.05, T-Test) (*113*). (**B**) Percentage of DEGs in LB-stage *Nipbl*^+/-^ embryos that are also DEGs in E7.5 *Nanog* Dox+ embryos (*113*). (**C**) Fold change in expression of DEGs in the germ layers of LB-stage *Nipbl*^+/-^ embryos that are also DEGs in E7.5 *Nanog* Dox+ embryos (*113*). (**D**) Fold change in expression of genes in LB-stage *Nipbl*^+/-^ embryos versus their fold change in E7.5 *Nanog* Dox+ embryos. DEGs in E7.5 *Nanog* Dox+ embryos (*113*) are colored pink and light blue. (**E**) Heatmap of the fold change in expression of DEGs from E7.5 *Nanog* Dox+ embryos (lowest *Q*-values from T-Test) (*113*) in the germ layers of LB-stage *Nipbl*^+/-^ embryos. (**F**) ChIP sequencing for Nanog in mESCs from (*119*) identified 3,646 genes with one or more Nanog binding sites within +/- 250 nt of their TSSs. (**G**) Fold change in expression of DEGs in LB-stage *Nipbl*^+/-^ embryos with one or more Nanog binding sites from (*119*). (**H**) Percentage of DEGs in LB-stage *Nipbl*^+/-^ embryos with one or more Nanog binding sites from (*119*). Fold change in expression of DEGs in mesoderm cells of (**I**) FHF and (**J**) PM lineages of LB-stage *Nipbl*^+/-^ embryos that are also DEGs in E7.5 *Nanog* Dox+ embryos (*113*). Transcription factor genes are labeled. (**K**) Reduction of *Nipbl* levels leads to the upregulation of *Nanog* in LB-stage *Nipbl^+/-^*embryos and downregulation of FHF drivers and PM antidrivers, resulting in the misallocation of mesoderm cells to PM at the expense of FHF.

Second, we again plotted the fold change in expression of genes in LB-stage *Nipbl^+/-^* embryos from that of WT embryos, but this time separately by germ layer, rather than across whole embryos (Fig. 8C). Genes colored red and blue are DEGs in *Nipbl^+/-^* embryos that are also overexpressed or underexpressed in *Nanog* Dox+ embryos. When we manually curated genes that were expressed with more than 2-fold change (Fig. 8C), we found that all germ layers showed upregulation of genes involved with Nodal signaling (*Tdgf1, Lefty2, Eomes, Lhx1,* and *Ffg5*) (*96–99, 114*) and downregulation of *Hox* genes (*Hoxb8 and Hoxb6*). Additionally, the mesoderm of *Nipbl^+/-^* embryos underexpressed genes associated with primitive erythropoiesis (*Hbb-bh1*, *Klf1*, and *Gata1*) (*74, 100*) and hedgehog signaling (*Pax2* and *Sox9*) (*103, 104*). (CC-stage gene expression changes are discussed in fig. S15C.) These findings are consistent with the idea that many of the largest changes in gene expression that occur in *Nipbl^+/-^* embryos, as well as the developmental pathways they regulate, could be attributable to overexpression of *Nanog*.

Third, we analyzed how closely *Nipbl^+/-^* embryos mirrored the gene expression alterations of *Nanog* Dox+ embryos in terms of magnitude and direction. To do this, we generated a plot illustrating the fold-changes in expression of individual genes in *Nipbl^+/-^*embryos compared to their fold change in *Nanog* Dox+ embryos (Fig. 8D). We colored genes that were significantly overexpressed in *Nanog* Dox+ embryos pink and those significantly underexpressed, light blue. Additionally, we calculated the regression line depicting the relationship between fold changes in gene expression in *Nipbl^+/-^ vs. Nanog* Dox+ embryos. At LB-stage, the slope of the regression line between *Nipbl^+/-^*and E7.5 *Nanog* Dox+ embryos was 0.8, implying that, on average, the fold changes in gene expression in *Nipbl^+/-^* embryos are quantitatively similar to those in *Nanog* Dox+ embryos (Fig. 8D). Fig. 8D also depicts that, at LB-stage, *Nipbl^+/-^* embryos upregulate 46% of the same genes that were overexpressed in E7.5 *Nanog* Dox+ embryos and downregulate 93% of the same genes that were underexpressed in E7.5 *Nanog* Dox+ embryos. (Similar analysis was performed at CC-stage, and is described in fig. S15D.) These findings highlight a remarkable similarity in gene expression changes between *Nipbl^+/-^* and *Nanog* Dox+ embryos.

Fourth, we examined the magnitudes of fold changes in individual gene expression in *Nipbl^+/-^* embryos and *Nanog* Dox+ embryos, focusing on the 50 most highly up- and down-regulated genes in *Nanog* Dox+ embryos. As shown in Fig. 8E, we categorized genes into two groups at each stage: those overexpressed in *Nanog* Dox+ embryos (ranked from top to middle) and those underexpressed in *Nanog* Dox+ embryos (ranked from bottom to middle), ordering them by ascending Q-value. Next, we constructed a heatmap that visually captures the fold changes in gene expression within *Nanog* Dox+ whole embryos, contrasting them with the corresponding fold changes in the individual germ layers of *Nipbl^+/-^*embryos (Fig. 8E). E7.5 *Nanog* Dox+ embryos overexpressed genes linked with the Nodal signaling pathway (*Tdgf1, Eomes, Fgf5, Lhx1, Foxd3, Lefty2,* and *Nodal*) (*96–99, 114–116*) and those integral to maintaining pluripotency (*Nanog* and *Pou5f1*) (Fig. 8E) (*112*). Concurrently, they underexpressed genes associated with primitive erythropoiesis (*Hbb-bh1, Tal1,* and *Lmo2*) (*117, 118*) and specific *Hox* genes (*Hoxa1, Hoxb3os,* and *Hoxb1*). Strikingly, this pattern of fold changes was mirrored, to a remarkable degree, in the germ layers of LB-stage *Nipbl^+/-^* embryos (Fig. 8E). (The corresponding analysis for CC-stage embryos is given in fig. S15E.)

### Differentially expressed genes from *Nipbl^+/-^* embryos are enriched for Nanog binding sites

Given the large overlap between the gene expression changes observed in LB-stage *Nipbl*^+/-^ embryos and E7.5 *Nanog* Dox+ embryos, we wondered how many differentially expressed genes in *Nipbl^+/-^* embryos might be direct targets of Nanog. Although genome-wide patterns of Nanog binding have not been characterized for embryos at this stage, many groups have used chromatin immunoprecipitation sequencing (ChIP-seq) to define Nanog binding sites in mESCs. We turned to a recent study (*119*), in which Avsec et al. captured 52,456 Nanog binding sites in the genome of mESCs, and reported that 3,645 genes showed one or more Nanog binding sites located within 250 nt on either side of their transcription start sites (Fig. 8F). On comparing these genes with those up- and down-regulated in LB-stage *Nipbl^+/-^* embryos, we found that 25% of the overexpressed genes and 22% of the underexpressed genes match those genes with Nanog binding sites in mESCs (Fig. 8, G and H). This represented significant enrichment for Nanog binding sites in differentially vs. non-differentially expressed genes (P<1×10^-12^ and P<2×10^-16^, Fisher’s Exact Tests). These data suggest that a substantial portion of differentially expressed genes in LB-stage *Nipbl^+/-^* embryos could indeed be direct targets of Nanog.

### *Nanog* overexpression may account for the downregulation of FHF drivers in *Nipbl^+/-^* mice

As shown above in Fig. 3K and fig. S13, FHF drivers and PM antidrivers are downregulated in LB-stage *Nipbl^+/-^* mice. Given the large overlap between downregulated genes in LB-stage *Nipbl^+/-^* embryos and downregulated genes in E7.5 *Nanog* Dox+ embryos (Fig. 8, B and D), we wondered how many of the downregulated FHF drivers and PM antidrivers in *Nipbl^+/-^*embryos could have their changes in expression attributed to the overexpression of *Nanog*. To answer this question, we compared the FHF drivers and PM antidrivers downregulated in LB-stage *Nipbl^+/-^* embryos to those genes exhibiting downregulation in E7.5 *Nanog* Dox+ embryos. We found that approximately 40% of the FHF drivers and 38% of the PM antidrivers that were downregulated in *Nipbl^+/-^* embryos were also downregulated in E7.5 *Nanog* Dox+ embryos (Fig. 8, I and J). This raises the possibility that overexpression of *Nanog* leads to the downregulation of FHF drivers and PM antidrivers, and that this results in the misallocation of mesoderm cells to a PM fate at the expense of FHF (Fig. 3 and Fig. 8K).

### *Nipbl^+/-^* mice exhibit delayed expression of anterior *Hox* genes

We noticed that LB-stage *Nipbl^+/-^* embryos exhibited a significant underexpression, exceeding 2-fold, in *Hox* genes across all germ layers, especially genes within the *Hoxb* cluster (Fig. 6D and Fig. 8C). As *Hox* genes are integral to spatial patterning in development, we compared the expression of all *Hox* genes in *Nipbl^+/-^* embryos with their WT counterparts. The results, shown in Fig. 9, A and B, revealed extensive misexpression at both LB- and CC-stage. At LB-stage, any *Hox* gene that was significantly differentially expressed was underexpressed in *Nipbl^+/-^* embryos in every germ layer where expression was detectable (Fig. 9A). At CC-stage, we observed a trend of underexpression across all germ layers for the majority of misexpressed *Hox* genes (Fig. 9B), but exceptions were found in the mesoderm and ectoderm, where a select few *Hox* genes were overexpressed. These included *Hoxb1* and *Hoxb2* in the mesoderm and *Hoxa1* in the ectoderm, all of which are characterized as anterior *Hox* genes.

**Fig. 9.**
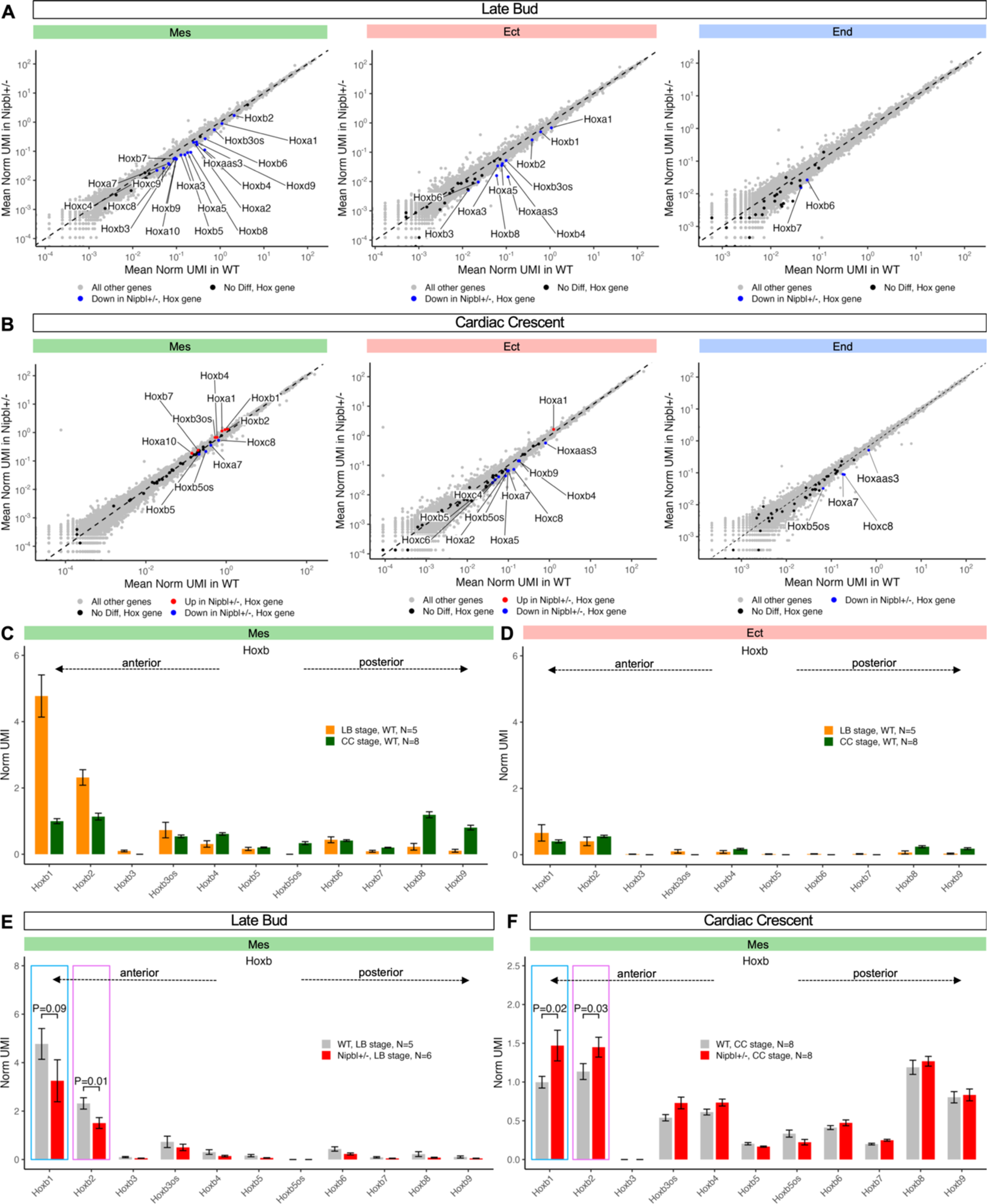
*Nipbl*^+/-^ mice show a delay in the expression of anterior *Hox* genes. Expression of *Hox* genes in germ layers of WT and *Nipbl*^+/-^ embryos at (**A**) LB- and (**B**) CC-stage. Expression of *Hoxb* genes in mesoderm (**C**) and ectoderm (**D**) of WT embryos from LB- to CC-stage ordered from left (5’) to right (3’) by their chromosomal position. Expression of *Hoxb* genes in mesoderm of WT embryos at (**E**) LB- and (**F**) CC-stage ordered from left (5’) to right (3’) by their chromosomal position.

*Hox* genes are organized into chromosomal clusters that turn on expression in a wave-like temporal pattern, producing highly structured and sequential activation during development. Specifically, genes at the 5’ end of the cluster, known as anterior *Hox* genes, are activated early in development, and progressively silenced as genes further forward the 3’ end (posterior *Hox* genes) are turned on (*120*). To investigate whether our data reflected this known pattern, we examined the *Hoxb* genes, as they were the most prominently expressed *Hox* genes at both LB- and CC-stage. After ordering them according to their chromosomal positions, we plotted their expression within the mesoderm and ectoderm of WT embryos at the corresponding developmental stages. In the mesoderm of LB-stage embryos anterior *Hoxb* genes were actively expressed, while posterior *Hoxb* genes were only faintly detectable, if at all (Fig. 9C and data S44). At CC-stage, the expression of anterior *Hoxb* genes decreased from their LB-stage levels, accompanied by the emergence of posterior *Hoxb* gene expression (Fig. 9C). This dynamic shift in expression aligns closely with the established understanding of *Hox* gene regulation during development. In contrast, the ectoderm exhibited a less pronounced pattern, with both anterior and posterior *Hoxb* genes expressing at minimal levels (Fig. 9D and data S45). No discernible difference in the expression of anterior *Hoxb* genes was detected in the ectoderm between LB- and CC-stage embryos (Fig. 9D).

Next we looked at the expression of *Hoxb* genes in the mesoderm of *Nipbl^+/-^* embryos. As in WT embryos, anterior *Hoxb* genes were actively expressed at LB-stage, while posterior *Hoxb* genes were silent (Fig. 9E). However, *Hoxb1* and *Hoxb2* were expressed at levels that were conspicuously lower than in WT (Fig. 9E). At CC-stage, *Nipbl^+/-^* embryos reflected the general trend found in WT embryos, with anterior *Hoxb* genes waning in expression as the posterior *Hoxb* genes began their ascent (Fig. 9F, note axis scale). Interestingly, the two most anterior Hox genes, *Hoxb1* and *Hoxb2* did not decline as much, proportionally, as they did in WT embryos, with the outcome being that, by CC-stage, expression of *Hoxb1* and *Hoxb2* in *Nipbl^+/-^* embryos was actually higher than in WT (Fig. 9F). These results collectively provide evidence for a temporal disruption in the *Hox* gene expression program in *Nipbl^+/-^* embryos. Specifically, *Hox* genes appear to be delayed in initiating expression at the LB-stage and similarly tardy in becoming suppressed at the CC-stage.

### *Nipbl^+/-^* mice show anteriorization of thoracic vertebrae, with left-right asymmetry

A previous study (*121*) showed that a knockout of *Hoxb1-Hoxb9* in mice led to the anteriorization of the axial skeleton, characterized by an increased number of thoracic vertebrae with ribs. Since *Nipbl^+/-^* mice underexpress *Hox* genes and show delayed regulation of anterior *Hox* genes, we were curious if similar anteriorizations could be detected in *Nipbl^+/-^* mice (Fig. 9). To explore this, we analyzed the vertebrae and ribs of 15 WT and 12 littermate *Nipbl^+/-^*embryos at E18.5, using alizarin red and alcian blue stains. In WT embryos, we confirmed the presence of the normal count of 13 ribs (Fig. 10A). In contrast, many *Nipbl^+/-^* embryos displayed 14 thoracic vertebrae, the 14th of which bore a range of rib growths, often displaying left-right asymmetry (Fig. 10A). We classified the growths of these 14^th^ vertebrae based on severity, ranking them from low (ss1) to high (ss5), and documented left-right differences (Fig. 7B). Three out of 7 *Nipbl^+/-^*embryos displayed partial growth of a 14th rib on the left side only (S1-S2). These growths were either cartilage only or disconnected formation of rib nub with cartilage, but never any whole ribs. Four out of 7 *Nipbl^+/-^* embryos showed bilateral growth (S3-S5). Partial growth occurred on both the left and right sides, but complete formation a whole rib only ever occurred, in one instance, on the right side (S5). In total, 58% of E18.5 *Nipbl^+/-^* embryos demonstrated some form of rib growth from a 14th thoracic vertebra (Fig. 10C). These findings suggest that anteriorization of the axial vertebrae does occur in *Nipbl^+/-^* embryos, possibly as a result of delayed activation (and inactivation) of anterior *Hox* gene expression.

**Fig. 10.**
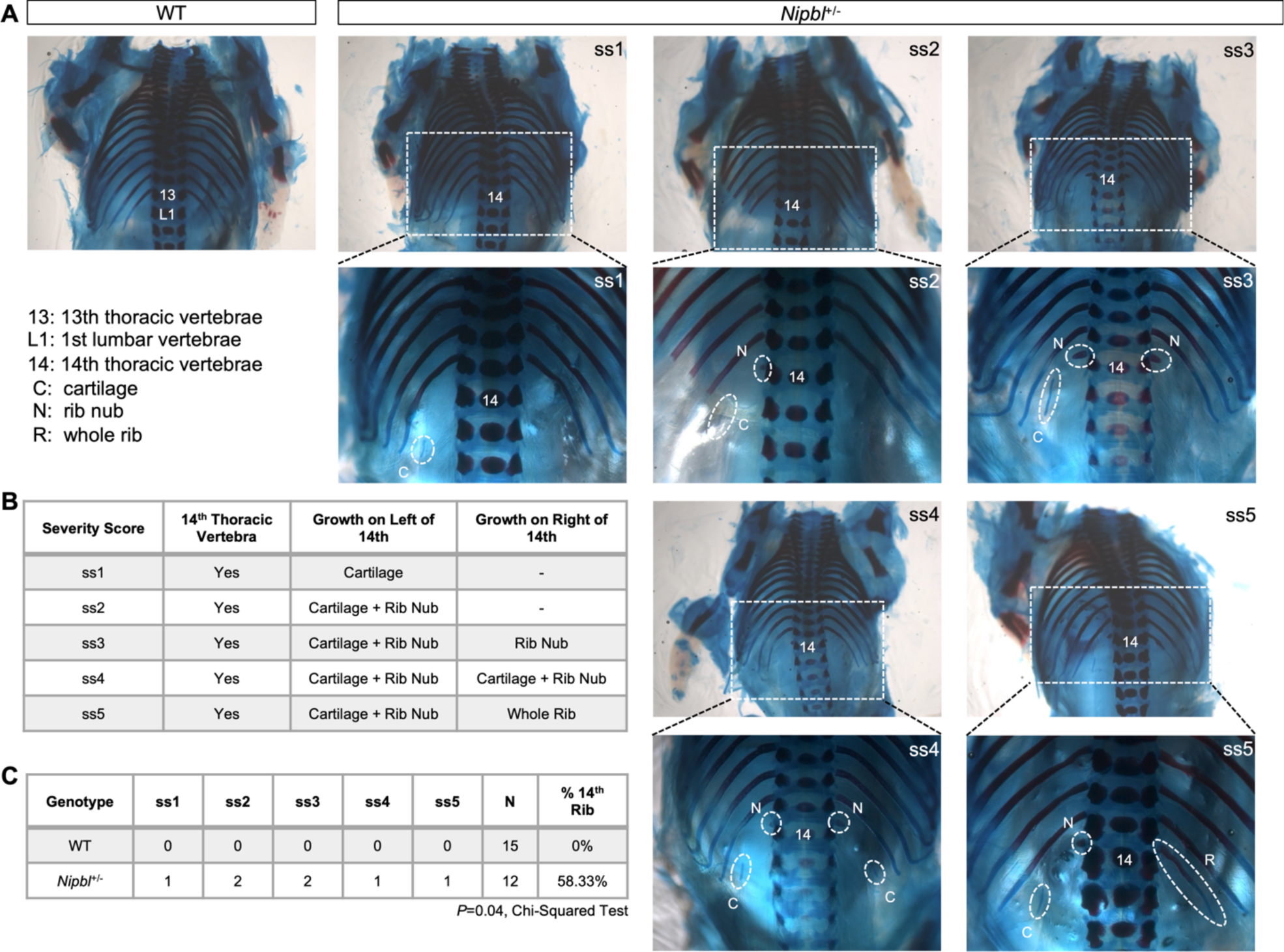
*Nipbl^+/-^* mice show anteriorization of thoracic vertebrae with left-right asymmetry. **(A)** Dorsal view of bone (alizarin red) and cartilage (alcian blue) stained rib cage of E18.5 WT and *Nipbl*^+/-^ embryos. WT embryos only show 13 ribs. *Nipbl*^+/-^ embryos show incomplete asymmetric growth of 14^th^ rib. ss1-ss5 refers to severity score in Fig. 10B. 13 = 13^th^ thoracic vertebrae, L1 = 1^st^ lumbar vertebrae, C = cartilage, N = rib nub, R = whole rib. (**B**) Table categorizing range of incomplete asymmetric growth of 14^th^ rib observed in E18.5 *Nipbl*^+/-^ embryos and ranking them by their severity (low = ss1, high = ss5). (**C**) Table quantifying numbers of WT and *Nipbl*^+/-^ embryos in which incomplete asymmetric growth of 14^th^ rib was observed per severity score. *P*-value from Chi-Squared Test.

## Discussion

The *Nipbl^+/-^* mouse, a model of the multisystem birth defects syndrome, Cornelia de Lange Syndrome (CdLS) (*122*), offers a distinct lens through which to explore the genetic origins of birth defects. *Nipbl^+/-^*mice display many of the same birth defects observed in human CdLS and a key feature of *Nipbl*-deficiency across various organisms is quantitative alterations in gene expression, including both upregulation and downregulation, affecting at least hundreds of genes in every tissue (*18, 24, 123*). Previous research into animal models of *Nipbl*-deficiency suggested that the root causes of birth defects in CdLS likely manifest during a period when progenitor cell populations are being formed for all major tissue and organ lineages (*22, 24*). Therefore, to elucidate how early, cell-type-specific changes in gene expression might contribute to the onset of birth defects, we used scRNAseq to conduct a comparative analysis between *Nipbl^+/-^* embryos and their WT littermates at key developmental stages, focusing on the conclusion of gastrulation, LB-stage (approximately E7.5) to early CC-stage (approximately E7.75) (*25*).

### Changes in the sizes of different mesodermal cell populations in *Nipbl*^+/-^ embryos foreshadow pathological changes in tissue composition and patterning in CdLS

In Fig. 1, we showed that *Nipbl^+/-^* embryos do not lack any cell populations found in WT embryos. However, LB-stage *Nipbl^+/-^* embryos possess fewer total mesoderm cells, and within the mesoderm, have fewer PEs and FHF cells, and more PM cells (Fig. 2). These changes, while not always large, are likely to be of physiological importance. For example, the observation that PEs are reduced in number is consistent with defects in blood formation and circulation observed in *Nipbl*-morphant zebrafish (*22*), and thrombocytopenia in CdLS (*124*).

In addition, the reduction in FHF cell number could contribute to the presence of CHDs in *Nipbl^+/-^* mice and individuals with CdLS (*24, 125*). In a prior study in *Nipbl*^+/-^ mice, the majority of CHDs observed were in heart regions (right ventricle, atrial/ventricular septa) considered to be derivatives of the second heart field (SHF) (*24*). Although FHF cells are not thought to give rise directly to “SHF structures”, they likely exert indirect effects on the development of SHF derivatives. This echoes the conclusions of Santos et al., 2016 (*24*), who found that complex interactions between different cell types (even between non-cardiogenic and cardiogenic cell types), influence the incidence of CHDs in *Nipbl*^+/-^ mice.

The increased number of PM cells at LB-stage is intriguing, as it correlates with observed defects in PM derivatives, including the axial skeleton (Fig. 10) and limb digits in *Nipbl^+/-^*mice (*126*). Notably, both such deficits are features of CdLS (*127*). How an overabundance of PM cells could contribute to limb defects is not obvious. Nevertheless, these findings suggest that birth defects in *Nipbl*-deficient organisms arise not only from the absence of specific progenitor cell populations, but also from progenitor cell misallocation events that alter the relative proportions of cell types.

### Progenitor cell misallocation in *Nipbl^+/-^* correlates with changes in the expression of cell fate driver genes

*Nipbl^+/-^* embryos at LB-stage did not exhibit global changes in apoptotic activity, cell proliferation, developmental timing, or overall lineage trajectories (fig. S11, fig. S12, Fig. 3, and Fig. 4). However, we produced evidence that nascent mesoderm cells in *Nipbl^+/-^* embryos differentiated more frequently into PM than FHF, compared to WT (Fig. 4, C to I). This led us to conclude that misallocation of mesoderm cells in *Nipbl^+/-^* embryos is driven by alterations in early cell fate choices that occur along the main pathways of otherwise unaltered lineage trajectories. These findings suggest that some types of structural birth defects, such as heart defects, arise from quantitative alterations to cell fate determination, rather than a complete disruption of an entire lineage pathway.

In Fig. 5, we presented evidence that reduced differentiation of mesoderm cells into FHF in *Nipbl^+/-^* embryos could be attributed to the underexpression of genes predicted to drive FHF differentiation. Interestingly, underexpressed FHF driver genes show strong enrichment for genes associated with EMT (Fig. 5, J and K). EMT not only occurs during the emergence of mesenchymal mesoderm and FHF cells during gastrulation; it also plays a role in the contribution of later cell lineages (SHF cells, endocardial cells, and neural crest cells) to various cardiac structures. SHF cells undergo EMT as they participate in the elongation of the heart tube and outflow tract (*128*), and endocardial cells undergo EMT as they form structures such as the cardiac cushion (*129*). Later in development, migratory neural crest cells, which arise from neural epithelium by EMT, contribute to the aorticopulmonary septum and parts of the outflow tract (*130*). Taken together, these observations suggest that dysregulation of EMT genes may play an important role in the development of heart defects in *Nipbl^+/-^* mice.

### Is *Nanog* overexpression responsible for gene expression changes in *Nipbl*^+/-^ embryos?

In contrast to our prior studies of *Nipbl^+/-^* mice, in which differences in gene expression were invariably found to be small (less than 2-fold) (*18*), the present study demonstrated that, in gastrulation stage embryos, some genes show much larger shifts in expression (as high as eight-fold) (Figs. 6D and 8H). One of the largest of such changes was in *Nanog,* which displayed overexpression in LB-stage *Nipbl^+/-^* embryos, and which persisted to CC-stage (Figs. 7 and 8). Using data from Tiana et al., 2022 (*113*), we demonstrated that many of the gene expression changes in *Nipbl^+/-^* embryos mirror those seen in embryos with induced *Nanog* overexpression (Fig. 8, A to E, and fig. S15, A to E). Moreover, many *Nipbl^+/-^* DEGs are likely to be direct Nanog targets, as the corresponding genes display enrichment for Nanog binding sites (Fig. 8, F and H). Comparisons of our data with those of Tiana et al., 2022 (*113*) also suggest that misallocation of mesoderm cells to a PM fate at the expense of FHF, resulting from downregulation of FHF drivers / PM antidrivers, may be a consequence of *Nanog* overexpression (Fig. 3 and Fig. 8K).

A question that arises from these observations is whether all gene expression alterations observed in *Nipbl*^+/-^ embryos might be due to *Nanog* overexpression. For a number of reasons, we do not think that this is the case. First, for the largest gene expression shifts (greater than two-fold over- or underexpressed), only 39% at LB-stage and 18% at CC-stage were misexpressed in the same direction as DEGs in embryos in which *Nanog* was overexpressed (Fig. 8 and fig. S15) (*113*). Second, although *Nanog* continues to be overexpressed in *Nipbl^+/-^* embryos from LB- to CC-stage, the magnitude of its expression decreases substantially by CC-stage (Fig. 7). Third, *Nipbl*-haploinsufficiency itself causes significant, large expression changes in other genes not implicated as consequences of *Nanog* overexpression. For instance, we observed reduced expression of genes linked to the Notch signaling pathway in the mesoderm, and increased expression of adult hemoglobin genes in the endoderm (Fig. 6). Therefore, we conclude that the extensive gene expression changes in *Nipbl^+/-^* embryos are unlikely to be due to *Nanog* overexpression alone.

### Skeletal anomalies in *Nipbl^+/-^* mice reflect earlier delays in anterior *Hox* gene expression

In Figs. 4D and 8G, we showed that large decreases in *Hox* gene expression occur across all germ layers of LB-stage *Nipbl^+/-^* embryos. While only a subset of *Hox* genes is underexpressed by more than two-fold, many *Hox* genes across all germ layers show small (less than two-fold) but significant decreases in expression (Fig. 9A). We also provide evidence for a temporal disturbance in anterior *Hox* gene expression in *Nipbl^+/-^*embryos, whereby *Nipbl*-haploinsufficiency postpones the onset of anterior *Hox* gene expression at LB-stage and delays their subsequent deactivation at CC-stage (Fig. 9, C to F). These shifts in *Hox* gene expression dynamics mirror those in a previous study of *nipbl*-morphant zebrafish, in which genome-location-specific misexpression of *hox* genes was also observed (*131*). In Fig. 10, we showed that anteriorization of thoracic vertebrae occurs in E18.5 *Nipbl^+/-^* mice (Fig. 9). Such alterations in axial skeletal development parallel skeletal anomalies that have been identified in individuals with CdLS, including fused, absent, or misshapen ribs (*132*). Since *Nipbl*-deficiency is the dominant form of CdLS, it is interesting to speculate that these skeletal anomalies might originate from underexpression of and delayed anterior *Hox* gene activity during gastrulation.

### Overexpression of Nodal signaling pathway genes may contribute to left-right patterning defects in CdLS

In Fig. 4D, we showed that *Nipbl^+/-^* embryos at LB-stage show large overexpression of genes associated with the Nodal signaling pathway across all germ layers. Nodal signaling, pivotal for left-right patterning (*133*), ensures the appropriate positioning and morphogenesis of the musculoskeletal system and internal organs (*134*). In Fig. 10, we showed that growth of a 14^th^ vertebra in E18.5 *Nipbl^+/-^*embryos displayed left-right asymmetry, with the growth of a whole 14^th^ rib only occurring on the right side of the embryo. Such patterning alterations are consistent with the heightened incidence of right-sided skeletal anomalies in CdLS (*127*). Changes in left-right patterning are also consistently associated with intestinal malrotation and some types of cardiac anomalies observed in *Nipbl^+/-^* mice, *nipbl*-morphant zebrafish, and CdLS (*18, 24, 135*). These data suggest that left-right patterning defects in CdLS might be due to overexpression of Nodal signaling genes as early as gastrulation.

### Gene expression changes observed in early *Nipbl^+/-^* embryos are likely driven by Nipbl’s effects on chromatin topology

The results of this study, together with prior studies of mouse and zebrafish models, reinforce the notion that the chromosomal location of genes is an important factor in their sensitivity to variations in *Nipbl* levels. For example, our prior studies of *Nipbl^+/-^* embryonic brain demonstrated that the largest expression changes among genes in the 22-gene protocadherin beta (*Pcdhb*) cluster were found in those genes situated at the 5’ and 3’ ends of the cluster (*18*), where CTCF sites are located (*136*). In developing pectoral fins of *nipbl*-morphant zebrafish, *hox* genes located near the 3’ end of three separate *hox* clusters (*hoxa*, *hoxc*, and *hoxd*) show a position-dependent pattern of overexpression (*131*). Similarly, in the present study, we showed that anterior *Hoxb* genes – located at the 3’ end of the *Hoxb* cluster – are preferentially underexpressed in the mesoderm of *Nipbl^+/-^* embryos at gastrulation (Fig. 8E). Thus, we consistently observe a strong influence of chromosomal location on gene sensitivity to *Nipbl* levels, particularly within gene clusters, and this influence is already apparent at gastrulation.

Recent studies suggest that changes in gene expression due to *Nipbl*-deficiency may be the result of global alterations to chromosomal structure and organization. For instance, drastic reduction of *Nipbl* expression in mouse hepatocytes resulted in genome-wide depletion of topologically associated domains (TADs) and Hi-C interaction peaks (*137*), both of which require chromatin looping (*138*). In fact, recent studies have demonstrated that both the formation and rate of chromatin loop extrusion were reduced *in vitro* when a CdLS pathogenic mutation was introduced into *NIPBL* (*139*).

Chromatin looping within TADs is thought to bring distant cis-regulatory elements, such as enhancers, into proximity with target promoters. Interestingly, Nipbl has been reported to preferentially bind to enhancers and promoters (*140*), and ChiP-seq studies show reduced enhancer-promoter interactions in *Nipbl*^+/-^ mouse embryonic fibroblasts (*141*). Since chromatin looping is essential for gene activation, impaired loop extrusion as a consequence of reduced Nipbl levels may provide an explanation for why *Nipbl*-haploinsufficiency causes more genes to be downregulated than upregulated in gastrula-stage mouse embryos (e.g., >80% of mesodermal genes, Fig. 6A), a trend that has been observed in studies of other tissues (*141*).

In light of such observations, it is interesting that overexpression of *Nanog* and its target genes emerged from the present study as particularly likely to play important roles in causing CdLS phenotypes. During normal gastrulation, *Nanog* expression falls to very low levels in most cell types, so what occurs in *Nipbl*^+/-^ embryos is perhaps best viewed as a failure of gene repression. Although it is possible that *Nipbl*^+/-^ haploinsufficiency leads to downregulation of a gene encoding a *Nanog* repressor, no obvious candidate stands out within the gene expression changes identified in the present study. We speculate, instead, that *Nipbl*-sensitive loop formation is required for the silencing of *Nanog* gene expression during gastrulation, and thus that changes in chromatin architecture play a key role in timing the termination of this critical embryonic event.

## Materials and Methods

### Statement on Care and Use of Animals

All animals were handled in accordance with approved procedures as defined by the National Institutes of Health, and all animal work was approved by the Institutional Animal Care and Use Committee of the University of California, Irvine. For collection of mouse tissues, pregnant dams were humanely killed by CO_2_ anesthesia followed by cervical dislocation.

### Generation of WT and *Nipbl*^+/-^ Mice

WT and *Nipbl*^+/-^ mouse littermates were generated by mating *Nanog*^Cre/+^ mice (*26*) and *Nipbl*^Flox/Flox^ mice (*24*). *Nipbl*^Flox/Flox^ mice have an inverted gene trap cassette encoding *β-geo* that is flanked by *Cre* recombinase target sites in intron 1 of *Nipbl* alleles (Fig. 1A) (*24*). In this inverted orientation, referred herein as Flox, there is no trapping of the *Nipbl* gene and *Nipbl* is expressed normally. However, when this cassette is exposed to Cre recombinase, the gene trap cassette gets inverted into a non-inverted orientation which we call FIN (Fig. 1A). In this non-inverted orientation, trapping of the *Nipbl* gene occurs and *β-geo* is expressed as a reporter of successful gene trapping. Therefore, the *Nipbl* FIN allele is a null allele. *Nanog*^Cre/+^ mice carry a transgene encoding a *Cre* recombinase downstream of a promoter of the *Nanog* gene and initiate recombination in the earliest cells of the embryo (*26*). Consequently, mating *Nanog*^Cre/+^ mice with *Nipbl*^Flox/Flox^ mice results in littermates that are either *Nipbl*^Flox/+^ or *Nipbl*^FIN/+^, entirely. *Nipbl*^Flox/+^ mice express *Nipbl* at wildtype levels (Fig. 1A) and show no defects, making them essentially WT (*24*). *Nipbl*^FIN/+^ mice express *Nipbl* at levels ∼50% lower than WT (fig. S3), and show defects similar to those observed in CdLS, making them essentially *Nipbl*^+/-^ (*24*).

### Timing of Mouse Pregnancies

To generate female mice that were pregnant on the same day, male mice were singly housed and female mice were group-housed in groups of five for a minimum of one week to synchronize their estrous cycles, thus taking advantage of the Lee-Boot effect (*142*). At the beginning of the night cycle, the bedding from female cages were discarded and the bedding from at least two male cages were transferred into each of them. At the beginning of the third night cycle after which the females were exposed to male bedding, all females were transferred into male cages, resulting in two females per male, thus taking advantage of the Whitten effect (*143*). At the end of the third night cycle, the females were inspected for vaginal plugs. Those that had vaginal plugs were considered potentially pregnant. To time the dissection of embryos from potentially pregnant females, we considered the end of the 12-hour night cycle after which the vaginal plug was discovered as E0 and dissected the embryos at the following times post E0: E7.5 (7 days + 12 hours) for late bud stage, E7.75 (7 days + 18 hours) for cardiac crescent stage, E7.41 (7 days + 10 hours) for early bud stage, E7.58 (7 days + 14 hours) for early head fold stage embryos, and at E18.5 (18 days + 12 hours).

### Dissection of Mouse Embryos

Pregnant female mice were euthanized by CO_2_ inhalation followed by cervical dislocation. The uterine horns were dissected out of their abdomens with dissection forceps and scissors and placed in a petri dish with 1X DEPC PBS on ice. Individual deciduae were separated from one another and transferred into their own petri dishes with 1X DEPC PBS on ice. Embryos were dissected out of each deciduae under a dissection microscope as described in (*144*). The Reichert’s membrane was removed from each embryo. The ectoplacental cone was separated from embryos collected for scRNAseq, and transferred by forcep into microcentrifuge tubes, where they were kept on ice or stored at 20°C as tissue for polymerase chain reaction (PCR) genotyping. The exocoelom was separated from embryos collected for scRNAseq and transferred into a welled plate by wide bore pipette tip with fixative on ice as tissue for genotyping by X-gal stain. Embryos for scRNAseq were transferred by wide bore pipette tip with 1X DEPC PBS into microcentrifuge tubes and kept on ice until dissociation. Tails were separated from E18.5 embryos collected for Alcian Blue – Alizarin Red staining. Embryos for Alcian Blue – Alizarin Red staining were transferred by forcep into 10% neutral buffered formalin in scintillation vials and kept at 4°C for 24 hours.

### Genotyping of Mouse Embryos and ES Cells

#### X-gal Stain

Since *Nipbl*^FIN/+^ (*Nipbl*^+/-^) mouse embryos express *β-geo* (beta-galactosidase), their tissues will turn blue when they are treated with X-gal, a substrate that releases a blue chromophore when enzymatically acted on by beta-galactosidase. The exocoeloms from embryos were transferred into welled plates containing fixative (0.2% Glutaraldehyde, 5mM EGTA, and 2mM MgCl2 in 0.1M phosphate buffer, pH 7.5) and kept on ice for a minimum of 15 min. After fixation, the exocoeloms were rinsed with a detergent rinse (0.02% Igepal, 0.01% Sodium Deoxycholate, and 2mM MgCl2 in 0.1M phosphate buffer, pH 7.5) before they were treated with X-gal stain (0.02% Igepal, 0.01% Sodium Deoxycholate, 5mM Potassium Ferricyanide, 5mM Potassium Ferrocyanide, and 2mM MgCl2 in 0.1M phosphate buffer, pH 7.5). The exocoeloms were incubated in X-gal stain at 37°C in the dark for a minimum of 1 hour after which they were inspected for coloration under a dissection microscope. Those that turned blue were considered *Nipbl^+/-^* (*Nipbl*^FIN/+^) and those that did not were considered WT.

#### Polymerase Chain Reaction

Ectoplacental cones were treated with 50 ul 60 ug/ml proteinase K in PBND (50 mM KCl, 10 mM Tris-HCl, 2.5 mM MgCl2, 0.1 mg/ml gelatin, 0.45% Nonident P40, and 0.45% Tween 20), for 1 hour at 55℃ to extract DNA for PCR. To deactivate proteinase K so that it does not interfere with DNA amplification, they were then incubated at 95℃ for 10 min. To genotype mouse embryos as either *Nipbl*^Flox/+^ (WT) or *Nipbl*^FIN/+^ (*Nipbl*^+/-^) and ES cells as either *Nipbl*^Flrt/+^ (WT) or *Nipbl*^FLEX/+^ (*Nipbl*^+/-^), standard Taq PCR was performed on the extracted DNA using the primers and thermocyling protocol previously described in (*24*). Primer 1: 5’-CTCCGC CTCCTCTTCCTCCATC-3’. Primer 2: 5’-CCTCCCCCGTGCCTTCCTTGAC-3’. Primer 3: 5’-TTTGAGGGGACGACGACAGTCT-3’. Thermocycling conditions: 1 cycle a 95°C for 30 sec, 30 cycles of 95°C for 1 min, 59°C for 30 sec, and 68°C for 1 min, 1 cycle at 68°C for 5 min, and hold at 4°C. PCR products were treated with loading dye and electrophoresed in agarose gel stained with SYBER Safe DNA Gel Stain (Invitrogen: S33102) without cleanup and visualized under UV light. Flox conformation: 782 base pairs (bp). FIN conformation 518 bp. Flrt conformation: 735 bp. FLEX conformation: 652 bp.

### Single Cell RNA Sequencing

Embryos were transferred by wide bore pipette tip in 20 ul 1X DEPC PBS into microcentrifuge tubes. 200 ul of 1X TrypLE Express Enzyme with phenol red (Gibco: 12605010) prewarmed to 37°C was added to each embryo. Embryos were triturated 4x with a wide bore pipette tip. Embryos were incubated at 37°C and triturated 4x with a wide bore pipette tip every minute (4-8 min) until no tissue aggregates were visible under a dissection microscope. TrypLE Expres Enzyme activity was inactivated with the addition of pre-ice-chilled 200 ul 0.04% (m/v) non-acetylated BSA in DPBS (Sigma-Aldrich: B6917-100MG, Gibco: 14190144). The resulting cell suspension was underlayed with 200 ul of 1% non-acetylated BSA in HBSS (Sigma: 55021C-1000ML) using a gel loading pipette tip. The suspension was centrifuged in a swing bucket centrifuge at 300 rcf for 5 min at 4°C. 600 ul of supernatant was removed without disturbing the cell pellet. 600 ul of HBSS was added to the cell pellet and the cell pellet was resuspended in it using a wide bore pipette tip. The resulting cell suspension was centrifuged in a swing bucket centrifuge at 300 rcf for 5 min at 4°C. 600 ul of supernatant was removed without disturbing the cell pellet. The cell pellet was resuspending in the remaining 20 ul of supernatant using a wide bore pipette tip.

20 ul single cell suspensions were submitted to the Genomics High Throughput Facility (GHTF) at the University of California, Irvine (*145*) for scRNAseq using 10X Genomics’ Chromium Next GEM Single Cell 3’ Kit v3.1 (10x Genomics: 1000268), Chromium Next GEM Chip G Single Cell Kit (10x Genomics: 1000120), and Chromium Controller. Passage through the Chromium Controller resulted in sample, cell, and transcript barcoded cDNA, which GHTF amplified by PCR. GHTF assessed the quality and quantity of the amplified cDNA by electrophoresis using Agilent’s Agilent High Sensitivity DNA Kit (Agilent: 5067-4626) and Bioanalyzer before library construction. Constructed libraries, representing embryonic samples, were multiplexed and sequenced by GHTF on the Illumina HiSeq 4000 to a minimum depth of 20 million read pairs per cell. GHTF demultiplexed Illumina’s raw base call (BCL) files and returned FASTQ files as deliverables.

### Read Mapping and Cell Calling

Cell Ranger v3.0 was used to map reads onto the GRCm38/mm10 C57BL/6J Mus musculus genome/transcriptome assembly and call cells. Cell Ranger does this by using a read mapper called STAR (*146*), which performs splicing-aware mapping of reads to the genome. Reads are considered confidentially mapped to the genome with a MAPQ of 255. Exonic reads are further mapped to annotated transcripts. A read that is compatible with the exons of an annotated transcript, and aligned to the same strand, is considered mapped to the transcriptome. CellRanger called cells using the EmptyDrops method described in (*147*).

### Normalization of Library Depth

Cell Ranger v3.0 was used to normalize the read depth between libraries of the same stage. Cell Ranger does this by subsampling reads from higher-depth libraries until all libraries of the same stage had an equal number of reads.

### Removal of Low-Quality Cells and Doublets

Cells exceeding three median absolute deviations in any one of the following criteria among cells of the same stage were considered either low-quality cells or doublets (*148*) and removed: 1) percentage of mitochondrial genes expressed, 2) number of genes expressed, and/or 3) number of transcripts detected (fig. S2 and fig. S8).

### Normalization of Cell Depth

Seurat v3.0 was used to normalize the read depth between cells of the same stage using the SCtransform method, which is described in (*149*). SCtransform does this by modeling the read counts in a regularized negative binomial model to determine the variation due to read depth and then adjusting that variance according to genes of similar abundances. SCtransform was also used to normalize the read depth between cells when cells were subset from the whole embryo into germ layers and clusters.

### Batch Effect Correction

Seurat v3.0 was used to correct for batch effects among libraries of the same stage and genotype. Seurat does this by identifying a set of shared variable genes among the libraries being considered, and using these genes, identifies pairs of cells between any two libraries whose expression of these genes are similar to other. These pairs of cells act as anchors between libraries for batch effect correction and integration (*30*).

### Clustering of WT Cells

Seurat v3.0 was used to cluster WT cells. Seurat does this by first performing principal component analysis on the shared variable genes identified during batch effect correction and integration. Principal components whose explained variances exceeded 2 median absolute deviations were used to calculate k-nearest neighbors and construct a shared nearest-neighbor graph. Clusters were determined by optimization of the modularity function using the Louvain algorithm. The number of clusters was controlled by modulating the resolution function.

The optimal number of clusters was determined by clustering cells at increasing consecutive numbers of clusters and generating a clustering tree (fig. S5) visualizing how cell cluster identities change as the number of clusters consecutively increase. Clusters are stable when a large proportion of cells are derived from a single preceding cluster rather than multiple preceding clusters. We adopted Shannon Entropy as a measure of these proportions as a measure of intra-cluster stability. A low Shannon Entropy represents high intra-cluster stability. We visually inspected the clustering tree to determine which number of clusters maximized the number of clusters while at the same time minimizing the total Shannon Entropy across all clusters at that number of clusters.

We further performed differential gene expression analysis between clusters and visualized the expression of the top differentially expressed genes in a heatmap. We visually inspected the heatmap to confirm that the number of clusters that was selected for intra-cluster stability also displayed inter-cluster differences in gene expression.

### Differential Gene Expression Analysis Between Clusters

Seurat v3.0 was used to perform differential gene expression analysis between clusters. For each cluster, Seurat performs the Mann-Whitney U Test (a non-parametric test) between cells in that cluster and all other cells using normalized read counts. P-values were corrected for false discovery using the Bonferroni correction method. Genes with Q-values less than 0.05 were considered statistically significant and differentially expressed.

### Projection of *Nipbl*^+/-^ Cells onto WT Clusters

Seurat v3.0 was used to project *Nipbl*^+/-^ cells onto WT clusters. Seurat does this by identifying a set of shared variable genes among WT samples, and using these genes, identifies pairs of cells between WT cells and *Nipbl*^+/-^ cells whose expression of these genes are similar to other. These pairs of cells act as anchors between WT cells and *Nipbl*^+/-^ cells for projecting and sorting *Nipbl*^+/-^ cells into WT clusters (*30*).

### Reverse Projection

Using the same method that was used to cluster WT cells (see Clustering of WT Cells), cells from LB-stage *Nipbl^+/-^* embryos were first clustered and annotated (fig. S6A). Cells from LB-stage WT embryos were then projected onto the *Nipbl^+/-^* clusters (fig. S6, B to D) (see Projection of *Nipbl*^+/-^ Cells onto WT Clusters).

### Differential Gene Expression Analysis Between Genotypes

Seurat v3.0 was used to perform differential gene expression analysis between WT and *Nipbl*^+/-^ cells. For each cluster, Seurat performs the Mann-Whitney U Test between WT and *Nipbl*^+/-^ cells using normalized read counts. P-values were corrected for false discovery using the Bonferroni correction method. Genes with Q-values less than 0.05 were considered statistically significant and differentially expressed.

### Pseudotime

URD was used to calculate the pseudotime of cells from EB-, LB-, and EHF-stage embryos (*83*). URD does this by constructing a diffusion map of transition probabilities and starting with an assigned group of root cells, performs a probabilistic breadth-first graph search using the transition probabilities. This moves step-wise outward from the root cells, until the entire graph is visited. Several simulations are run, and then pseudotime is calculated as the average iteration that visited each cell.

### Construction of Lineage Trajectories

Velocyto (*150*) was used to count the numbers of spliced and unspliced transcripts per gene, using default parameters. Reads aligning to exonic regions were counted as spliced. Reads aligning to intronic regions were counted as unspliced. Reads aligning to exon-intron boundaries were considered ambiguous and excluded from downstream analyses. scVelo (*84*) was used estimate RNA velocities. Counts were normalized using the pp.filter_and_normalize() function, moments of unspliced vs spliced abundances were computed using the pp.moments() function, and velocities were computed using the tl.velocity() function, all using default parameters. Lineage trajectories were visually inferred from stream plots of computed RNA velocities.

### Calculating Fate Probabilities

CellRank was used to calculate the fate probabilities of mesoderm cells (*86*). CellRank does this by performing RNA velocity-directed random walks from initial cell states to terminal cell states. Fate probabilities correspond to the fraction of walks in which a cell was a part of that that terminated in a particular terminal cell state. Terminal states were set with the *set_terminal_states()* function, absorption probabilities were computed with the *compute_absorption_probabilities()* function, and driver genes were computed with the *compute_lineage_drivers()* function, all with default parameters.

### Cell Cycle Phase Assignment

Seurat was used to assign cells into G1, S, or G2/M phase based on the expression of markers of S phase and G2/M phase provided by Seurat. Using the CellCycleScoring() function, Seurat calculated scores for the expression of S phase and G2/M phase markers, while considering the expression of these marker genes to be anticorrelated to one another. When cells express neither, they are considered to be in G1 phase.

### Identification of Drivers and Anti-drivers

CellRank was used to identify the drivers and anti-drivers of FHF and PM fates. CellRank does this by calculating a correlation coefficient between the fate probabilities of cells towards their lineage fate and the expression of their genes. Those genes with positive correlation coefficients are considered drivers, since their expression increases as absorption probabilities increase and those with negative correlation coefficients are considered anti-drivers, since their expression decreases as absorption probabilities decrease. To reduce the likelihood of false discovery, we considered those genes with correlation coefficients greater than 0.25 as drivers and those less than -0.25 as anti-drivers.

### Gene Set Over-Representation Analysis

clusterProfiler (*151*) was used to perform gene set over-representation analysis. It does this by performing a Fisher’s Exact Test on a contingency table of the genes in a gene set that match or do not match the genes of interest. P-values were corrected for false discovery using the Bonferroni correction method. Gene sets with Q-values less than 0.05 were considered statistically significant and over represented by the genes of interest.

### Gene Set Enrichment Analysis

FGSEA (*152*) was used to perform gene set enrichment analysis.

### Generation of Nipbl^FLEX/+^ and Nipbl^Flrt/+^ ESCs

ES cells were grown in Glasgow’s MEM, 15% heat-inactivated FBS (ES cell qualified, Hyclone SH30071.03E), 1X Glutamine, 1X Pen-Strep, 5mM mercaptoethanol, and 1000 U/ml leukemia inhibitory factor (LIF: ESGRO Millipore). Following limited dilution single cell cloning, a clone of EUC313f02 *Nipbl^FLEX/+^* ES cells (European Conditional Mouse Mutagenesis Program) was transfected with pCAG-Cre:GFP (Addgene #13776) using Lipofectamine 2000 (Invitrogen) to convert the *Nipbl^FLEX^* allele to the *Nipbl^Flrt^* conformation in vitro. 48 hours post-transfection, cells were plated at 1 cell/well into several 96 well plates. Clonal colonies were isolated, and the clones were stained for β-galactosidase (lacZ) activity using X-gal. For LacZ staining, the ES cells were fixed for 5 minutes in 2 mM MgCl2, 0.5% glutaraldehyde in 1X PBS, followed by 3 washes with 1xPBS at room temperature. X-gal staining (5 mM K3Fe(CN)6, 5 mM K4Fe(CN)6, 2 mM MgCl2, 1 mg/mL X-gal in 1X PBS) was performed at 37°C until the blue precipitate was detected. Colonies positive for X-gal staining, *Nipbl^FLEX/+^* ES cells, and negative for X-gal staining, *Nipbl^Flrt/+^,* were verified by PCR genotyping, described above.

### Reverse Transcription-Quantitative Polymerase Chain Reaction

Clones for both *Nipbl^FLEX/+^* and *Nipbl^Flrt/+^* were expanded, 9 each, and RNA was extracted using the Monarch Total RNA Miniprep Kit (New England Biolabs). cDNA was made using iSCRIPT reverse transcriptase (BioRad). qRT-PCR for *Nipbl*, *Nanog*, and *Pouf5f1* was performed using iTaq SYBR green (BioRad) as per manufacturer’s instructions, *Rpl4* was used as the housekeeping gene. *Rpl4* primer 1: 5’-ATCTGGACGGAGAGTGCTTT-3’. *Rpl4* primer 2: 5’-GGTCGGTGTTCATCATCTTG-3’. *Nipbl* primer 1: 5’-AGTCCATATGCCCCACAGAG-3’. *Nipbl* primer 2: 5’-ACCGGCAACAATAGGACTTG-3’. *Nanog* primer 1: 5’-AAATCCCTTCCCTCGCCATC-3’. *Nanog* primer 2: 5’-GCCCTGACTTTAAGCCCAGA-3’. *Pou5f1* primer 1: 5’-CACCCTGGGCGTTCTCTTT-3’. *Pou5f1* primer 2: 5’-GTCTCCGATTTGCATATCTCCTG-3’.

### Identification of Genes with Nanog Binding Sites

In Avsec et al., 2021 (*119*), ChIP-seq was performed for Nanog in mESCs and 56,459 peaks were called (Fig. 8F). We considered the called peaks from that study as representing Nanog binding sites. Using the narrowPeak file from that study (which contains the called peaks), the GRCm38/mm10 C57BL/6J Mus musculus genome annotation, and ChIPseeker v3, we identified which genes had a peak +/- 250 nt of their TSS. We identified a total of 3,646 genes with a Nanog binding site at their TSS (Fig. 8F).

### Alcian Blue – Alizarin Red Staining

E18.5 embryos were fixed in 10% neutral buffered formalin in scintillation vials at 4°C overnight. Embryos were washed with H_2_O twice over 2 days at room temperature. Embryos were immersed in 95% EtOH for 1 week. Skin was removed from embryos with forceps under dissection microscope. Embryos were stained with 0.02% Alcian Blue (10 mg Alcian Blue + 80 ml EtOH + 20 ml glacial acetic acid) over three days. Embryos were washed with 70% EtOH twice in 1 day, 40% EtOH overnight, 15% EtOH in 1 day, and H_2_O overnight. Embryos were washed with 1% KOH twice over 3 days until they became translucent. Embryos were stained with 0.015% Alizarin Red (15 mg Alizarin Red + 100 ml 1% KOH) over 3 days. Embryos were washed with 1% KOH three times in 1 day. Fat pads and internal organs were removed from embryos with forceps under dissection microscope. Embryos were washed with 20% glycerol + 1% KOH overnight, 50% glycerol + 1% KOH overnight, and 80% glycerol + 1% KOH overnight. Embryos were stored in 100% glycerol at room temperature.

## Supporting information

Supplementary Figures S1-S15, Tables S1-S11, and Legends for Supplementary Data S1-S45

## Acknowledgments

The authors are grateful to Henry Chang, Lorelle Meidt, Han Nguyen, Arianna Favela, Christopher Wu, Louise Villagomez, and Brian Bui for assistance with mouse husbandry, genotyping, and data analysis; to Melanie Oakes of the UCI Genomics and Research Technology Hub (GRTH) for 10x library preparation and sequencing; and to Qing Nie (UCI Depts of Mathematics and Developmental & Cell Biology) and Rosaysela Santos (Kaiser Permanente Bernard J. Tyson School of Medicine) for helpful discussions. The authors are grateful to the UCI Center for Complex Biological Systems and the NSF-Simons Center for Multiscale Cell Fate Research for support of early pilot studies in this project. SC acknowledges support from a training grant award (NINDS/NIH T32NS082174), and JK acknowledges support by a Momental Foundation Mistletoe Research Fellowship. ADL and ALC dedicate this paper to the memory of Isabel Eva Calof Lander.

## Funding

National Institutes of Health R01HL138659 (ALC, ADL)

National Institutes of Health R01DE019638 (ADL)

National Institutes of Health R35GM143019 (ALM)

## Author contributions

Conceptualization: ALC, ADL, SC, ALM, JK

Methodology: ADL, SC, ALC, ALM

Software: SC, ADL, JK, ALM

Validation: ALC, ADL, SC, ALM

Formal Analysis: SC, ADL, ALC, JK, ALM

Investigation: SC, JK, MEL

Resources: ADL, ALC

Data Curation: SC

Writing – Original Draft: SC, ALC, ADL

Writing – Review & Editing: ADL, SC, ALC, ALM, JK

Visualization: SC, ADL, ALC

Supervision: ALC, ADL, ALM

Project Administration: ALC, ADL

Funding Acquisition: ALC, ADL, ALM

## Data and materials availability

All raw and processed data are available in the main text, Sequence Read Archive (Accession: SRP265522) and Gene expression Omnibus (Accession: GSE151589). All Supplementary Figures and Tables cited in the main text have been uploaded under Supplementary Material as a single PDF file. All Supplementary Data cited in the main text have been uploaded under Auxiliary Supplementary Materials as a single ZIP file. No restrictions have been made on the availability of the data. All code is available in Zenodo (DOI: 10.5281/zenodo.10001570). *Nipbl^Flox/+^* mice will be provided by ALC and ADL pending scientific review and a completed Material Transfers Agreement.

