## Supplementary Figures S1-S15, Tables S1-S11, and Legends for Supplementary Data S1-S45 for "Gastrulation-stage gene expression in *Nipbl*^+/-^ mouse embryos foreshadows the development of syndromic birth defects"

Science Advances

AAAS

Stephenson Chea *et al.*

**This PDF file includes:**

Figs. S1 to S15
Tables S1 to S11
Data S1 to S45

**Other Supplementary Materials for this manuscript include the following:**

Data S1 to S45

**Table of Contents**

| | |
|---|---|
| Fig. S1 | Images of LB- and CC-stage WT and *Nipbl*$^{+/-}$ mouse embryos used for scRNAseq |
| Fig. S2 | Metrics used to identify low-quality cells and doublets among LB- and CC-stage embryos |
| Fig. S3 | Expression of Cre, Bgeo, and Nipbl in LB- and CC-stage WT and *Nipbl*$^{+/-}$ embryos |
| Fig. S4 | Batch effect correction and integration of cells of the same stage (LB- or CC-stage) and genotype (WT or *Nipbl*$^{+/-}$) |
| Fig. S5 | Example of optimized clustering using clustering tree and Shannon Entropy |
| Fig. S6 | Clustering of *Nipbl*$^{+/-}$ cells from LB-stage embryos and projection of WT cells of the same stage onto them |
| Fig. S7 | Images of EB- and EHF-stage WT mouse embryos used for scRNAseq |
| Fig. S8 | Metrics used to identify low-quality cells and doublets among EB- and EHF-stage embryos |
| Fig. S9 | Batch effect correction and integration of cells of the same stage (EB- or EHF-stage) |
| Fig. S10 | Clustering of EB- and EHF-stage cells from WT embryos and annotation thereof as either ectoderm, mesoderm, or endoderm |
| Fig. S11 | Expression of genes associated with apoptosis in *Nipbl*$^{+/-}$ embryos relative to WT |
| Fig. S12 | Expression of meta-PCNA genes and cell cycle phase composition of *Nipbl*$^{+/-}$ embryos relative to WT |
| Fig. S13 | Paraxial mesoderm lineage of LB-stage *Nipbl*$^{+/-}$ embryos underexpresses paraxial mesoderm anti-drivers |
| Fig. S14 | Differentially expressed genes between germ layers of LB-stage WT and *Nipbl*$^{+/-}$ embryos following projection of WT cells onto *Nipbl*$^{+/-}$ germ layers |
| Fig. S15 | CC-stage *Nipbl*$^{+/-}$ mice replicate the gene expression changes of Nanog overexpression |
| Table S1 | Number of cells, transcripts, and genes captured per LB- and CC-stage WT and *Nipbl*$^{+/-}$ embryos |

| | |
|---|---|
| Table S2 | Differential overexpression of markers of germ layer identity by cell populations in LB- and CC-stage embryos |
| Table S3 | Differential overexpression of markers of biological identity by endodermal cell populations in LB- and CC-stage embryos |
| Table S4 | Differential overexpression of markers of biological identity by ectodermal cell populations in LB- and CC-stage embryos |
| Table S5 | Differential overexpression of markers of biological identity by mesodermal cell populations in LB- and CC-stage embryos |
| Table S6 | Differential overexpression of markers of germ layer identity by cell populations in LB-stage *Nipbl*$^{+/-}$ embryos |
| Table S7 | Differential overexpression of markers of biological identity by mesodermal cell populations in LB-stage *Nipbl*$^{+/-}$ embryos |
| Table S8 | Number of cells, transcripts, and genes captured by scRNAseq of EB- and EHF-stage WT embryos |
| Table S9 | Differential overexpression of markers of germ layer identity by cell populations in EB- and EHF-stage embryos |
| Table S10 | Expression of *Nanog* in LB- and CC-stage embryos |
| Table S11 | Expression of *Pou5f1* in LB- and CC-stage embryos |
| Data S1 | Differentially expressed genes among clusters of LB-stage WT embryos |
| Data S2 | Differentially expressed genes among clusters of CC-stage WT embryos |
| Data S3 | Differentially expressed genes among endodermal clusters of LB-stage WT embryos |
| Data S4 | Differentially expressed genes among endodermal clusters of CC-stage WT embryos |
| Data S5 | Differentially expressed genes among ectodermal clusters of LB-stage WT embryos |
| Data S6 | Differentially expressed genes among ectodermal clusters of CC-stage WT embryos |
| Data S7 | Differentially expressed genes among mesodermal clusters of LB-stage WT embryos |

| | |
|---|---|
| Data S8 | Differentially expressed genes among mesodermal clusters of CC-stage WT embryos |
| Data S9 | Numbers of cells per germ layer per LB-stage embryo |
| Data S10 | Numbers of cells per mesodermal cell population per LB-stage embryo |
| Data S11 | Differentially expressed genes among clusters of LB-stage *Nipbl*$^{+/-}$ embryos |
| Data S12 | Differentially expressed genes among mesodermal clusters of LB-stage *Nipbl*$^{+/-}$ embryos |
| Data S13 | Numbers of cells per germ layer per LB-stage embryo following projection of WT cells onto *Nipbl*$^{+/-}$ germ layers |
| Data S14 | Numbers of cells per mesodermal cell population per LB-stage embryo following projection of WT cells onto *Nipbl*$^{+/-}$ mesodermal cell populations |
| Data S15 | Differentially expressed genes among clusters of EB-stage WT embryos |
| Data S16 | Differentially expressed genes among clusters of EHF-stage WT embryos |
| Data S17 | Pseudotime values of cells from EB-, LB-, and CC-stage embryos |
| Data S18 | RNA velocity vectors of mesoderm cells from LB-stage WT embryos |
| Data S19 | RNA velocity vectors of mesoderm cells from LB-stage *Nipbl*$^{+/-}$ embryos |
| Data S20 | Fate probabilities of mesoderm cells into first heart field, second heart field, and paraxial mesoderm fates from LB-stage WT embryos |
| Data S21 | Fate probabilities of mesoderm cells into first heart field, second heart field, and paraxial mesoderm fates from LB-stage *Nipbl*$^{+/-}$ embryos |
| Data S22 | Differentially expressed genes from Reactome, Hallmark, and Langemeijer Apoptosis Gene Set between germ layers of LB-stage WT and *Nipbl*$^{+/-}$ embryos |
| Data S23 | Differentially expressed genes from meta-PCNA Gene Set between germ layers of LB-stage WT and *Nipbl*$^{+/-}$ embryos |
| Data S24 | Cell cycle scores and phases of cells in germ layers of LB-stage WT and *Nipbl*$^{+/-}$ embryos |
| Data S25 | First heart field drivers and anti-drivers predicted by CellRank |
| Data S26 | Paraxial mesoderm drivers and anti-drivers predicted by CellRank |

| | |
|---|---|
| Data S27 | Differentially expressed genes between first heart field lineage of LB-stage WT and *Nipbl*$^{+/-}$ embryos |
| Data S28 | Differentially expressed genes between paraxial mesoderm lineage of LB-stage WT and *Nipbl*$^{+/-}$ embryos |
| Data S29 | Differentially expressed genes between mesoderms of LB-stage WT and *Nipbl*$^{+/-}$ embryos |
| Data S30 | Differentially expressed genes between ectoderms of LB-stage WT and *Nipbl*$^{+/-}$ embryos |
| Data S31 | Differentially expressed genes between endoderms of LB-stage WT and *Nipbl*$^{+/-}$ embryos |
| Data S32 | Differentially expressed genes between mesoderms of LB-stage WT and *Nipbl*$^{+/-}$ embryos following projection of WT cells onto *Nipbl*$^{+/-}$ germ layers |
| Data S33 | Differentially expressed genes between ectoderms of LB-stage WT and *Nipbl*$^{+/-}$ embryos following projection of WT cells onto *Nipbl*$^{+/-}$ germ layers |
| Data S34 | Differentially expressed genes between endoderms of LB-stage WT and *Nipbl*$^{+/-}$ embryos following projection of WT cells onto *Nipbl*$^{+/-}$ germ layers |
| Data S35 | Network of gene interactions predicted by STRING for genes differentially expressed more than two-fold up or down in germ layers of LB-stage *Nipbl*$^{+/-}$ embryos than WT embryos |
| Data S36 | Expression of *Nanog* in mesodermal cell populations of LB-stage embryos |
| Data S37 | Expression of *Pou5f1* in mesodermal cell populations of LB-stage embryos |
| Data S38 | qRT-PCR results for *Nipbl*, *Nanog*, and *Pou5f1* in *Nipbl*$^{FLEX/+}$ and *Nipbl*$^{Flrt/+}$ ES cells |
| Data S39 | Differentially expressed genes between whole LB-stage WT and *Nipbl*$^{+/-}$ embryos |
| Data S40 | Differentially expressed genes between whole CC-stage WT and *Nipbl*$^{+/-}$ embryos |
| Data S41 | Differentially expressed genes between mesoderms of CC-stage WT and *Nipbl*$^{+/-}$ embryos |
| Data S42 | Differentially expressed genes between ectoderms of CC-stage WT and *Nipbl*$^{+/-}$ embryos |

| | |
|---|---|
| Data S43 | Differentially expressed genes between endoderms of CC-stage WT and *Nipbl*$^{+/-}$ embryos |
| Data S44 | Expression of *Hox* genes in mesoderms of LB- and CC-stage embryos |
| Data S45 | Expression of *Hox* genes in ectoderms of LB- and CC-stage WT embryos |

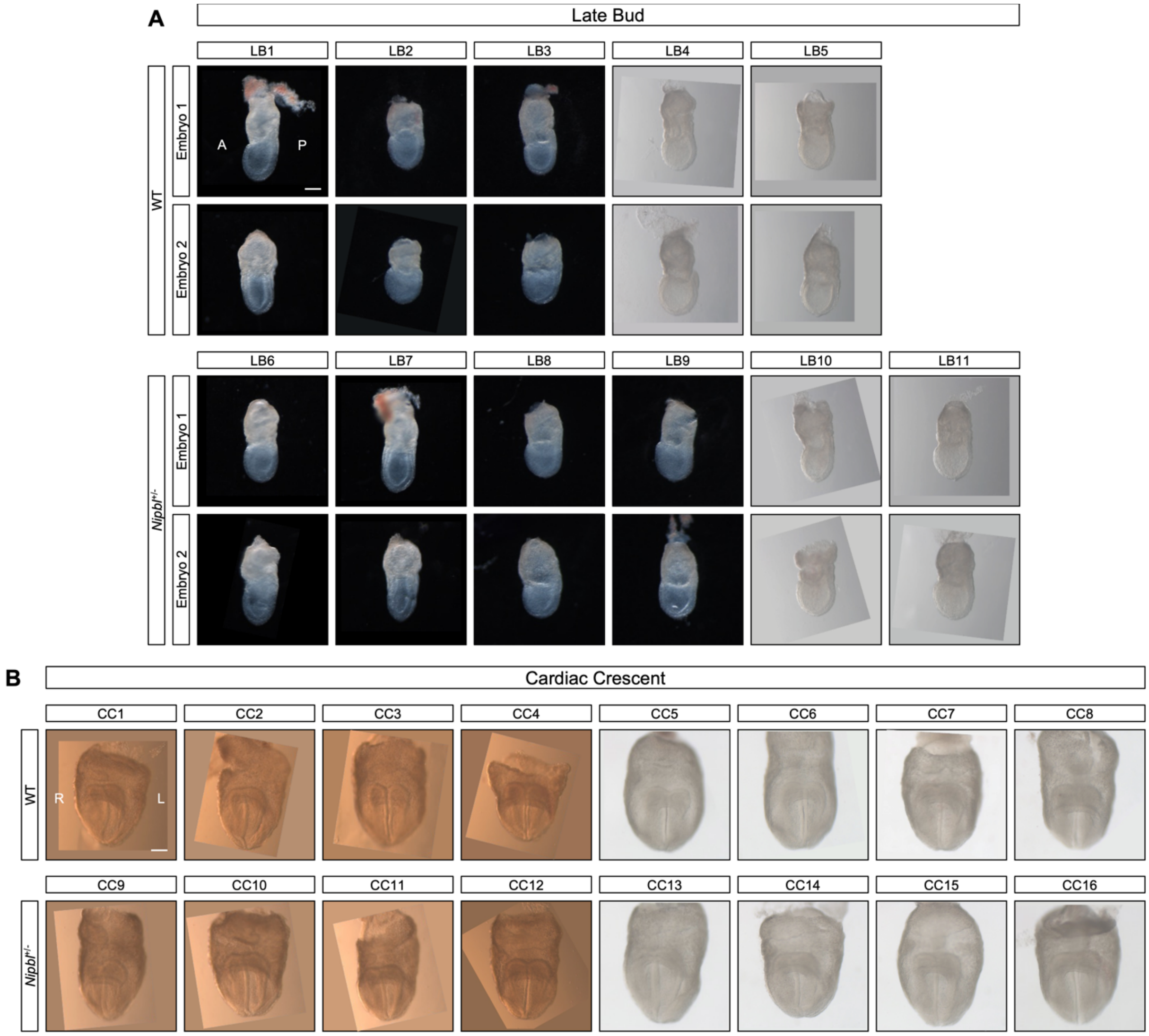

**Figure S1. Images of LB- and CC-stage WT and *Nipbl*$^{+/-}$ mouse embryos used for scRNAseq.**

Lateral view of (**A**) LB-stage and (**B**) anterior view of CC-stage WT and *Nipbl*$^{+/-}$ mouse embryos. A = anterior, P = posterior, L = left, R = right, and scale bar = 100 um.

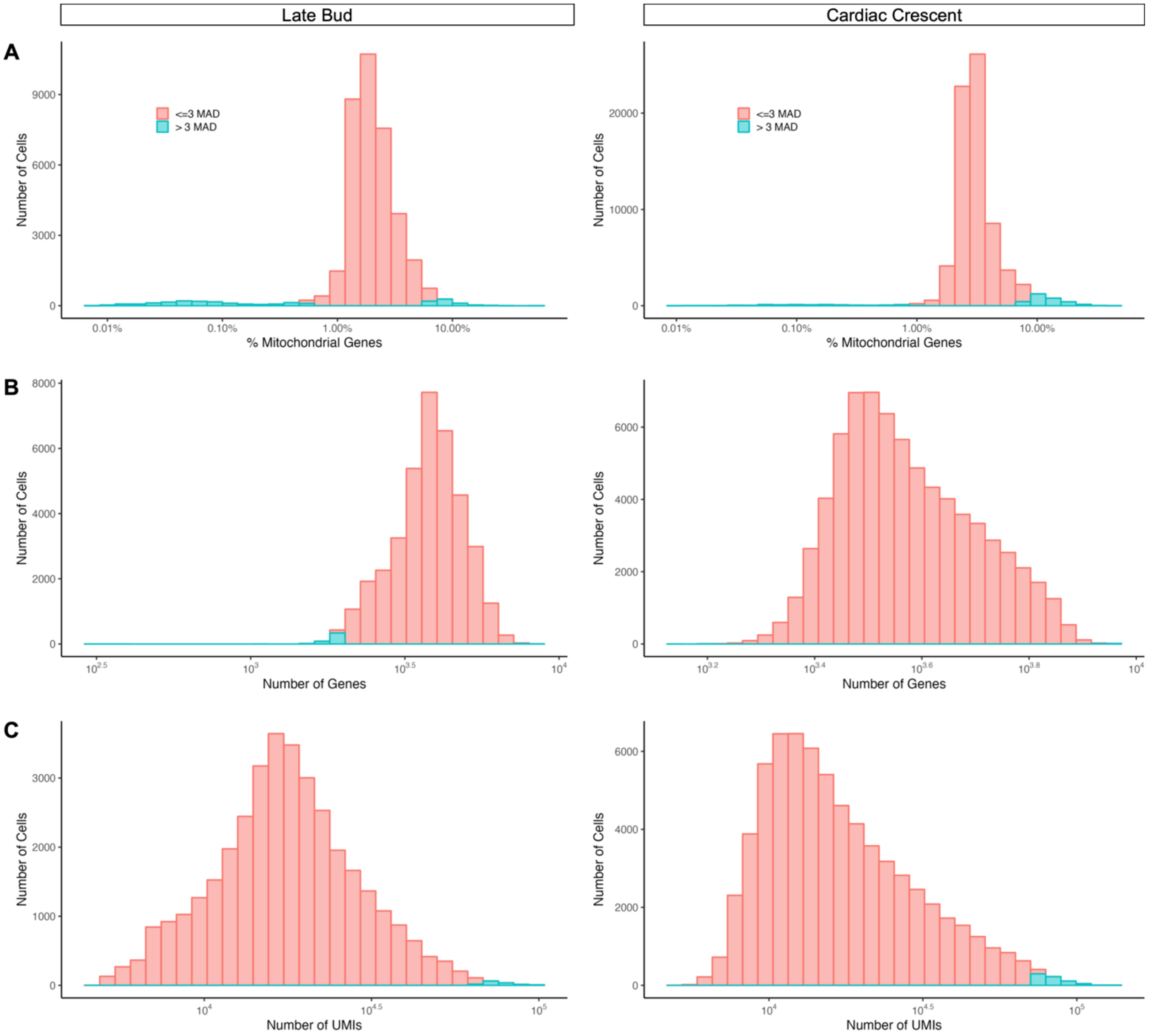

**Figure S2. Metrics used to identify low-quality cells and doublets among LB- and CC-stage embryos.**

At each stage, we considered cells that exceeded three median absolute deviations from percent mitochondrial genes expressed, number of genes expressed, and number of transcripts detected per cell, low-quality cells and/or doublets and removed them. Histograms of (**A**) percent mitochondrial genes expressed, (**B**) number of genes expressed, and (**C**) number of transcripts detected per cell at LB- and CC-stage colored by whether cells were less than or equal to 3 median absolute deviations (red) or greater than 3 median absolute deviations (blue).

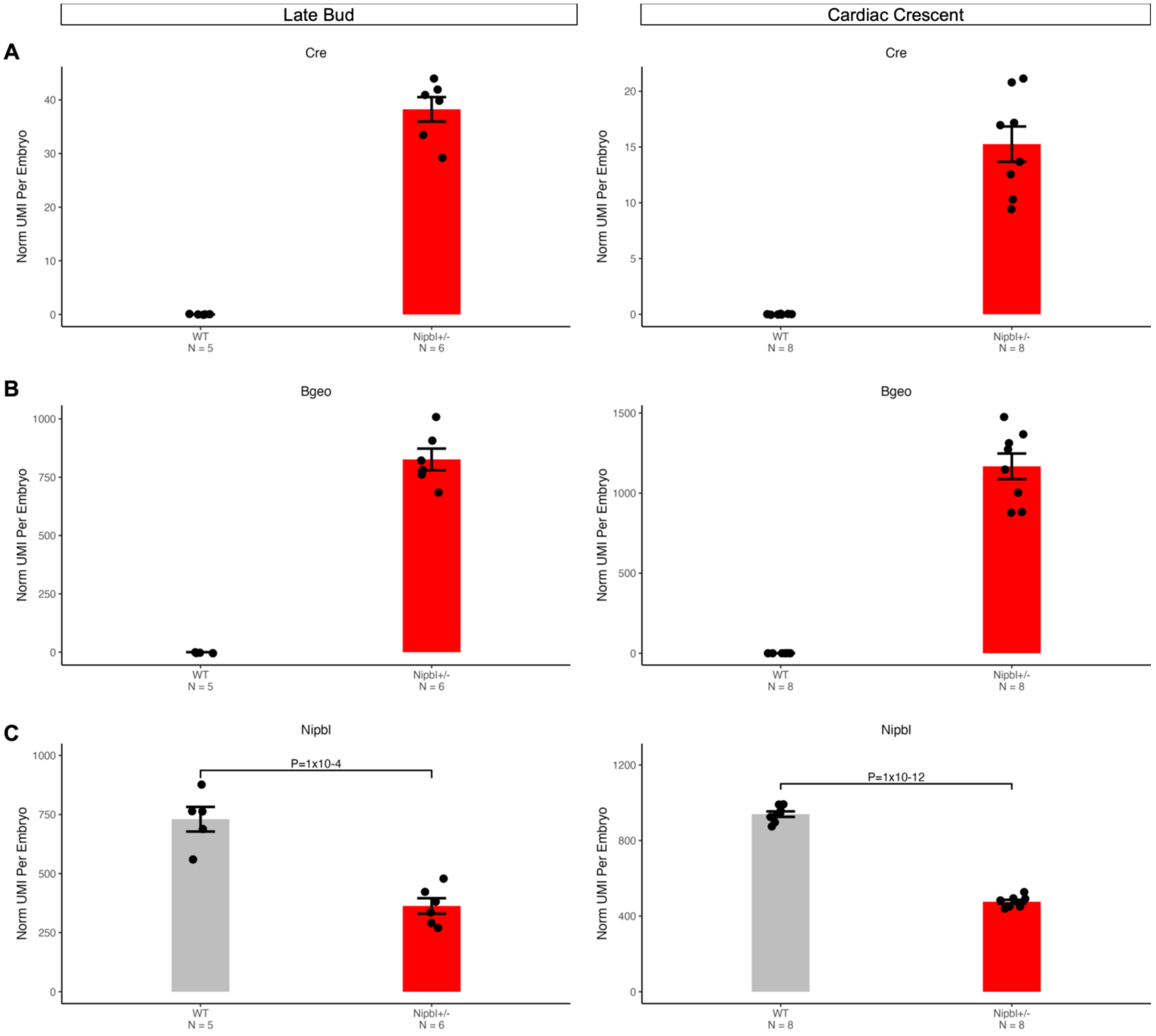

**Figure S3. Expression of Cre, Bgeo, and Nipbl in LB- and CC-stage WT and *Nipbl*$^{+/-}$ embryos.**

*Nipbl*$^{+/-}$ embryos at LB- and CC-stage express Cre and Bgeo, while WT embryos do not. *Nipbl*$^{+/-}$ embryos also express *Nipbl* at levels lower than that of WT embryos. Normalized transcripts of (A) *Cre*, (B) *Bgeo*, and (C) *Nipbl* in LB- and CC-stage embryos. Each dot indicates an individual sample. Error bars show standard error of the mean. *P*-value from T-Test.

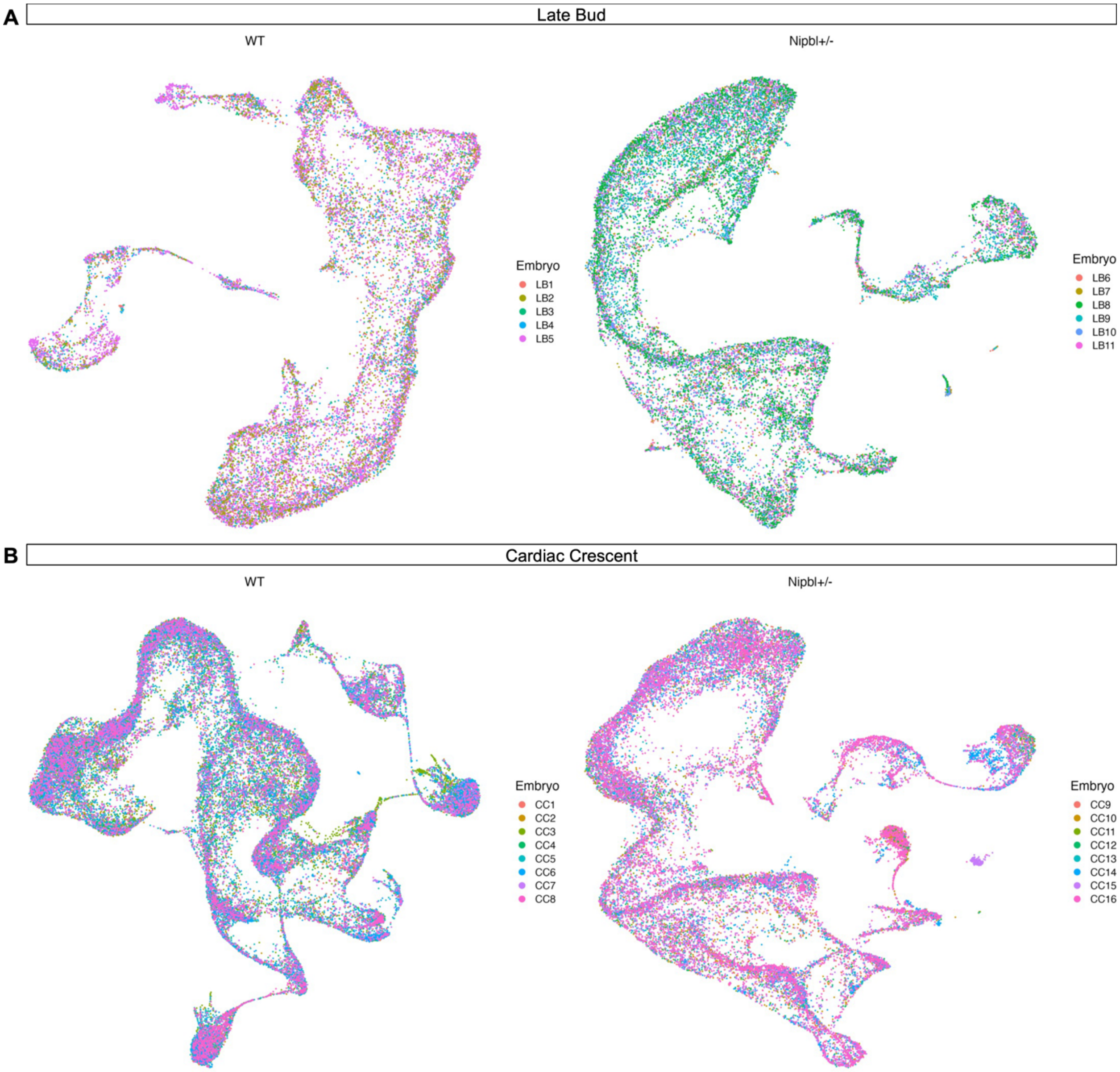

**Fig. S4. Batch effect correction and integration of cells of the same stage (LB- or CC-stage) and genotype (WT or *Nipbl*$^{+/-}$).**

We corrected batch effects among embryos by integrating cells from the same stage and genotype together. UMAP of WT and *Nipbl*$^{+/-}$ cells colored by embryo at (A) LB- and (B) CC-stage.

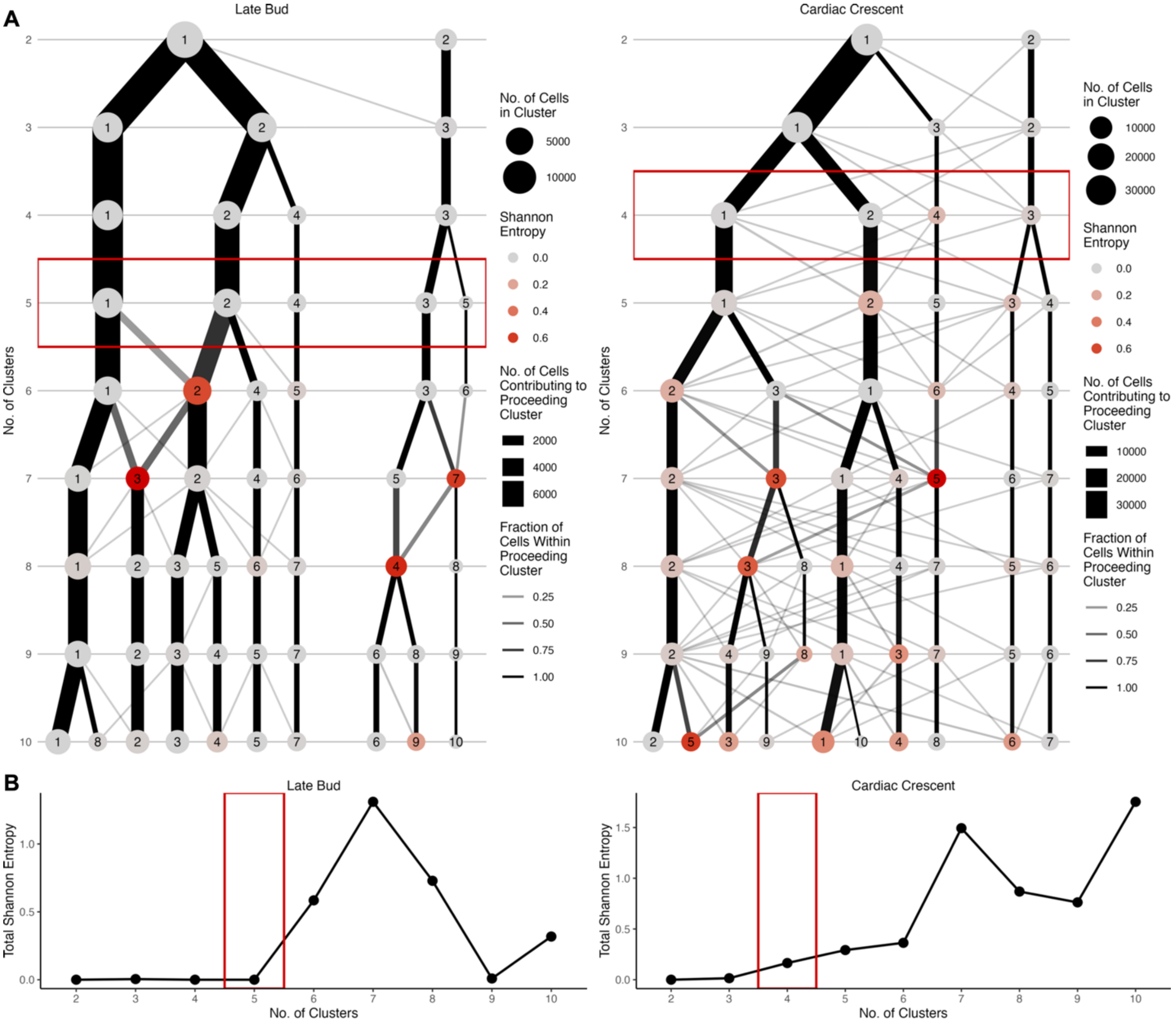

**Fig. S5. Example of optimized clustering using clustering tree and Shannon Entropy.**

(**A**) Following consecutive clustering of WT cells from, we generated a clustering tree to visualize how cell cluster identities change as the number of clusters consecutively increase. Nodes represent clusters and edges represent cells from preceding clusters. Clusters are stable when a large proportion of cells are derived from a single preceding cluster rather than multiple preceding clusters. We adopted Shannon Entropy as a measure of these proportions as a measure of intra-cluster stability. A low Shannon Entropy represents high intra-cluster stability. Here, WT cells from LB-stage embryos can only be clustered as high as 5 clusters before they can be subclustered into further subclusters. WT cells from CC-stage embryos can only be clustered as high as 4 clusters before they can be subclustered into further clusters. (**B**) Total Shannon Entropy of all clusters among WT cells from LB- or CC-stage embryos at consecutively increasing number of clusters.

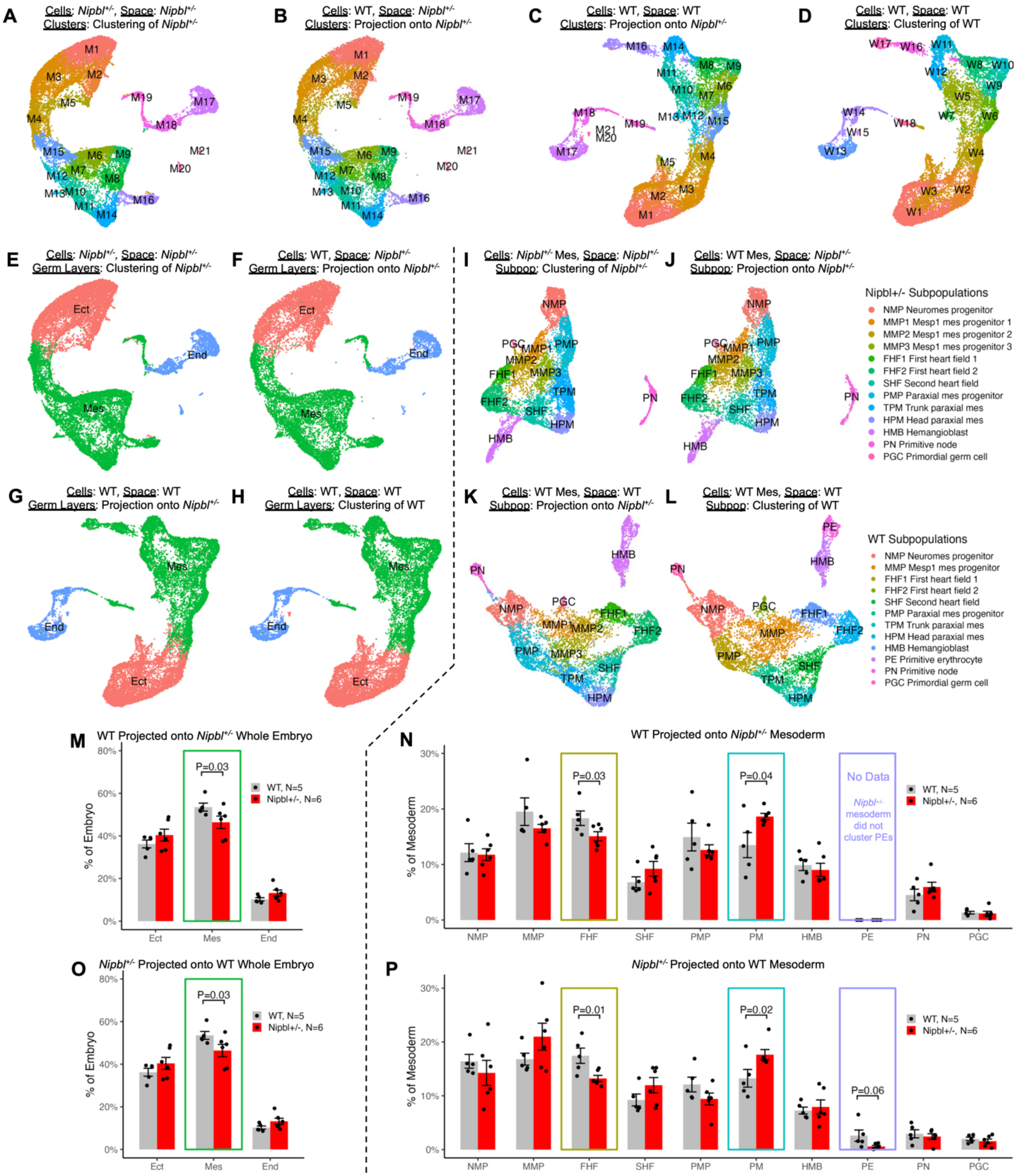
A
Cells: Nipbl+/-, Space: Nipbl+/-
Clusters: Clustering of Nipbl+/-
B
Cells: WT, Space: Nipbl+/-
Clusters: Projection onto Nipbl+/-
C
Cells: WT, Space: WT
Clusters: Projection onto Nipbl+/-
D
Cells: WT, Space: WT
Clusters: Clustering of WT
E
Cells: Nipbl+/-, Space: Nipbl+/-
Germ Layers: Clustering of Nipbl+/-
F
Cells: WT, Space: Nipbl+/-
Germ Layers: Projection onto Nipbl+/-
G
Cells: WT, Space: WT
Germ Layers: Projection onto Nipbl+/-
H
Cells: WT, Space: WT
Germ Layers: Clustering of WT
Ect
Mes
End
I
Cells: Nipbl+/- Mes, Space: Nipbl+/-
Subpop: Clustering of Nipbl+/-
J
Cells: WT Mes, Space: Nipbl+/-
Subpop: Projection onto Nipbl+/-
Nipbl+/- Subpopulations
NMP Neuromes progenitor
MMP1 Mesp1 mes progenitor 1
MMP2 Mesp1 mes progenitor 2
MMP3 Mesp1 mes progenitor 3
FHF1 First heart field 1
FHF2 First heart field 2
SHF Second heart field
PMP Paraxial mes progenitor
TPM Trunk paraxial mes
HPM Head paraxial mes
HMB Hemangioblast
PN Primitive node
PGC Primordial germ cell
K
Cells: WT Mes, Space: WT
Subpop: Projection onto Nipbl+/-
L
Cells: WT Mes, Space: WT
Subpop: Clustering of WT
WT Subpopulations
NMP Neuromes progenitor
MMP Mesp1 mes progenitor
FHF1 First heart field 1
FHF2 First heart field 2
SHF Second heart field
PMP Paraxial mes progenitor
TPM Trunk paraxial mes
HPM Head paraxial mes
HMB Hemangioblast
PE Primitive erythrocyte
PN Primitive node
PGC Primordial germ cell
M
WT Projected onto Nipbl+/- Whole Embryo
% of Embryo
P=0.03
WT, N=5
Nipbl+/-, N=6
Ect
Mes
End
N
WT Projected onto Nipbl+/- Mesoderm
% of Mesoderm
P=0.03
P=0.04
No Data
Nipbl+/- mesoderm did not cluster PEs
WT, N=5
Nipbl+/-, N=6
NMP
MMP
FHF
SHF
PMP
PM
HMB
PE
PN
PGC
O
Nipbl+/- Projected onto WT Whole Embryo
% of Embryo
P=0.03
WT, N=5
Nipbl+/-, N=6
Ect
Mes
End
P
Nipbl+/- Projected onto WT Mesoderm
% of Mesoderm
P=0.01
P=0.02
P=0.06
WT, N=5
Nipbl+/-, N=6
NMP
MMP
FHF
SHF
PMP
PM
HMB
PE
PN
PGC

**Fig. S6. Percentage of cells in germ layers and mesodermal subpopulations of LB-stage embryos following projection of WT cells onto clustered *Nipbl*$^{+/-}$ cells.**

(**A**) LB-stage *Nipbl*$^{+/-}$ cells in UMAP of *Nipbl*$^{+/-}$ cells colored by cluster after clustering *Nipbl*$^{+/-}$ cells separately from WT. (**B**) LB-stage WT cells in UMAP of *Nipbl*$^{+/-}$ cells colored by clusters found in *Nipbl*$^{+/-}$ cells after projecting WT cells onto *Nipbl*$^{+/-}$ cells. (**C**) LB-stage WT cells in UMAP of WT cells colored by clusters found in *Nipbl*$^{+/-}$ cells after projecting WT cells onto *Nipbl*$^{+/-}$ cells. (**D**) LB-stage WT cells in UMAP of WT cells colored by clusters after clustering WT cells separately from *Nipbl*$^{+/-}$. (**E**) LB-stage *Nipbl*$^{+/-}$ cells in UMAP of *Nipbl*$^{+/-}$ cells colored by germ layer after clustering *Nipbl*$^{+/-}$ cells separately from WT. (**F**) LB-stage WT cells in UMAP of *Nipbl*$^{+/-}$ cells colored by germ layers found in *Nipbl*$^{+/-}$ cells after projecting WT cells onto *Nipbl*$^{+/-}$ cells. (**G**) LB-stage WT cells in UMAP of WT cells colored by germ layers found in *Nipbl*$^{+/-}$ cells after projecting WT cells onto *Nipbl*$^{+/-}$ cells. (**H**) LB-stage WT cells in UMAP of WT cells colored by germ layer after clustering WT cells separately from *Nipbl*$^{+/-}$. (**I**) LB-stage *Nipbl*$^{+/-}$ mesoderm cells in UMAP of *Nipbl*$^{+/-}$ mesoderm colored by cell population after clustering *Nipbl*$^{+/-}$ mesoderm separately from WT. (**J**) LB-stage WT mesoderm cells in UMAP of *Nipbl*$^{+/-}$ mesoderm colored by cell populations found in *Nipbl*$^{+/-}$ mesoderm after projecting WT mesoderm onto *Nipbl*$^{+/-}$ mesoderm. (**K**) LB-stage WT mesoderm cells in UMAP of WT mesoderm colored by cell populations found in *Nipbl*$^{+/-}$ mesoderm after projecting WT mesoderm onto *Nipbl*$^{+/-}$ mesoderm. (**L**) LB-stage WT mesoderm cells in UMAP of WT mesoderm colored by cell population after clustering WT mesoderm separately from *Nipbl*$^{+/-}$. (**M**) Percentage of cells in germ layers from all cells in LB-stage WT and *Nipbl*$^{+/-}$ embryos after projecting WT cells onto *Nipbl*$^{+/-}$ cells. Error bars show standard error of the mean. *P*-values from T-Test. (**N**) Percentage of cells in mesodermal cell populations from all mesoderm cells in LB-stage WT and *Nipbl*$^{+/-}$ embryos after projecting WT mesoderm onto *Nipbl*$^{+/-}$ mesoderm. Error bars show SEM. *P*-values from T-Test. (**O**) Percentage of cells in germ layers from all cells in LB-stage WT and *Nipbl*$^{+/-}$ embryos after projecting *Nipbl*$^{+/-}$ cells onto WT cells. Same as This figure is from Fig. 3B. Error bars show standard error of the mean. *P*-values from T-Test. (**P**) Percentage of cells in mesodermal cell populations from all mesoderm cells in LB-stage WT and *Nipbl*$^{+/-}$ embryos after projecting *Nipbl*$^{+/-}$ mesoderm onto WT mesoderm. This figure is from Fig. 3D. Error bars show SEM. *P*-values from T-Test.

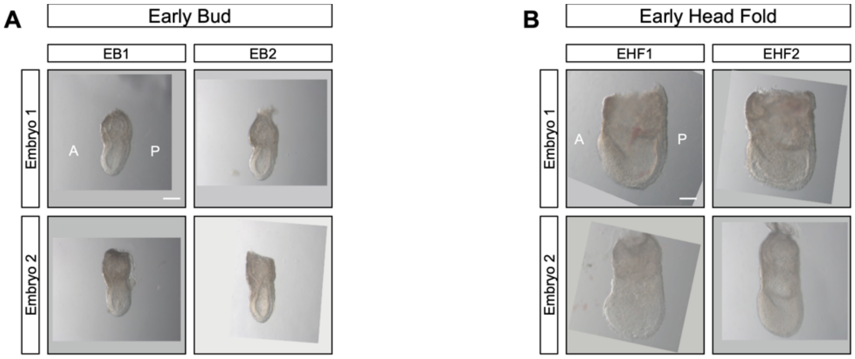

**Fig. S7. Images of EB- and EHF-stage WT mouse embryos used for scRNAseq.**

Lateral view of WT mouse embryos at (**A**) EB- and (**B**) EHF- stage. A = anterior, P = posterior. Dashed line represents where embryonic tissue was separated from extraembryonic tissue. Scale bar = 100 um.

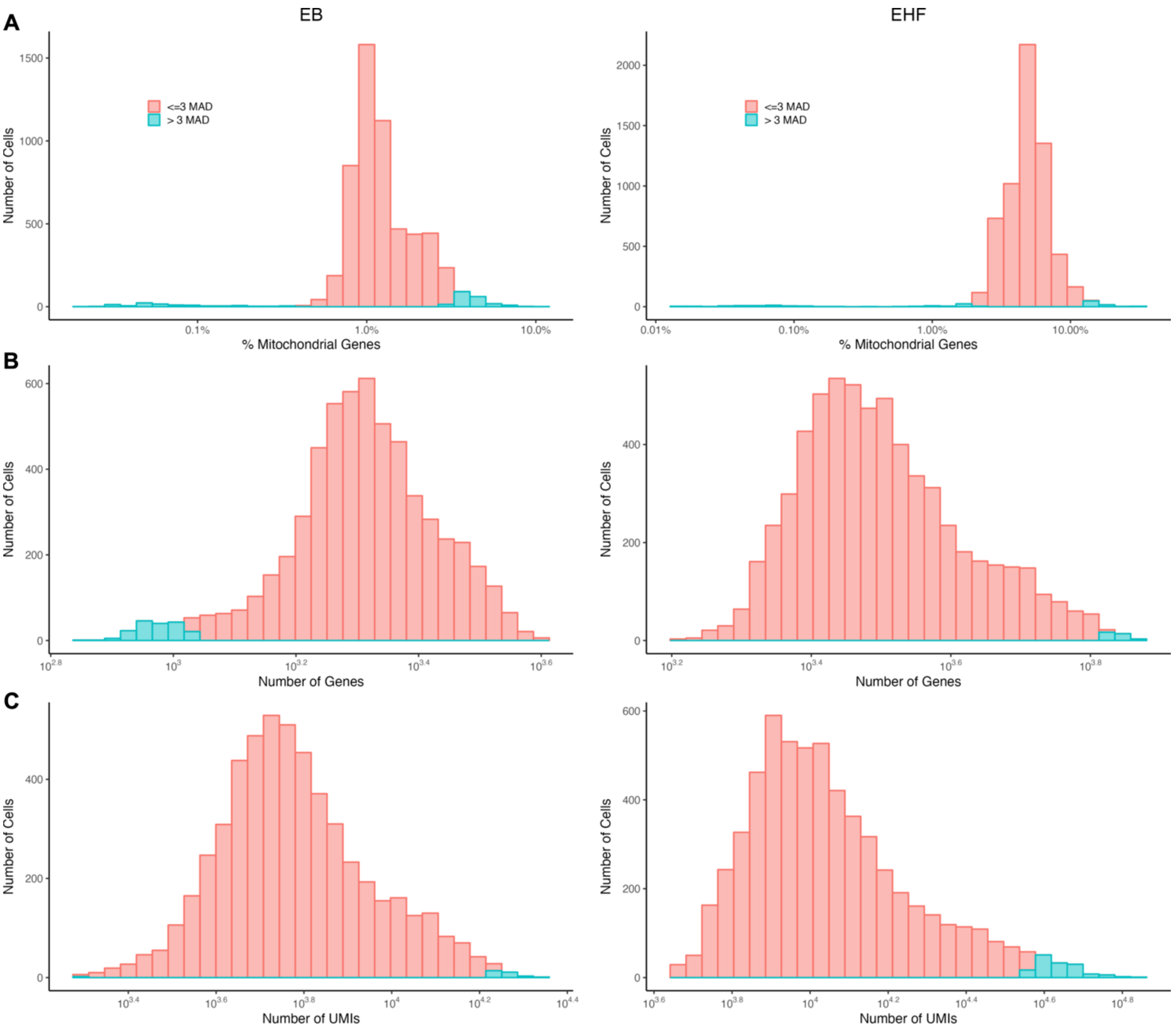

**Figure S8. Metrics used to identify low-quality cells and doublets among EB- and EHF-stage embryos.**

At each stage, we considered cells that exceeded three median absolute deviations from percent mitochondrial genes expressed, number of genes expressed, and number of transcripts detected per cell low-quality cells and/or doublets and removed them. Histograms of (**A**) percent mitochondrial genes expressed, (**B**) number of genes expressed, and (**C**) number of transcripts detected per cell at EB- and EHF-stage colored by whether cells were less than or equal to 3 median absolute deviations (red) or greater than 3 median absolute deviations (blue). Cells colored in blue were removed.

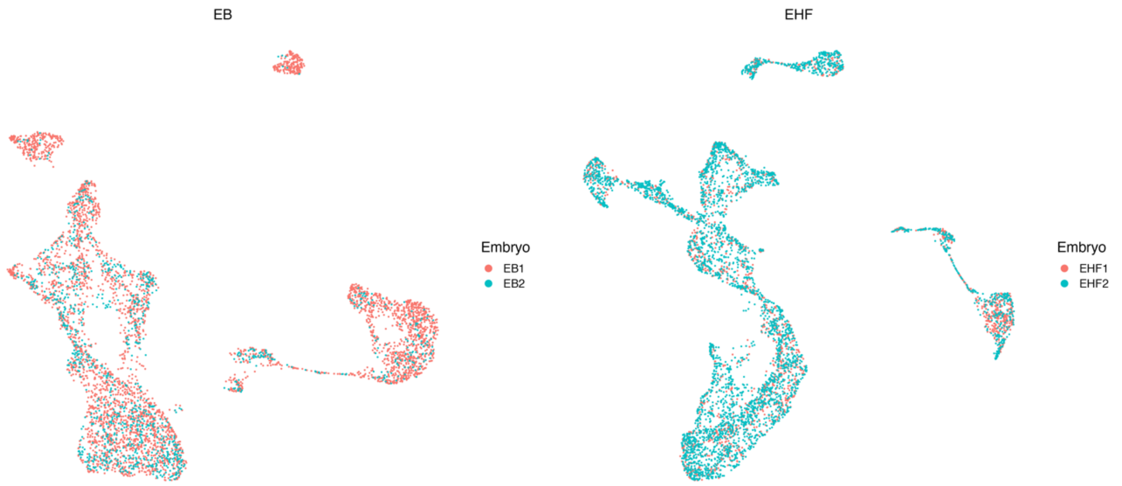

**Fig. S9. Batch effect correction and integration of cells of the same stage (EB- or EHF-stage).**

We removed batch effects among EB- and EHF-stage embryos of the same stage and integrated embryos of the same stage together. UMAP of EB- and EHF-stage cells colored by embryo.

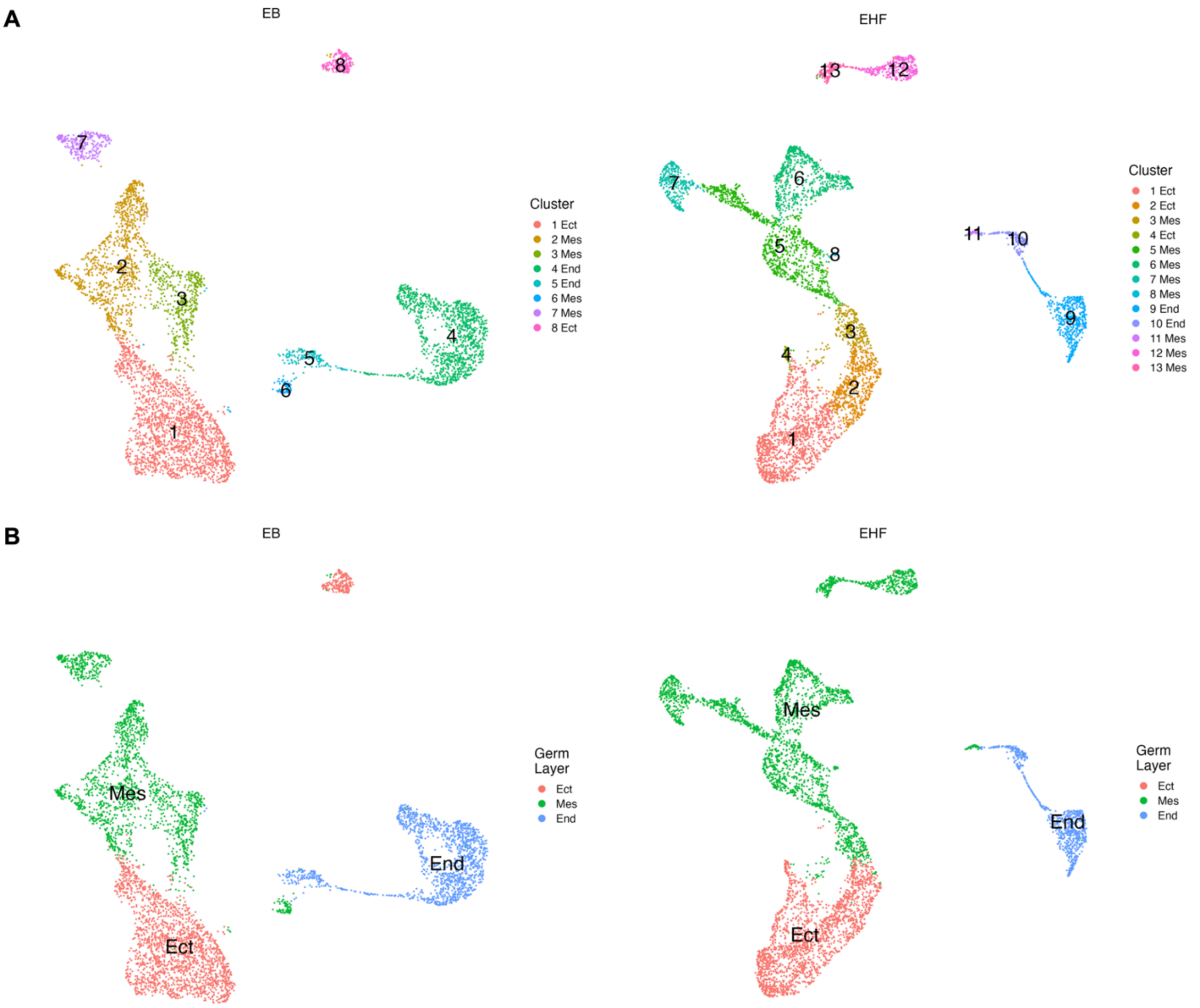

**Fig. S10. Clustering of EB- and EHF-stage cells from WT embryos and annotation thereof as either ectoderm, mesoderm, or endoderm.**

UMAP of EB- and EHF-stage cells colored by (**A**) cell population or (**B**) germ layer.

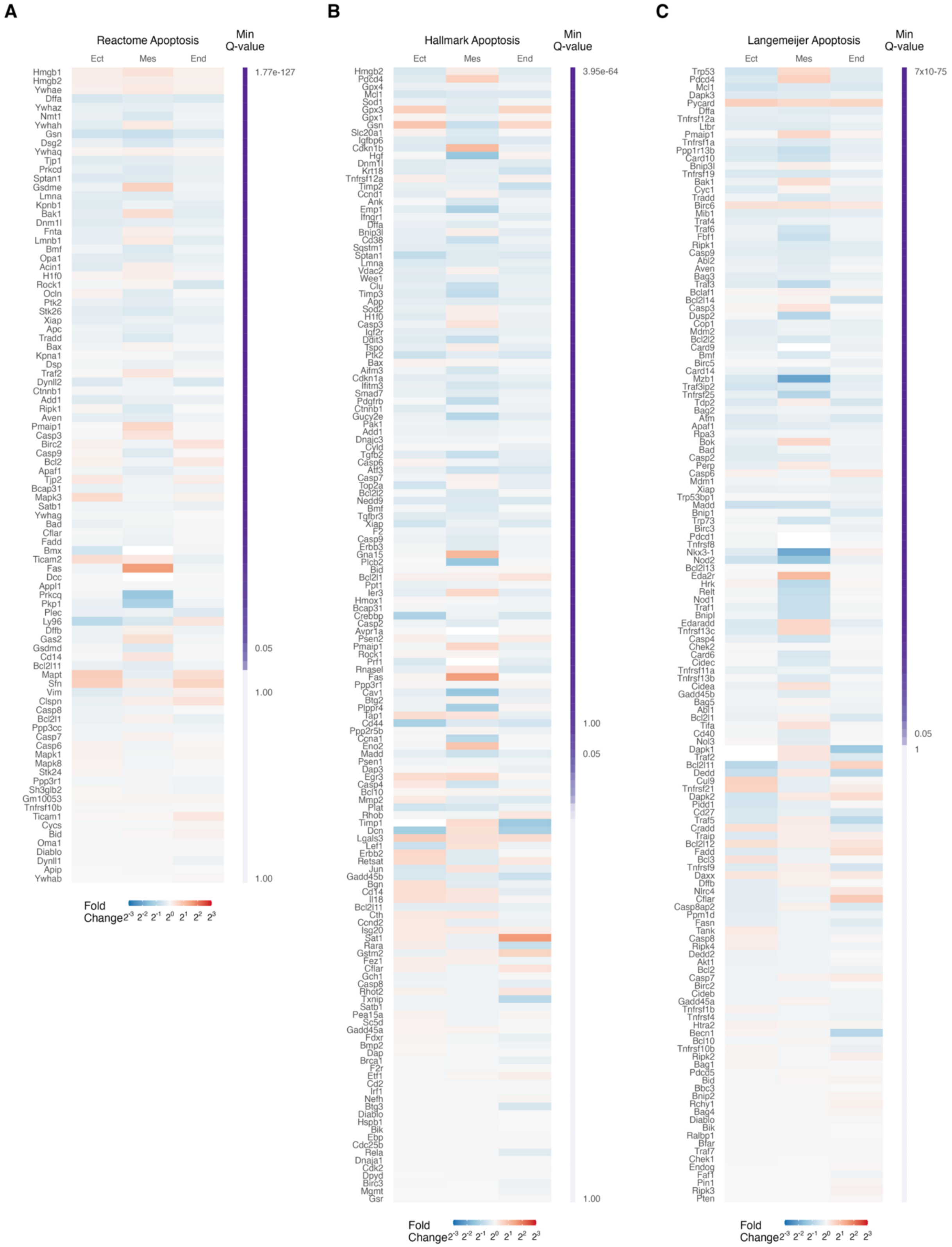
A
Reactome Apoptosis
Min Q-value
Ect
Mes
End
1.77e-127
0.05
1.00
1.00
Fold Change
B
Hallmark Apoptosis
Min Q-value
Ect
Mes
End
3.95e-64
1.00
0.05
1.00
Fold Change
C
Langemeijer Apoptosis
Min Q-value
Ect
Mes
End
7x10-75
0.05
1
Fold Change

**Fig. S11. Expression of genes associated with apoptosis in *Nipbl*$^{+/-}$ embryos relative to WT.**

The germ layers of *Nipbl*$^{+/-}$ embryos do not substantially overexpress or underexpress genes associated with apoptosis. Heatmap of fold change in expression in genes from the (**A**) Reactome Apoptosis, (**B**) Hallmark Apoptosis, and (**C**) Langemeijer Apoptosis gene sets in germ layers of *Nipbl*$^{+/-}$ embryos from that of WT embryos. Genes are ordered from top to bottom by minimum *Q*-value. *Q*-value from Bonferroni corrected *P*-value from Mann-Whitney U test.

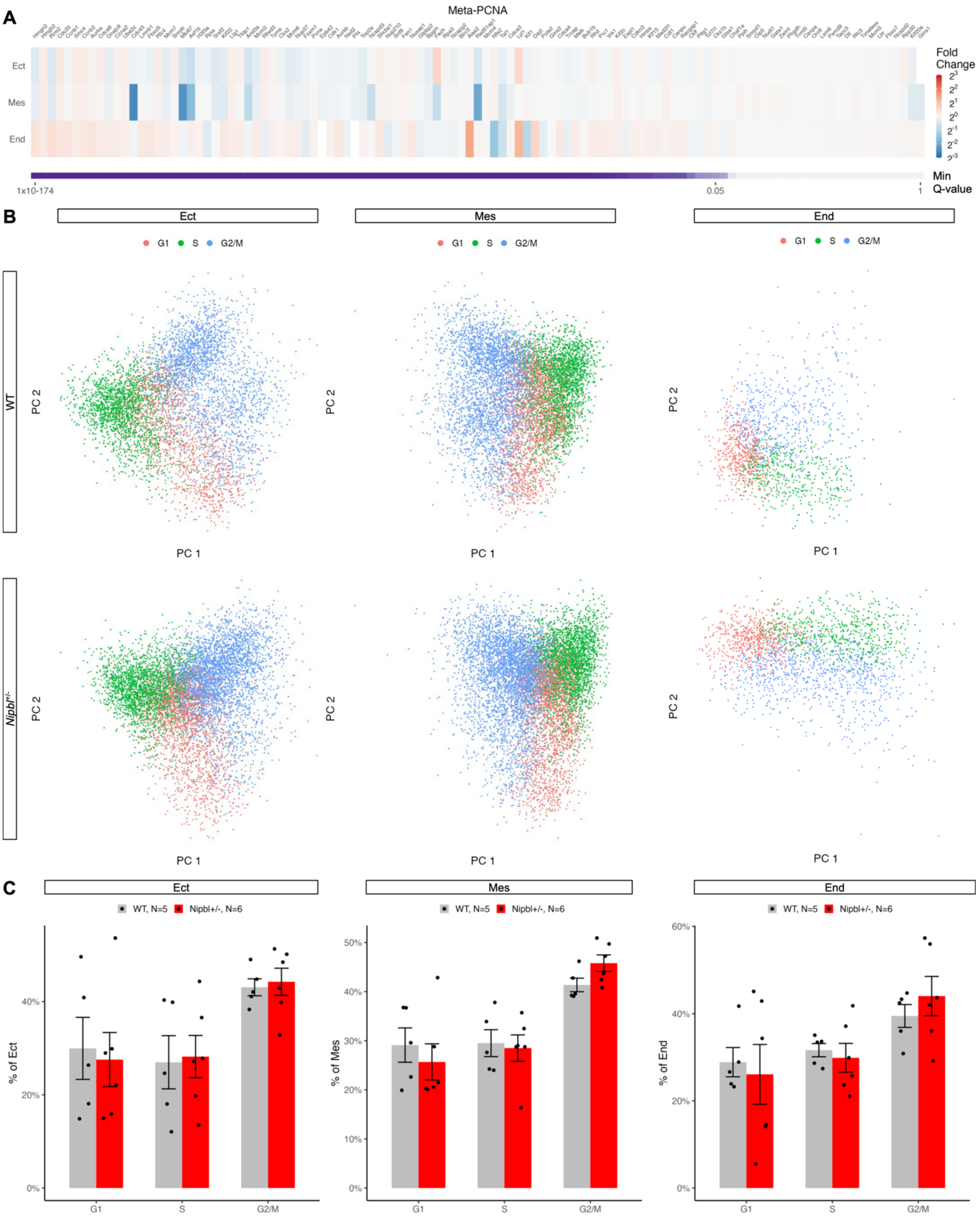
A
Meta-PCNA
Ect
Mes
End
Fold Change
Min Q-value
1x10-174
0.05
1
B
Ect
Mes
End
G1 S G2/M
WT
Nipbl+/-
PC 1
PC 2
C
WT, N=5 Nipbl+/-, N=6
% of Ect
% of Mes
% of End
G1
S
G2/M

**Fig. S12. Expression of meta-PCNA genes and cell cycle phase composition of *Nipbl*$^{+/-}$ embryos relative to WT.**

The germ layers of *Nipbl*$^{+/-}$ embryos do not substantially overexpress or underexpress genes associated with cell proliferation. (**A**) Heatmap of fold change in expression in genes from the meta-PCNA gene set in germ layers of *Nipbl*$^{+/-}$ embryos from that of WT embryos. Genes are ordered from top to bottom by minimum *Q*-value. *Q*-value from Bonferroni corrected *P*-value from Mann-Whitney U test. (**B**) Principal component analysis was performed cells from each germ layer of WT and *Nipbl*$^{+/-}$ embryos using cell cycle phase genes marking S phase and G2/M phase. Seurat was used to assign cells into G1, S, or G2/M phase based on the expression of these markers. Expression of these marker sets are considered anticorrelated. When cells express neither, they are considered to be in G1 phase. In all germ layers, cells from WT and *Nipbl*$^{+/-}$ embryos assigned into all three phases. (**C**) Percentage of cells in each cell cycle phase across all germ layers of WT and *Nipbl*$^{+/-}$ embryos. There was no statistically significant difference in the percentage of cells in each cell cycle phase between that of WT and *Nipbl*$^{+/-}$ embryos. Error bars show standard error of the mean.

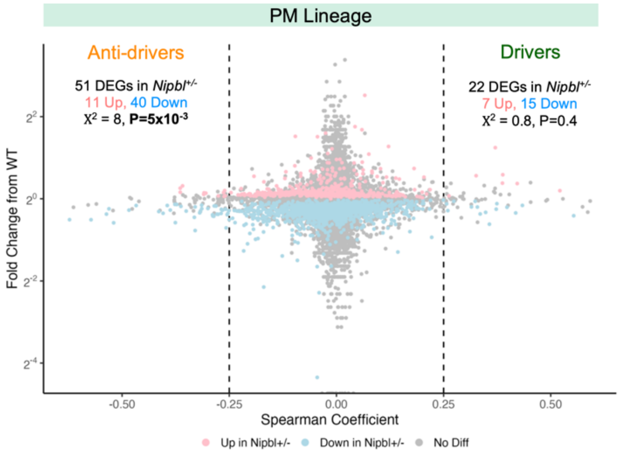

**Fig. S13. Paraxial mesoderm lineage of LB-stage *Nipbl*$^{+/-}$ embryos, showing underexpression of paraxial mesoderm anti-drivers.** Fold change in expression of genes in mesoderm cells of PM lineage of LB-stage *Nipbl*$^{+/-}$ embryos differentially expressed ($Q < 0.05$, Mann-Whitney U Test) from that of WT embryos along Spearman's Rank Correlation coefficient from Fig. 5E.

### Mes

**A**

| | | WT Projected onto *Nipbl*+/- **(Reverse Projection)** | | |
|---|---|---|---|---|
| | | Up in *Nipbl*+/- (613) | Down in *Nipbl*+/- (2,900) | No Diff in *Nipbl*+/- (12,851) |
| *Nipbl*+/- Projected onto WT **(Forward Projection)** | Up in *Nipbl*+/- (604) | 588 **(95.9%)** | 0 (0.0%) | 16 (0.1%) |
| | Down in *Nipbl*+/- (2,840) | 2 (0.3%) | 2,743 **(94.6%)** | 95 (0.7%) |
| | No Diff in *Nipbl*+/- (12,920) | 23 (3.8%) | 157 (5.4%) | 12,740 **(99.1%)** |

**D**

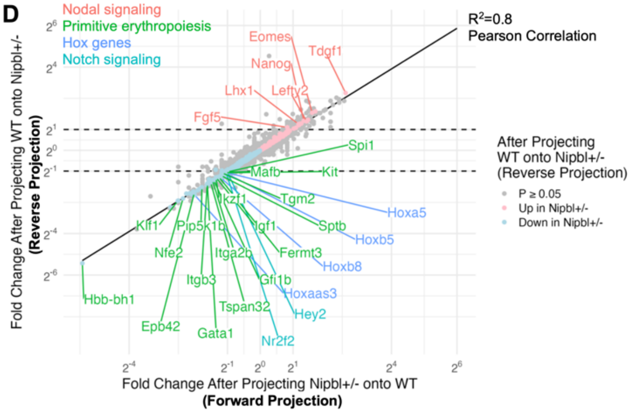

### Ect

**B**

| | | WT Projected onto *Nipbl*+/- **(Reverse Projection)** | | |
|---|---|---|---|---|
| | | Up in *Nipbl*+/- (683) | Down in *Nipbl*+/- (3,655) | No Diff in *Nipbl*+/- (11,947) |
| *Nipbl*+/- Projected onto WT **(Forward Projection)** | Up in *Nipbl*+/- (678) | 642 **(94.0%)** | 6 (0.2%) | 30 (0.3%) |
| | Down in *Nipbl*+/- (3,625) | 0 (0%) | 3,463 **(94.7%)** | 162 (1.4%) |
| | No Diff in *Nipbl*+/- (11,982) | 41 (6.0%) | 186 (5.1%) | 11,755 **(98.4%)** |

**E**

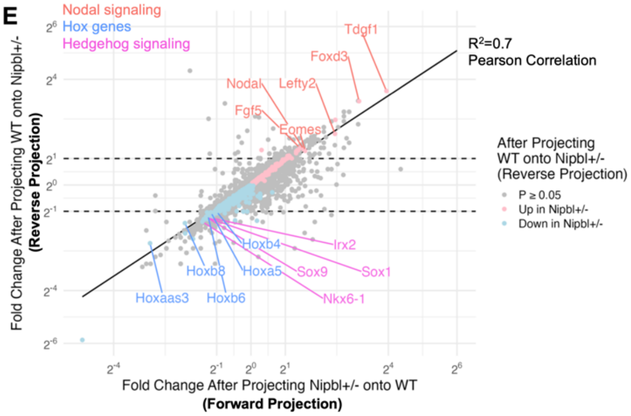

### End

**C**

| | | WT Projected onto *Nipbl*+/- **(Reverse Projection)** | | |
|---|---|---|---|---|
| | | Up in *Nipbl*+/- (488) | Down in *Nipbl*+/- (1,323) | No Diff in *Nipbl*+/- (14,027) |
| *Nipbl*+/- Projected onto WT **(Forward Projection)** | Up in *Nipbl*+/- (412) | **369 (75.6%)** | 1 (0.1%) | 42 (0.3%) |
| | Down in *Nipbl*+/- (1,116) | 3 (0.6%) | **1,019 (77.0%)** | 94 (0.7%) |
| | No Diff in *Nipbl*+/- (14,310) | 116 (23.8%) | 303 (22.9%) | **13,891 (99.0%)** |

**F**

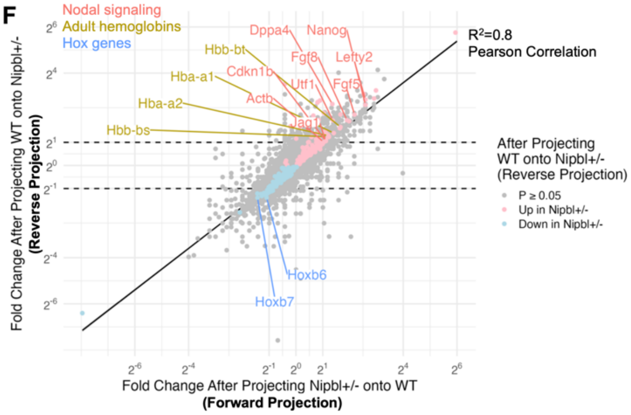

**Fig. S14. Differentially expressed genes between germ layers of WT and *Nipbl*$^{+/-}$ embryos, determined following projection of WT cells onto *Nipbl*$^{+/-}$ germ layers.**

Percentage of differentially expressed genes ($Q < 0.05$, Mann-Whitney U Test) in (**A**) mesoderm, (**B**) ectoderm, and (**C**) endoderm of LB-stage *Nipbl*$^{+/-}$ embryos resulting from the projection of WT cells onto *Nipbl*$^{+/-}$ germ layers (reverse projection) matching those differentially expressed in the germ layers of *Nipbl*$^{+/-}$ embryos resulting from the projection of *Nipbl*$^{+/-}$ cells onto WT germ layers (forward projection). Fold change in expression of differentially expressed genes ($Q < 0.05$, Mann-Whitney U Test) in the (**D**) mesoderm, (**E**) ectoderm, and (**F**) endoderm of LB-stage *Nipbl*$^{+/-}$ embryos following projection of WT cells onto *Nipbl*$^{+/-}$ germ layers (reverse projection) versus the fold change in expression of the same genes in the germ layers of WT embryos following projection of *Nipbl*$^{+/-}$ cells onto WT germ layers (forward projection).

**DEGs in CC-stage *Nipbl*$^{+/-}$ Embryos that are also DEGs in E9.5 *Nanog* Dox+ Embryos** (cf. Ref #*113*)

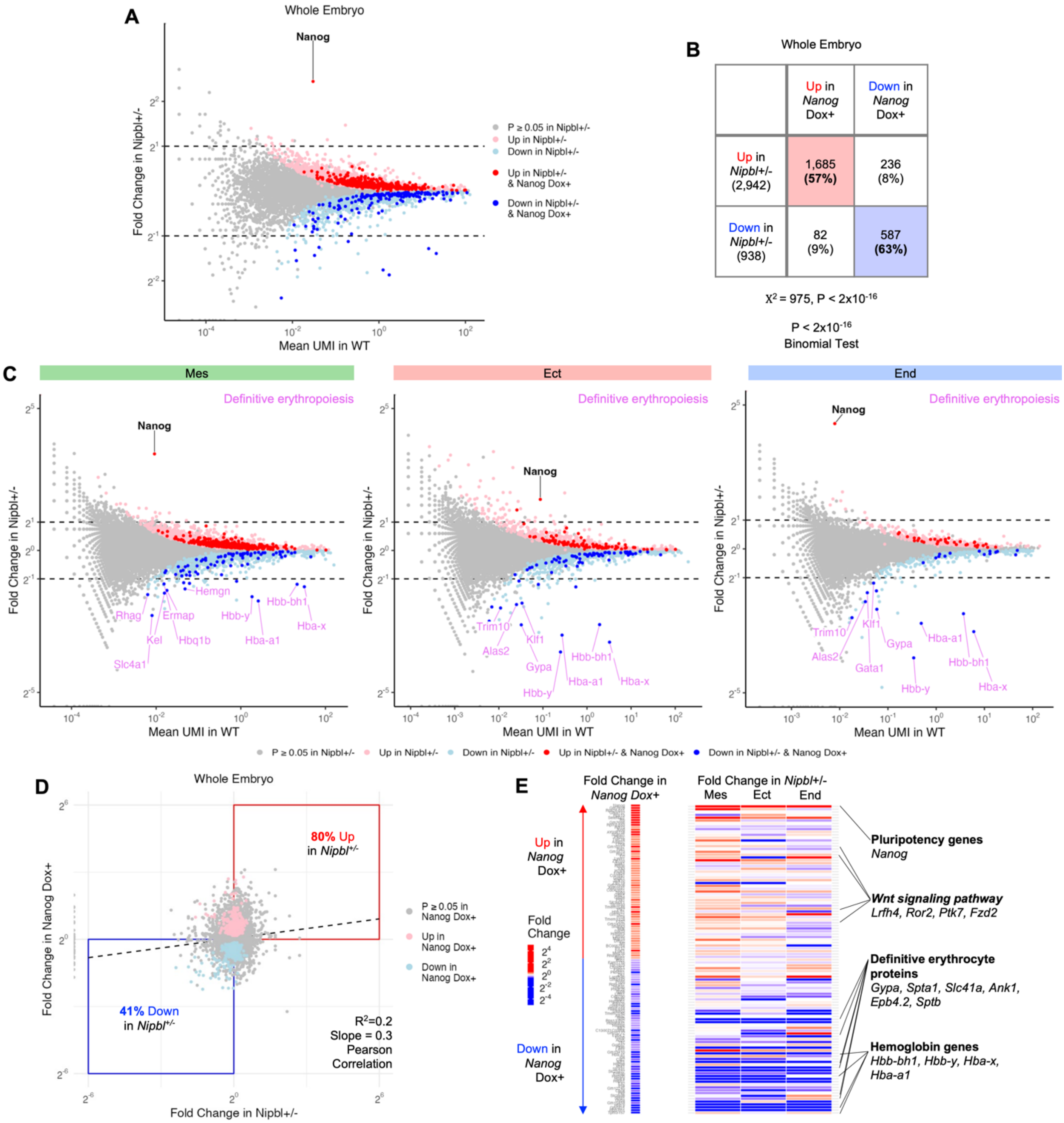

**Fig. S15. CC-stage *Nipbl*$^{+/-}$ mice replicate the gene expression changes of *Nanog* overexpression.**

(**A**) Fold change in expression of genes in CC-stage *Nipbl*$^{+/-}$ embryos that were differentially expressed ($Q < 0.05$, Mann-Whitney U Test) from WT embryos along with their average expression in WT embryos. Genes that are were also differentially expressed ($Q < 0.05$, T-Test) in E9.5 mouse embryos transgenic for doxycycline inducible expression of *Nanog* (*Nanog* Dox+) from that of WT embryos (*Nanog* Dox-) from *(113)* are colored dark red (upregulated) and dark blue (downregulated). (**B**) Percentage of genes in CC-stage *Nipbl*$^{+/-}$ embryos that were differentially expressed ($Q < 0.05$, Mann-Whitney U Test) from WT embryos, and that were also differentially expressed in the same direction in E9.5 *Nanog* Dox+ embryos. At CC-stage, 57% of overexpressed and 63% of underexpressed genes in *Nipbl*$^{+/-}$ embryos were also overexpressed or underexpressed, respectively, in E9.5 *Nanog* Dox+ embryos. (**C**) Fold change in expression of genes in the germ layers of CC-stage *Nipbl*$^{+/-}$ embryos that were differentially expressed ($Q <$ 0.05, Mann-Whitney U Test) from WT embryos along with their average expression in WT embryos. Genes that are were also differentially expressed ($Q < 0.05$, T-Test) in E9.5 *Nanog* Dox+ embryos are colored dark red and dark blue. At CC-stage, all germ layers of *Nipbl*$^{+/-}$ embryos showed downregulation of genes associated with definitive erythropoiesis, encompassing adult hemoglobins including *Hbb-y, Hba-a1,* and *Hba-x* (data S41, data S42, and data S43). (**D**) Fold change in expression of genes in CC-stage *Nipbl*$^{+/-}$ embryos versus their fold change in E9.5 *Nanog* Dox+ embryos. Genes differentially expressed ($Q < 0.05$, T-Test) in E9.5 *Nanog* Dox+ embryos are colored dark red and dark blue. At CC-stage, the slope of the regression line comparing misexpression in *Nipbl*$^{+/-}$ and E9.5 *Nanog* Dox+ embryos was found to be 0.3 (panel D). Additionally, at CC-stage, *Nipbl*$^{+/-}$ embryos upregulated 80% of the same genes that were overexpressed in E7.5 *Nanog* Dox+ embryos and downregulated 41% of the same genes underexpressed in E7.5 *Nanog* Dox+ embryos. (**E**) Heatmap of fold change in expression in germ layers of CC-stage *Nipbl*$^{+/-}$ embryos from that of WT embryos of the most differentially expressed genes from E9.5 *Nanog* Dox+ embryos from that of Nanog Dox- embryos (lowest *Q*-values from T-Test) from *(113)*. Genes are ordered from top to middle (overexpressed in *Nanog* Dox+ embryos) and bottom to middle (underexpressed in *Nanog* Dox+ embryos) by lowest to higher *Q*-value. Underexpressed genes in *Nanog* Dox+ embryos were strongly mirrored in the germ layers of CC-stage *Nipbl*$^{+/-}$ embryos (panel E). In CC-stage *Nipbl*$^{+/-}$ embryos, only a few overexpressed genes observed in *Nanog* Dox+ embryos were replicated, the most prominent of which was *Nanog* itself.

**Table S1. Number of cells, transcripts, and genes captured per LB- and CC-stage WT and *Nipbl*$^{+/-}$ embryos.**

scRNAseq of LB- and CC-stage WT and *Nipbl*$^{+/-}$ embryos captured thousands of cells per embryo, thousands of transcripts per cell, and detected thousands of genes per cell. Table shows stage of each embryo, genotype of each embryo, number of cells captured per embryo, median number of UMIs captured per cell, and median number of genes detected per cell.

| Stage | Genotype | Sample | Cells | Median UMIs/Cell | Median Genes/Cell |
|---|---|---|---|---|---|
| LB | WT | LB1 | 2,445 | 17,998 | 4,242 |
| LB | WT | LB2 | 1,247 | 17,011 | 3,947 |
| LB | WT | LB3 | 2,537 | 16,476 | 3,830 |
| LB | WT | LB4 | 7,119 | 18,624 | 3,940 |
| LB | WT | LB5 | 2,321 | 18,413 | 4,076 |
| LB | WT | LB1-LB5 | 15,669 | 18,078 | 4,001 |
| LB | *Nipbl*$^{+/-}$ | LB6 | 1,700 | 9,953.5 | 2,997 |
| LB | *Nipbl*$^{+/-}$ | LB7 | 1,411 | 18,111 | 4,110 |
| LB | *Nipbl*$^{+/-}$ | LB8 | 4,218 | 11,875 | 3,168 |
| LB | *Nipbl*$^{+/-}$ | LB9 | 3,969 | 15,752 | 3,702 |
| LB | *Nipbl*$^{+/-}$ | LB10 | 3,431 | 17,123 | 3,847 |
| LB | *Nipbl*$^{+/-}$ | LB11 | 4,840 | 18,910.5 | 4,000.5 |
| LB | *Nipbl*$^{+/-}$ | LB6-LB11 | 19,569 | 16,424 | 3,780 |
| LB | *WT & Nipbl*$^{+/-}$ | LB1-LB11 | 35,238 | 17,300 | 3,894 |
| CC | WT | CC1 | 1858 | 15,682 | 3,796 |
| CC | WT | CC2 | 3537 | 14,967 | 3,711 |
| CC | WT | CC3 | 4550 | 13,788 | 3,609 |
| CC | WT | CC4 | 5202 | 14,782 | 3,549 |
| CC | WT | CC5 | 7448 | 14,882 | 3,453 |
| CC | WT | CC6 | 7759 | 14,547 | 3,438 |
| CC | WT | CC7 | 4767 | 15,386 | 3,716 |
| CC | WT | CC8 | 6573 | 15,797 | 3,652 |
| CC | WT | CC1-CC8 | 41,694 | 14,916 | 3,580 |
| CC | *Nipbl*$^{+/-}$ | CC9 | 2104 | 15,861 | 3,837 |
| CC | *Nipbl*$^{+/-}$ | CC10 | 2197 | 14,968 | 3,643 |
| CC | *Nipbl*$^{+/-}$ | CC11 | 669 | 16,370 | 3,994 |
| CC | *Nipbl*$^{+/-}$ | CC12 | 1614 | 15,140 | 3,750 |
| CC | *Nipbl*$^{+/-}$ | CC13 | 2025 | 15,525 | 3,797 |
| CC | *Nipbl*$^{+/-}$ | CC14 | 3683 | 13,302 | 3,493 |
| CC | *Nipbl*$^{+/-}$ | CC15 | 4919 | 14,986 | 3,656 |
| CC | *Nipbl*$^{+/-}$ | CC16 | 8236 | 15,888 | 3,699 |
| CC | *Nipbl*$^{+/-}$ | CC9-16 | 25,447 | 15,122 | 3,677 |
| CC | *WT & Nipbl*$^{+/-}$ | CC1-CC16 | 67,141 | 14,997 | 3,618 |
| LB & CC | *WT & Nipbl*$^{+/-}$ | LB1-LB11 & CC1-CC16 | 102,379 | 15,429 | 3,676 |

### Table S2. Differential overexpression of markers of germ layer identity by cell populations in LB- and CC-stage embryos.

| Stage | Cell Population | Markers | Germ Layer |
|---|---|---|---|
| LB | 1 | *Dnmt3b, Utf1 (31), Pou3f1 (33), Eras, Sox2 (32), Epcam* | Ectoderm |
| LB | 2 | *Gbx2, Pou5f1, Cdx2, Cdkn1a* | Ectoderm |
| LB | 3 | *Tfap2a, Gata3, Tfap2c* | Ectoderm |
| LB | 4 | *T (34), Mixl1, Wnt3a, Nkx1-2, Pou5f1, Nkx2-1* | Mesoderm |
| LB | 5 | *Mesp1, Hand1 (35), Foxf1, Fn1, Snai1, Isl1, Bmp4, Vim (36)* | Mesoderm |
| LB | 6 | *Hoxb1, Tbx6, Dll3, Dll1, Lef1, Frzb* | Mesoderm |
| LB | 7 | *Dppa3, Psmb8, Ifitm3, Tapbp, Irf1* | Mesoderm |
| LB | 8 | *Smarcd3, Myl7, Gata6, Cdkn1c, Mef2c, Meis1* | Mesoderm |
| LB | 9 | *Foxc2, Twist1 (37), Fn1, Hmga2* | Mesoderm |
| LB | 10 | *Foxc2, Twist1 (37), Pitx2, Foxc1, Cd24a, Hmga2, Prrx2 (38)* | Mesoderm |
| LB | 11 | *Hand1 (35), Bmp4, Foxf1, Msx2* | Mesoderm |
| LB | 12 | *Hand1 (35), Pitx1, Bambi, Foxf1, Bmp4* | Mesoderm |
| LB | 13 | *Cubn, Apob, Amn, Apom, Apoa4, Dab2, Pla2g12b* | Endoderm |
| LB | 14 | *Cldn6, Krt18, Krt8, Foxa2, Sox17, Cer1, Cldn7* | Endoderm |
| LB | 15 | *Tex19.1, Dppa4, Zfp4* | Ectoderm |
| LB | 16 | *Tal1, Kdr, Lmo2, Etv2* | Mesoderm |
| LB | 17 | *Gata1, Tal1, Lmo2, Hbb-bh1, Gfi1b* | Mesoderm |
| LB | 18 | *1110017D15Rik, Fam183b, Tekt1, Dynlrb2* | Mesoderm |
| CC | 1 | *Lef1, Tbx6, Dll3, Hes7, Hoxb1, Dll1, Cdx1* | Mesoderm |
| CC | 2 | *Hoxd9, Lef1, Msx1, Hoxb6, Hoxb8, Nkd1, Msx2, Hoxb9* | Mesoderm |
| CC | 3 | *Mdk, Vim (36), Hand2, Arg1, Fn1, Nrp1, Pdgfra, Ppp3ca* | Mesoderm |
| CC | 4 | *Dppa3, Psmb8, Ifitm3, Gdf3, Psme1, Irf1* | Mesoderm |
| CC | 5 | *Twist1 (37), Prrx2 (38), Foxc2, Foxd1, Prrx1* | Mesoderm |
| CC | 6 | *Meis1, Gata6, Isl1, Arg1, Mef2c, Meis2, Smarcd3, Hand2, Nr2f2, Irx3* | Mesoderm |
| CC | 7 | *Foxc2, Foxc1, Prrx2 (38), Twist1 (37)* | Mesoderm |
| CC | 8 | *Arg1, Pkm, Tubb5, Trim28, Eef1a1, Sox2 (32), Actb, Igf2bp1, Pou3f1 (33)* | Ectoderm |
| CC | 9 | *Acta2, Krt8, Krt18, Tagln* | Mesoderm |
| CC | 10 | *Phlda2, Cdkn1c, Peg10, Slc38a4* | Mesoderm |
| CC | 11 | *Etv2, Egfl7, Ramp2, Kdr, Fli1, Hhex, Tal1, Lmo2, Flt1, Rasip1, Esam* | Mesoderm |
| CC | 12 | *Myl7, Actc1, Tnnt2, Myl4* | Mesoderm |
| CC | 13 | *Pou3f1 (33), Sox2 (32), En1, Pax2, Dnmt3b* | Ectoderm |
| CC | 14 | *Hoxa1, Pax6, Sox2 (32), Meis2, Sox3* | Ectoderm |
| CC | 15 | *Six3, Hesx1, Pou3f1 (33), Otx2, Sox2 (32)* | Ectoderm |
| CC | 16 | *Pax3, Sox9, Foxd3* | Ectoderm |
| CC | 17 | *Ptch1, Foxa2, Sox2 (32), Nkx6-1, Pou3f1 (33), Sfrp1* | Ectoderm |
| CC | 18 | *Cdx2, Nr6a1, Pou5f1, Ccnd1* | Mesoderm |
| CC | 19 | *Fst, Fgf8, T (34), Etv4* | Mesoderm |
| CC | 20 | *Msx1, Cdx4, Cdx2, Fgf8, Etv4, T (34), Evx1, Klf5, Nkx1-2* | Mesoderm |
| CC | 21 | *Epcam, Cldn7, Rab25, Cldn6, Krt18, Krt8* | Ectoderm |
| CC | 22 | *Krt18, Epcam, Krt8* | Endoderm |
| CC | 23 | *Shh, T (34), Nog, Foxa2* | Mesoderm |
| CC | 24 | *Apob, Amn, Afp, Apom, Rbp4, Ttr (39), Apoa4, Apoa1, Cubn* | Endoderm |
| CC | 25 | *Ttr (39), Rbp4, Apoa1, Apom* | Endoderm |
| CC | 26 | *Hba-x, Hbb-bh1, Hba-a1, Hba-a2* | Mesoderm |

### Table S3. Differential overexpression of markers of biological identity by endodermal cell populations in LB- and CC-stage embryos.

| Stage | Cell Population | Markers | Biological Identity |
|---|---|---|---|
| LB | 13 | *Hnf4a (42), Tbx3 (43), S100a1, Rbp4, Cubn, Mt1, Slc2a2, Apoa2, Ctsb, Aldob, Apom, Lrp2, Cltc, Aprt* | Extraembryonic endoderm |
| LB | 14 | *Sox4 (40) Mdk, Tead2, Cer1, Ccnd2, Dkk1, Sfrp1, Foxa2 (41), Bambi, Fzd2, Ccnd1* | Definitive endoderm progenitor |
| CC | 22 | *Krt18, Slc2a1, Cd24a, Epcam, Ptma, Hmgn1, Tpt1, Rps2, Bsg, Rps23, Cfl1, Ppia,* | Definitive endoderm progenitor |
| CC | 24 | *Esx1 (44), Ldha, Fam162a, Eno1, Slc16a3, Pgk1, Tubb5, Pkm, Aldoa, Hspa8* | Visceral endoderm |
| CC | 25 | *Rbp4, Apoa1, Ttr, Apom* | Extraembryonic endoderm |

**Table S4. Differential overexpression of markers of biological identity by ectodermal cell populations in LB- and CC-stage embryos.**

| Stage | Cell Population | Markers | Biological Identity |
|---|---|---|---|
| LB | 1 | *Dnmt3b, Pou3f1 (33), Otx2 (45), Utf1, Eras, Irx3, Hesx1, Sox2* | Ectoderm progenitor |
| LB | 2 | *Cdx1 (46), Nkx1-2 (47), Cdx2, Fst, Gbx2, Pou5f1, Cdkn1a, Fgf8* | Neural ectoderm progenitor |
| LB | 3 | *Id1 (45) Msx2 (45), Msx1 (45), Tfap2a* | Surface ectoderm |
| LB | 15 | *Elf5 (49), Dppa4, Tex19.1, Grem2, Bmp8b, Sox7, Tead4 (50), Esrrb, Dppa2, Gata6, Nog, Zfp42, Sox17* | Extraembryonic ectoderm |
| CC | 8 | *Sox2 (60), Ezr (61), Arg1, Fn1, Twist1, Foxc2, Prrx2, Hmga2* | Spinal neural plate |
| CC | 13 | *En1 (54), Pax2 (55), Pou3f1, Zic3, Zeb2, Wnt1, Sox2, Pax5, Sox3* | Mesencephalic neural plate |
| CC | 14 | *Hoxa1 (56), Hes3 (57), Mdk, Meis2, Tshz1, Pax6, Pou5f1, Ptn* | Rhombencephalic neural plate |
| CC | 15 | *Six3 (52), Hesx1 (53), Pou3f1, Otx2, Lhx5, Fezf1, Rax* | Prosencephalic neural plate |
| CC | 16 | *Pax3 (62), Foxd3 (63), Sox9* | Neural crest |
| CC | 17 | *Ptch1, Foxa2 (58), Nkx2-9 (59), Nkx6-1, Cfc1, Sox2, Gli1, Lefty1, Ccnd1, Sfrp1* | Midline neural plate |
| CC | 21 | *Krt8, Krt18, Epcam* | Surface ectoderm |

### Table S5. Differential overexpression of markers of biological identity by mesodermal cell populations in LB- and CC-stage embryos.

| Stage | Cell Population | Markers | Biological Identity |
|---|---|---|---|
| LB | 4 | *Pou5f1, T (34), Nkx1-2 (47), Epcam, Cdh1, Wnt3a, Mixl1, Cdx2, Mgst1, Dnmt3b, Cdkn1a* | Neuromesodermal progenitor |
| LB | 5 | *Mesp1 (64), Hand1, Isl1, Cdx2, Evx1* | *Mesp1*-expressing mesoderm progenitor |
| LB | 6 | *Tbx6 (67), Meis2 (68), Hoxb1, Dll1, Dll3, Lef1, Frzb* | Paraxial mesoderm progenitor |
| LB | 7 | *Dppa3, Ifitm3, Psmb8, Tfap2c (77), Msx1 (78), Psme2, Psme1, Irf1, Tapbp* | Primordial germ cell |
| LB | 8 | *Id2 (65), Smarcd3, Myl7, Cdkn1c, Mef2c (66), Gata6* | Second heart field |
| LB | 9 | *Foxc2 (69), Tcf15 (70), Twist1, Cd24a, Tcf7l1, Lef1* | Trunk paraxial mesoderm |
| LB | 10 | *Alx1 (70), Tcf15 (70), Cd24a, Pitx2, Twist1, Foxc2, Foxc1, Prrx2, Six1, Bmp7, Jag1* | Head paraxial mesoderm |
| LB | 11 | *Hand1 (35), Bmp4, Msx2, Foxf1* | First heart field 2 |
| LB | 12 | *Pitx1, Hand1 (35), Bambi, Krt8, Msx1, Krt18, Foxf1, Bmp4, Col1a1, Msx2, Cdx2* | First heart field 1 |
| LB | 16 | *Kdr, Tal1 (72), Lmo2, Etv2 (73)* | Hemangioblast |
| LB | 17 | *Gata1 (72), Hbb-bh1, Tal1 (72), Runx1* | Primitive erythrocyte |
| LB | 18 | *Noto (75), Foxj1 (76), 1110017D15Rik, Tekt1, Pifo, Enkur, Fam183b* | Primitive node |
| CC | 1 | *Tbx6, Dll3, Lef1, Hes7, Dll1, Hoxb1, Cdx1* | Trunk paraxial mesoderm |
| CC | 2 | *Mesp1 (64), Hand1, Msx1, Lef1, Msx2, Cdx2, Nkd1, Sall4, Tbx3* | *Mesp1*-expressing mesoderm progenitor 1 |
| CC | 3 | *Mesp1 (64), Meis1, Meis2, Hoxa1, Tshz1, Hoxa5, Foxp1, Pbx3* | *Mesp1*-expressing mesoderm progenitor 2 |
| CC | 4 | *Dppa3, Psmb8, Ifitm3, Irf1, Dnd1, Gdf3, Psme1, Parp12, Psme2, Nanos3, Ifitm2, Tapbp* | Primordial germ cell |
| CC | 5 | *Twist1, Prrx2, Foxc2, Foxd1, Prrx1* | Head paraxial mesoderm |
| CC | 6 | *Meis1, Gata6, Isl1, Irx3, Meis2, Irx5* | Second heart field |
| CC | 7 | *Tcf15 (70), Meox1 (82), Foxc2, Ptn, Pax3, Foxc1, Cadm1, Mdk, H3f3a, Epb41l3, Foxd1, Fbn2, Six1, Ncam1, Notch1, Prrx2, Efnb2, Pdgfra, Twist1, Zic2* | Presomitic mesoderm |
| CC | 9 | *Krt18, Krt8, Tagln, Hand1, Acta2, Sparc, Cnn2, Vim, Bmp4, Csrp2, Nrp1, Serpinh1, Myl6* | First heart field 2 |
| CC | 10 | *Hand1, Bambi, Bmp4, Twist1* | First heart field 1 |
| CC | 11 | *Etv2, Ramp2, Egfl7, Kdr, Fli1, Hhex, Rasip1, Flt1, Tal1, Esam, Lmo2* | Hemangioblast |
| CC | 12 | *Mef2c (80), Nkx2-5 (81), Myl7, Actc1, Tnnt2, Myl4* | Cardiomyocyte |
| CC | 18 | *Cdx2, Sox2, Nr6a1, Pou5f1* | Neuromesodermal progenitor |
| CC | 19 | *T, Fgf8, Fst, Etv4, Fgf17* | Paraxial mesoderm progenitor |
| CC | 20 | *Tlx2 (79), Msx1, Cdx4, Cdx2, Evx1os, Etv4, Fgf8, T (34)* | Lateral plate mesoderm progenitor |
| CC | 23 | *Shh, Foxa2, Nog, Krt18* | Primitive node |
| CC | 26 | *Hba-x, Hbb-bh1, Hba-a1, Hba-a2* | Primitive erythrocyte |

### Table S6. Differential overexpression of markers of germ layer identity by cell populations in LB-stage *Nipbl*$^{+/-}$ embryos.

| Stage | Cell Population | Markers | Germ Layer |
|---|---|---|---|
| LB | 1 | *Dnmt3b, Utf1 (31)* | Ectoderm |
| LB | 2 | *Dlx5, Tfap2a* | Ectoderm |
| LB | 3 | *Pou5f1, Dnmt3b, Hes3* | Ectoderm |
| LB | 4 | *Cdx1, Mixl1, Pou5f1* | Mesoderm |
| LB | 5 | *Msx1, Tfap2a* | Ectoderm |
| LB | 6 | *Foxc2, Lefty2* | Mesoderm |
| LB | 7 | *Mesp1, Rspo3, Isl1, Wnt2* | Mesoderm |
| LB | 8 | *Myl7, Smarc3d, Cdkn1c, Gata6, Phlda2, Mef2c* | Mesoderm |
| LB | 9 | *Tcf15, Foxc2* | Mesoderm |
| LB | 10 | *Hand1 (35), Bmp4* | Mesoderm |
| LB | 11 | *Hoxb6, Hand1 (35), Hoxd9, Pitx1, Foxf1* | Mesoderm |
| LB | 12 | *Mesp1, Mixl1* | Mesoderm |
| LB | 13 | *Dppa3, Dnd1* | Mesoderm |
| LB | 14 | *Hand1 (35), Krt18, Krt8, Foxf1, Bmp4,* | Mesoderm |
| LB | 15 | *Tbx6, Hoxb1* | Mesoderm |
| LB | 16 | *Tal1, Lmo2* | Mesoderm |
| LB | 17 | *Ttr, Apoa1* | Endoderm |
| LB | 18 | *Slc116a1, Emb* | Endoderm |
| LB | 19 | *Pifo, Cfap126* | Mesoderm |
| LB | 20 | *Tex19.1, Dppa4* | Endoderm |
| LB | 21 | *Hs3st1, Srgn* | Endoderm |

### Table S7. Differential overexpression of markers of biological identity by mesodermal cell populations in LB-stage *Nipbl*$^{+/-}$ embryos.

| Stage | Cell Population | Markers | Biological Identity |
|---|---|---|---|
| LB | 4 | *Pou5f1, T (34)* | Neuromesodermal progenitor |
| LB | 6 | *Foxc2 (69), Tcf15 (70)* | Trunk paraxial mesoderm |
| LB | 7 | *Mesp1 (64), Rspo3, Hoxb1, Wnt2, Isl1, Meis2 (68)* | Mesp1-expressing mesoderm progenitor 3 |
| LB | 8 | *Myl7, Smarcd3, Gata6, Cdkn1c, Mef2c (66)* | Second heart field |
| LB | 9 | *Otx2, Cer1* | Head paraxial mesoderm |
| LB | 10 | *Bmp4, Hand1 (35)* | Mesp1-expressing mesoderm progenitor 2 |
| LB | 11 | *Hoxb6, Hoxd9* | First heart field 2 |
| LB | 12 | *Mixl1, Mesp1 (64)* | Mesp1-expressing mesoderm progenitor 1 |
| LB | 13 | *Dppa3, Psmb8, Dnd1, Nanos3, Ifitm3, Prdm14* | Primordial germ cell |
| LB | 14 | *Krt18, Krt8* | First heart field 2 |
| LB | 15 | *Tbx6 (67), Rbp1, Hoxb1, Aldh1a2* | Paraxial mesoderm progenitor |
| LB | 16 | *Tal1 (72), Lmo2* | Hemangioblast |
| LB | 19 | *Pifo, Cfap126, Foxj1 (76)* | Primitive node |

**Table S8. Number of cells, transcripts, and genes captured by scRNAseq of EB- and EHF-stage WT embryos.**

scRNA-seq of EB- and EHF-stage of WT mouse embryos captured thousands of cells per embryo, thousands of transcripts per cell, and detected thousands of genes per cell. Table shows genotype of each embryo, number of cells captured per embryo, median number of UMIs captured per cell, and median number of genes detected per cell.

| Embryo(s) | Genotype | Cells | Median UMIs/Cell | Median Genes/Cell |
|---|---|---|---|---|
| EB1 | WT | 4109 | 5693 | 2064 |
| EB2 | WT | 1185 | 5943 | 2120 |
| EB1-EB2 | WT | 5294 | 5740 | 2073.5 |
| EHF1 | WT | 1969 | 11208 | 3271 |
| EHF2 | WT | 3839 | 9587 | 2891 |
| EHF1-EHF2 | WT | 5808 | 10162 | 3030 |

### Table S9. Differential overexpression of markers of germ layer identity by cell populations in EB- and EHF-stage embryos.

| Stage | Cell Population | Markers | Germ Layer |
|---|---|---|---|
| EB | 1 | *Pou5f1, Dnmt3b, Epcam, Utf1, Pkm* | Ectoderm |
| EB | 2 | *Vim, Hand1, Hmga2, Fn1, Actb, Fgf3, Tpm4, Myl7, Cald1* | Mesoderm |
| EB | 3 | *Lhx1, Eomes, Fgf3, Snai1, Fn1, Mixl1* | Mesoderm |
| EB | 4 | *Apom, Apoa1, Dab2, Ttr, Apob, Amn* | Endoderm |
| EB | 5 | *Cer1, Gsc, Lhx1, Sox17, Fgf5, Foxa2* | Endoderm |
| EB | 6 | *Foxj1, 1110017D15Rik, Tekt1, Pifo, Cfap126* | Mesoderm |
| EB | 7 | *Lmo2, Fgf3, Tal1, Runx1, Fli1, Kdr* | Mesoderm |
| EB | 8 | *Elf5, S100a6, Nr0b1, Anxa5, Rhox5, Trap1a, Tex19.1, Hspb1, Tfap2c, Dnmt3l, Dppa4, Cd9, Lgals1, Zfp42* | Ectoderm |
| EHF | 1 | *Dnmt3b, Gapdh, Epcam, Sox2, Pou3f1* | Ectoderm |
| EHF | 2 | *Pou5f1, Gbx2, Sox3, Cdx2* | Ectoderm |
| EHF | 3 | *T, Fst, Nkx1-2, Cdx2, Pou5f1, Wnt3a, Fgf8* | Mesoderm |
| EHF | 4 | *Tacstd2, Wnt6, Wnt4, Lrp2, Lrpap1, Krt18, Krt8, Wt1* | Ectoderm |
| EHF | 5 | *Lef1, Dll1, Pcdh19, Dll3, Tbx6, Cdh2* | Mesoderm |
| EHF | 6 | *Hmga2, Twist1, Prrx2, Foxc2* | Mesoderm |
| EHF | 7 | *Hand1, Krt18, Krt8, Foxf1, Tagln, Vim, Serpinh1, Ppic, Cnn2, Tpm1, Nrp1, Acta2, Bmp4, Myl6* | Mesoderm |
| EHF | 8 | *Dppa3, Ifitm3, Gdf3, Dnd1, Parp9, Tfap2c, Klf2, Parp12, Prdm1* | Mesoderm |
| EHF | 9 | *Amn, Apom, Apoa1, Cubn, Apob, Ttr* | Endoderm |
| EHF | 10 | *Krt8, Krt18, Sox17, Foxa1, Foxa2* | Endoderm |
| EHF | 11 | *Foxj1, 1110017D15Rik, Dynlrb2, Cfap52, Fam183b, Cfap126* | Mesoderm |
| EHF | 12 | *Lmo2, Gata1, Hbb-bh1, Runx1* | Mesoderm |
| EHF | 13 | *Kdr, Fli1, Etv2, Lmo2* | Mesoderm |

**Table S10. Expression of *Nanog* in LB- and CC-stage embryos.**

Genotype = genotype of embryo, Embryo = embryo identifier, Cell_Type = cell type in embryo, Nanog = normalized transcript counts for *Nanog* in cell type.

| Stage | Genotype | Embryo | *Nanog* |
|---|---|---|---|
| Late bud | Wildtype | LB1 | 0.05051302 |
| Late bud | Wildtype | LB2 | 0.11359867 |
| Late bud | Wildtype | LB3 | 0.08 |
| Late bud | Wildtype | LB4 | 0.10866373 |
| Late bud | Wildtype | LB5 | 0.07430341 |
| Late bud | *Nipbl*$^{+/-}$ | LB6 | 0.3121175 |
| Late bud | *Nipbl*$^{+/-}$ | LB7 | 0.20864662 |
| Late bud | *Nipbl*$^{+/-}$ | LB8 | 0.24953096 |
| Late bud | *Nipbl*$^{+/-}$ | LB9 | 0.18756219 |
| Late bud | *Nipbl*$^{+/-}$ | LB10 | 0.16984127 |
| Late bud | *Nipbl*$^{+/-}$ | LB11 | 0.19027661 |
| Cardiac crescent | Wildtype | CC1 | 0.00668338 |
| Cardiac crescent | Wildtype | CC2 | 0.01766304 |
| Cardiac crescent | Wildtype | CC3 | 0.01175446 |
| Cardiac crescent | Wildtype | CC4 | 0.00583771 |
| Cardiac crescent | Wildtype | CC5 | 0.01085305 |
| Cardiac crescent | Wildtype | CC6 | 0.00340408 |
| Cardiac crescent | Wildtype | CC7 | 0.010921 |
| Cardiac crescent | Wildtype | CC8 | 0.01019946 |
| Cardiac crescent | *Nipbl*$^{+/-}$ | CC9 | 0.15524194 |
| Cardiac crescent | *Nipbl*$^{+/-}$ | CC10 | 0.12248629 |
| Cardiac crescent | *Nipbl*$^{+/-}$ | CC11 | 0.08695652 |
| Cardiac crescent | *Nipbl*$^{+/-}$ | CC12 | 0.12486993 |
| Cardiac crescent | *Nipbl*$^{+/-}$ | CC13 | 0.11223629 |
| Cardiac crescent | *Nipbl*$^{+/-}$ | CC14 | 0.09711432 |
| Cardiac crescent | *Nipbl*$^{+/-}$ | CC15 | 0.07511408 |
| Cardiac crescent | *Nipbl*$^{+/-}$ | CC16 | 0.07357057 |

### Table S11. Expression of *Pou5f1* in LB- and CC-stage embryos.

Genotype = genotype of embryo, Embryo = embryo identifier, Cell_Type = cell type in embryo, Pou5f1 = normalized transcript counts for *Pou5f1* in cell type.

| Stage | Genotype | Embryo | *Pou5f1* |
|---|---|---|---|
| Late bud | Wildtype | LB1 | 8.3875296 |
| Late bud | Wildtype | LB2 | 8.28772803 |
| Late bud | Wildtype | LB3 | 8.98193548 |
| Late bud | Wildtype | LB4 | 10.6762115 |
| Late bud | Wildtype | LB5 | 7.02786378 |
| Late bud | *Nipbl*$^{+/-}$ | LB6 | 7.82741738 |
| Late bud | *Nipbl*$^{+/-}$ | LB7 | 12.1409774 |
| Late bud | *Nipbl*$^{+/-}$ | LB8 | 8.35897436 |
| Late bud | *Nipbl*$^{+/-}$ | LB9 | 10.3502488 |
| Late bud | *Nipbl*$^{+/-}$ | LB10 | 6.68201058 |
| Late bud | *Nipbl*$^{+/-}$ | LB11 | 7.03855826 |
| Cardiac crescent | Wildtype | CC1 | 0.66165414 |
| Cardiac crescent | Wildtype | CC2 | 1.05661232 |
| Cardiac crescent | Wildtype | CC3 | 0.48193296 |
| Cardiac crescent | Wildtype | CC4 | 0.83712785 |
| Cardiac crescent | Wildtype | CC5 | 0.76383764 |
| Cardiac crescent | Wildtype | CC6 | 0.43211854 |
| Cardiac crescent | Wildtype | CC7 | 1.3815071 |
| Cardiac crescent | Wildtype | CC8 | 0.96033545 |
| Cardiac crescent | *Nipbl*$^{+/-}$ | CC9 | 0.92271505 |
| Cardiac crescent | *Nipbl*$^{+/-}$ | CC10 | 0.74710542 |
| Cardiac crescent | *Nipbl*$^{+/-}$ | CC11 | 4.35652174 |
| Cardiac crescent | *Nipbl*$^{+/-}$ | CC12 | 1.9760666 |
| Cardiac crescent | *Nipbl*$^{+/-}$ | CC13 | 2.9164557 |
| Cardiac crescent | *Nipbl*$^{+/-}$ | CC14 | 0.91287458 |
| Cardiac crescent | *Nipbl*$^{+/-}$ | CC15 | 1.2965953 |
| Cardiac crescent | *Nipbl*$^{+/-}$ | CC16 | 1.39984006 |

**Data S1. Differentially expressed genes among clusters of LB-stage WT embryos.**

Differential gene expression analysis was performed between clusters of LB-stage WT embryos using the Mann-Whitney U test. P-values were adjusted for multiple hypotheses by Bonferroni Correction. Cluster = cluster number, Gene = gene, Pct_Expr_Cluster = percentage of cells in cluster expressing indicated gene, Pct_Expr_Other = percentage of cells in all other clusters expressing indicated gene, Avg_FC = average fold change in expression of indicated gene in cluster versus all other clusters, P_Val = p-value from Mann-Whitney U test, Bon_P_Val = Bonferroni corrected p-value, Germ_Layer = germ layer assigned to cluster.

**Data S2. Differentially expressed genes among clusters of CC-stage WT embryos.**

Differential gene expression analysis was performed between clusters of CC-stage WT embryos using the Mann-Whitney U test. P-values were adjusted for multiple hypotheses by Bonferroni Correction. Cluster = cluster number, Gene = gene, Pct_Expr_Cluster = percentage of cells in cluster expressing indicated gene, Pct_Expr_Other = percentage of cells in all other clusters expressing indicated gene, Avg_FC = average fold change in expression of indicated gene in cluster versus all other clusters, P_Val = p-value from Mann-Whitney U test, Bon_P_Val = Bonferroni corrected p-value, Germ_Layer = germ layer assigned to cluster.

**Data S3. Differentially expressed genes among endodermal clusters of LB-stage WT embryos.**

Differential gene expression analysis was performed between endodermal clusters of LB-stage WT embryos using the Mann-Whitney U test. P-values were adjusted for multiple hypotheses by Bonferroni Correction. Cluster = cluster number, Gene = gene, Pct_Expr_Cluster = percentage of cells in cluster expressing indicated gene, Pct_Expr_Other = percentage of cells in all other clusters expressing indicated gene, Avg_FC = average fold change in expression of indicated gene in cluster versus all other clusters, P_Val = p-value from Mann-Whitney U test, Bon_P_Val = Bonferroni corrected p-value, Cell_Type = cell type assigned to cluster, Transcription_Factor = Y, predicted by Animal Transcription Factor Database to be a transcription factor.

**Data S4. Differentially expressed genes among endodermal clusters of CC-stage WT embryos.**

Differential gene expression analysis was performed between endodermal clusters of CC-stage WT embryos using the Mann-Whitney U test. P-values were adjusted for multiple hypotheses by Bonferroni Correction. Cluster = cluster number, Gene = gene, Pct_Expr_Cluster = percentage of cells in cluster expressing indicated gene, Pct_Expr_Other = percentage of cells in all other clusters expressing indicated gene, Avg_FC = average fold change in expression of indicated gene in cluster versus all other clusters, P_Val = p-value from Mann-Whitney U test, Bon_P_Val = Bonferroni corrected p-value, Cell_Type = cell type assigned to cluster, Transcription_Factor = Y, predicted by Animal Transcription Factor Database to be a transcription factor.

**Data S5. Differentially expressed genes among ectodermal clusters of LB-stage WT embryos.**

Differential gene expression analysis was performed between ectodermal clusters of LB-stage WT embryos using the Mann-Whitney U test. P-values were adjusted for multiple hypotheses by Bonferroni Correction. Cluster = cluster number, Gene = gene, Pct_Expr_Cluster = percentage of cells in cluster expressing indicated gene, Pct_Expr_Other = percentage of cells in all other clusters expressing indicated gene, Avg_FC = average fold change in expression of indicated gene in cluster versus all other clusters, P_Val = p-value from Mann-Whitney U test, Bon_P_Val = Bonferroni corrected p-value, Cell_Type = cell type assigned to cluster, Transcription_Factor = Y, predicted by Animal Transcription Factor Database to be a transcription factor.

**Data S6. Differentially expressed genes among ectodermal clusters of CC-stage WT embryos.**

Differential gene expression analysis was performed between ectodermal clusters of CC-stage WT embryos using the Mann-Whitney U test. P-values were adjusted for multiple hypotheses by Bonferroni Correction. Cluster = cluster number, Gene = gene, Pct_Expr_Cluster = percentage of cells in cluster expressing indicated gene, Pct_Expr_Other = percentage of cells in all other clusters expressing indicated gene, Avg_FC = average fold change in expression of indicated gene in cluster versus all other clusters, P_Val = p-value from Mann-Whitney U test, Bon_P_Val = Bonferroni corrected p-value, Cell_Type = cell type assigned to cluster, Transcription_Factor = Y, predicted by Animal Transcription Factor Database to be a transcription factor.

**Data S7. Differentially expressed genes among mesodermal clusters of LB-stage WT embryos.**

Differential gene expression analysis was performed between mesodermal clusters of LB-stage WT embryos using the Mann-Whitney U test. P-values were adjusted for multiple hypotheses by Bonferroni Correction. Cluster = cluster number, Gene = gene, Pct_Expr_Cluster = percentage of cells in cluster expressing indicated gene, Pct_Expr_Other = percentage of cells in all other clusters expressing indicated gene, Avg_FC = average fold change in expression of indicated gene in cluster versus all other clusters, P_Val = p-value from Mann-Whitney U test, Bon_P_Val = Bonferroni corrected p-value, Cell_Type = cell type assigned to cluster, Transcription_Factor = Y, predicted by Animal Transcription Factor Database to be a transcription factor.

**Data S8. Differentially expressed genes among mesodermal clusters of CC-stage WT embryos.**

Differential gene expression analysis was performed between mesodermal clusters of CC-stage WT embryos using the Mann-Whitney U test. P-values were adjusted for multiple hypotheses by Bonferroni Correction. Cluster = cluster number, Gene = gene, Pct_Expr_Cluster = percentage of cells in cluster expressing indicated gene, Pct_Expr_Other = percentage of cells in all other clusters expressing indicated gene, Avg_FC = average fold change in expression of indicated gene in cluster versus all other clusters, P_Val = p-value from Mann-Whitney U test, Bon_P_Val = Bonferroni corrected p-value, Cell_Type = cell type assigned to cluster, Transcription_Factor = Y, predicted by Animal Transcription Factor Database to be a transcription factor.

**Data S9. Numbers of cells per germ layer per LB-stage embryo.**

Genotype = genotype of embryo, Embryo = embryo identifier, Germ_Layer = germ layer of embryo, No_Cells = number of cells in germ layer.

**Data S10. Numbers of cells per mesodermal cell population per LB-stage embryo.**

Genotype = genotype of embryo, Embryo = embryo identifier, Cell_Type = cell type of embryo, No_Cells = number of cells in cell type.

**Data S11. Differentially expressed genes among clusters of LB-stage *Nipbl*$^{+/-}$ embryos.**

Differential gene expression analysis was performed between clusters of LB-stage *Nipbl*$^{+/-}$ embryos using the Mann-Whitney U test. P-values were adjusted for multiple hypotheses by Bonferroni Correction. Cluster = cluster number, Gene = gene, Pct_Expr_Cluster = percentage of cells in cluster expressing indicated gene, Pct_Expr_Other = percentage of cells in all other clusters expressing indicated gene, Avg_FC = average fold change in expression of indicated gene in cluster versus all other clusters, P_Val = p-value from Mann-Whitney U test, Bon_P_Val = Bonferroni corrected p-value, Germ_Layer = germ layer assigned to cluster.

**Data S12. Differentially expressed genes among mesodermal clusters of LB-stage *Nipbl*$^{+/-}$ embryos.**

Differential gene expression analysis was performed between mesodermal clusters of LB-stage *Nipbl*$^{+/-}$ embryos using the Mann-Whitney U test. P-values were adjusted for multiple hypotheses by Bonferroni Correction. Cluster = cluster number, Gene = gene, Pct_Expr_Cluster = percentage of cells in cluster expressing indicated gene, Pct_Expr_Other = percentage of cells in all other clusters expressing indicated gene, Avg_FC = average fold change in expression of indicated gene in cluster versus all other clusters, P_Val = p-value from Mann-Whitney U test, Bon_P_Val = Bonferroni corrected p-value, Cell_Type = cell type assigned to cluster.

**Data S13. Numbers of cells per germ layer per LB-stage embryo following projection of WT cells onto *Nipbl*$^{+/-}$ germ layers.**

Genotype = genotype of embryo, Embryo = embryo identifier, Germ_Layer = germ layer of embryo, No_Cells = number of cells in germ layer.

**Data S14. Numbers of cells per mesodermal cell population per LB-stage embryo following projection of WT cells onto *Nipbl*$^{+/-}$ mesodermal cell populations.**

Genotype = genotype of embryo, Embryo = embryo identifier, Cell_Type = cell type of embryo, No_Cells = number of cells in cell type.

**Data S15. Differentially expressed genes among clusters of EB-stage WT embryos.**

Differential gene expression analysis was performed between clusters of EB-stage WT embryos using the Mann-Whitney U test. P-values were adjusted for multiple hypotheses by Bonferroni Correction. Cluster = cluster number, Gene = gene, Pct_Expr_Cluster = percentage of cells in cluster expressing indicated gene, Pct_Expr_Other = percentage of cells in all other clusters expressing indicated gene, Avg_FC = average fold change in expression of indicated gene in cluster versus all other clusters, P_Val = p-value from Mann-Whitney U test, Bon_P_Val = Bonferroni corrected p-value, Germ_Layer = germ layer assigned to cluster.

**Data S16. Differentially expressed genes among clusters of EHF-stage WT embryos.**

Differential gene expression analysis was performed between clusters of EHF-stage WT embryos using the Mann-Whitney U test. P-values were adjusted for multiple hypotheses by Bonferroni Correction. Cluster = cluster number, Gene = gene, Pct_Expr_Cluster = percentage of cells in cluster expressing indicated gene, Pct_Expr_Other = percentage of cells in all other clusters expressing indicated gene, Avg_FC = average fold change in expression of indicated gene in cluster versus all other clusters, P_Val = p-value from Mann-Whitney U test, Bon_P_Val = Bonferroni corrected p-value, Germ_Layer = germ layer assigned to cluster.

**Data S17. Pseudotime values of cells from EB-, LB-, and CC-stage embryos.**

Pseudotime of cells from EB-, LB-, and CC-stage embryos were calculated using URD. Stage = stage of embryo, Genotype = genotype of embryo, Embryo = identifier of embryo, Cell_Barcode = barcode identifying cell, Cluster = cluster number, Germ_Layer = germ layer assigned to cell, Pseudotime = pseudotime value of cell.

**Data S18. RNA velocity vectors of mesoderm cells from LB-stage WT embryos.**

scVelo was used to calculate the RNA velocities of mesoderm cells from LB-stage WT embryos. Embryo = embryo identifier, Cell_Barcode = barcode identifying cell, UMAP_1 = first coordinate in UMAP, UMAP_2 = second coordinate in UMAP, Cluster = cluster number, Cell_Type = cell type assigned to cell, Velocity_1 = first coordinate of RNA velocity vector, Velocity_2 = second coordinate of RNA velocity vector.

**Data S19. RNA velocity vectors of mesoderm cells from LB-stage *Nipbl*$^{+/-}$ embryos.**

scVelo was used to calculate the RNA velocities of mesoderm cells from LB-stage *Nipbl*$^{+/-}$ embryos. Embryo = embryo identifier, Cell_Barcode = barcode identifying cell, UMAP_1 = first coordinate in UMAP, UMAP_2 = second coordinate in UMAP, Cluster = cluster number, Cell_Type = cell type assigned to cell, Velocity_1 = first coordinate of RNA velocity vector, Velocity_2 = second coordinate of RNA velocity vector.

**Data S20. Fate probabilities of mesoderm cells into first heart field, second heart field, and paraxial mesoderm fates from LB-stage WT embryos.**

CellRank was used to calculate the fate probabilities of mesoderm cells into first heart field, second heart field, and paraxial mesoderm fates from LB-stage WT embryos. Embryo = embryo identifier, Cell_Barcode = barcode identifying cell, UMAP_1 = first coordinate in UMAP, UMAP_2 = second coordinate in UMAP, Cluster = cluster number, Cell_Type = cell type assigned to cell, FHF_Prob = first heart field fate probability, SHF_Prob = second heart field fate probability, PM_Prob = paraxial mesoderm fate probability.

**Data S21. Fate probabilities of mesoderm cells into first heart field, second heart field, and paraxial mesoderm fates from LB-stage *Nipbl*$^{+/-}$ embryos.**

CellRank was used to calculate the fate probabilities of mesoderm cells into first heart field, second heart field, and paraxial mesoderm fates from LB-stage *Nipbl*$^{+/-}$ embryos. Embryo = embryo identifier, Cell_Barcode = barcode identifying cell, UMAP_1 = first coordinate in UMAP, UMAP_2 = second coordinate in UMAP, Cluster = cluster number, Cell_Type = cell type assigned to cell, FHF_Prob = first heart field fate probability, SHF_Prob = second heart field fate probability, PM_Prob = paraxial mesoderm fate probability.

**Data S22. Differentially expressed genes from Reactome, Hallmark, and Langemeijer Apoptosis Gene Set between germ layers of LB-stage WT and *Nipbl*$^{+/-}$ embryos.**

Differential gene expression analysis was performed using genes from the Reactome, Hallmark, and Langemeijer Apoptosis Gene Set between the germ layers of LB-stage WT and *Nipbl*$^{+/-}$ embryos using the Mann-Whitney U test. P-values were adjusted for multiple hypotheses by Bonferroni Correction. Gene = gene, Avg_Expr_Wildtype = average expression of indicated gene in Wildtype cells, Avg_Expr_Nipbl+/- = average expression of indicated gene in *Nipbl*$^{+/-}$ cells, SE_Wildtype = standard error of average expression in Wildtype cells, SE_Nipbl+/- = standard error of average expression in *Nipbl*$^{+/-}$ cells, Avg_FC = average fold change in expression of indicated gene in *Nipbl*$^{+/-}$ cells versus WT cells, P_Val = p-value from Mann-Whitney U test, Bon_P_Val = Bonferroni corrected p-value.

**Data S23. Differentially expressed genes from meta-PCNA Gene Set between germ layers of LB-stage WT and *Nipbl*$^{+/-}$ embryos.**

Differential gene expression analysis was performed using genes from the meta-PCNA Gene Set between the germ layers of LB-stage WT and *Nipbl*$^{+/-}$ embryos using the Mann-Whitney U test. P-values were adjusted for multiple hypotheses by Bonferroni Correction. Gene = gene, Avg_Expr_Wildtype = average expression of indicated gene in Wildtype cells, Avg_Expr_Nipbl+/- = average expression of indicated gene in *Nipbl*$^{+/-}$ cells, SE_Wildtype = standard error of average expression in Wildtype cells, SE_Nipbl+/- = standard error of average expression in *Nipbl*$^{+/-}$ cells, Avg_FC = average fold change in expression of indicated gene in *Nipbl*$^{+/-}$ cells versus WT cells, P_Val = p-value from Mann-Whitney U test, Bon_P_Val = Bonferroni corrected p-value.

**Data S24. Cell cycle scores and phases of cells in germ layers of LB-stage WT and *Nipbl*$^{+/-}$ embryos.**

Seurat was used to score and assign cells from all germ layers of LB-stage WT and *Nipbl*$^{+/-}$ embryos into either G1, S, or G2/M phase on the basis of the expression of markers of S and G2/M phase. Expression of these marker sets are considered anticorrelated and when cells express neither, they are considered to be in G1 phase. Dimensional reduction was performed using principal component analysis on S phase and G2/M markers. Genotype = genotype of embryo, Embryo = identifier of embryo, Cell_Barcode = barcode identifying cell, PC_1 = principal component 1, PC_2 = principal component 2, Score_S_Phase = score for S phase markers, Score_G2M_Phase = score for G2/M phase markers, Phase = assigned cell cycle phase.

**Data S25. First heart field drivers and anti-drivers predicted by CellRank.**

Gene = gene, Slope = change in expression of gene over change in absorption probability towards FHF fate in WT cells in Euclidean space, Spearman_Coefficient = correlation coefficient from Spearman's Rank Correlation Test between expression of gene and absorption probability, P_Val = p-value from Spearman's Rank Correlation Test, Ben_Hoch_P_Val = Benjamini-Hochberg corrected p-value, Avg_Expr_WT = average expression of gene in WT cells, Transcription_Factor = Y, predicted by Animal Transcription Factor Database to be a transcription factor.

**Data S26. Paraxial mesoderm drivers and anti-drivers predicted by CellRank.**

Gene = gene, Slope = change in expression of gene over change in absorption probability towards PM fate in WT cells in Euclidean space, Spearman_Coefficient = correlation coefficient from Spearman's Rank Correlation Test between expression of gene and absorption probability, P_Val = p-value from Spearman's Rank Correlation Test, Ben_Hoch_P_Val = Benjamini-Hochberg corrected p-value, Avg_Expr_WT = average expression of gene in WT cells, Transcription_Factor = Y, predicted by Animal Transcription Factor Database to be a transcription factor.

**Data S27. Differentially expressed genes between first heart field lineage of LB-stage WT and *Nipbl*$^{+/-}$ embryos.**

Differential gene expression analysis was performed between FHF lineage of LB-stage WT and *Nipbl*$^{+/-}$ embryos using the Mann-Whitney U test. P-values were adjusted for multiple hypotheses by Bonferroni Correction. Gene = gene, Avg_FC = average fold change in expression of indicated gene in *Nipbl*$^{+/-}$ cells versus WT cells, P_Val = p-value from Mann-Whitney U test, Bon_P_Val = Bonferroni corrected p-value, Transcription_Factor = Y, predicted by Animal Transcription Factor Database to be a transcription factor.

**Data S28. Differentially expressed genes between paraxial mesoderm lineage of LB-stage WT and *Nipbl*$^{+/-}$ embryos.**

Differential gene expression analysis was performed between PM lineage of LB-stage WT and *Nipbl*$^{+/-}$ embryos using the Mann-Whitney U test. P-values were adjusted for multiple hypotheses by Bonferroni Correction. Gene = gene, Avg_FC = average fold change in expression of indicated gene in *Nipbl*$^{+/-}$ cells versus WT cells, P_Val = p-value from Mann-Whitney U test, Bon_P_Val = Bonferroni corrected p-value, Transcription_Factor = Y, predicted by Animal Transcription Factor Database to be a transcription factor.

**Data S29. Differentially expressed genes between mesoderms of LB-stage WT and *Nipbl*$^{+/-}$ embryos.**

Differential gene expression analysis was performed between mesoderms of LB-stage WT and *Nipbl*$^{+/-}$ embryos using the Mann-Whitney U test. P-values were adjusted for multiple hypotheses by Bonferroni Correction. Gene = gene, Avg_Expr_Wildtype = average expression of indicated gene in Wildtype cells, Avg_Expr_Nipbl+/- = average expression of indicated gene in *Nipbl*$^{+/-}$ cells, SE_Wildtype = standard error of average expression in Wildtype cells, SE_Nipbl+/- = standard error of average expression in *Nipbl*$^{+/-}$ cells, Avg_FC = average fold change in expression of indicated gene in *Nipbl*$^{+/-}$ cells versus WT cells, P_Val = p-value from Mann-Whitney U test, Bon_P_Val = Bonferroni corrected p-value, STRING_Interactor = Y, predicted by STRING to interact with other genes in a network, Nanog_Target = predicted by Fig. 8G to be a target of *Nanog*.

**Data S30. Differentially expressed genes between ectoderms of LB-stage WT and *Nipbl*$^{+/-}$ embryos.**

Differential gene expression analysis was performed between ectoderms of LB-stage WT and *Nipbl*$^{+/-}$ embryos using the Mann-Whitney U test. P-values were adjusted for multiple hypotheses by Bonferroni Correction. Gene = gene, Avg_Expr_Wildtype = average expression of indicated gene in Wildtype cells, Avg_Expr_Nipbl+/- = average expression of indicated gene in *Nipbl*$^{+/-}$ cells, SE_Wildtype = standard error of average expression in Wildtype cells, SE_Nipbl+/- = standard error of average expression in *Nipbl*$^{+/-}$ cells, Avg_FC = average fold change in expression of indicated gene in *Nipbl*$^{+/-}$ cells versus WT cells, P_Val = p-value from Mann-Whitney U test, Bon_P_Val = Bonferroni corrected p-value, STRING_Interactor = Y, predicted by STRING to interact with other genes in a network, Nanog_Target = predicted by Fig. 8G to be a target of *Nanog*.

**Data S31. Differentially expressed genes between endoderms of LB-stage WT and *Nipbl*$^{+/-}$ embryos.**

Differential gene expression analysis was performed between endoderms of LB-stage WT and *Nipbl*$^{+/-}$ embryos using the Mann-Whitney U test. P-values were adjusted for multiple hypotheses by Bonferroni Correction. Gene = gene, Avg_Expr_Wildtype = average expression of indicated gene in Wildtype cells, Avg_Expr_Nipbl+/- = average expression of indicated gene in *Nipbl*$^{+/-}$ cells, SE_Wildtype = standard error of average expression in Wildtype cells, SE_Nipbl+/- = standard error of average expression in *Nipbl*$^{+/-}$ cells, Avg_FC = average fold change in expression of indicated gene in *Nipbl*$^{+/-}$ cells versus WT cells, P_Val = p-value from Mann-Whitney U test, Bon_P_Val = Bonferroni corrected p-value, STRING_Interactor = Y, predicted by STRING to interact with other genes in a network, Nanog_Target = predicted by Fig. 8G to be a target of *Nanog*.

**Data S32. Differentially expressed genes between mesoderms of LB-stage WT and *Nipbl*$^{+/-}$ embryos following projection of WT cells onto *Nipbl*$^{+/-}$ germ layers.**

Differential gene expression analysis was performed between mesoderms of LB-stage WT and *Nipbl*$^{+/-}$ embryos using the Mann-Whitney U test. P-values were adjusted for multiple hypotheses by Bonferroni Correction. Gene = gene, Avg_Expr_Wildtype = average expression of indicated gene in Wildtype cells, Avg_Expr_Nipbl+/- = average expression of indicated gene in *Nipbl*$^{+/-}$ cells, SE_Wildtype = standard error of average expression in Wildtype cells, SE_Nipbl+/- = standard error of average expression in *Nipbl*$^{+/-}$ cells, Avg_FC = average fold change in expression of indicated gene in *Nipbl*$^{+/-}$ cells versus WT cells, P_Val = p-value from Mann-Whitney U test, Bon_P_Val = Bonferroni corrected p-value.

**Data S33. Differentially expressed genes between ectoderms of LB-stage WT and *Nipbl*$^{+/-}$ embryos following projection of WT cells onto *Nipbl*$^{+/-}$ germ layers.**

Differential gene expression analysis was performed between ectoderms of LB-stage WT and *Nipbl*$^{+/-}$ embryos using the Mann-Whitney U test. P-values were adjusted for multiple hypotheses by Bonferroni Correction. Gene = gene, Avg_Expr_Wildtype = average expression of indicated gene in Wildtype cells, Avg_Expr_Nipbl+/- = average expression of indicated gene in *Nipbl*$^{+/-}$ cells, SE_Wildtype = standard error of average expression in Wildtype cells, SE_Nipbl+/- = standard error of average expression in *Nipbl*$^{+/-}$ cells, Avg_FC = average fold change in expression of indicated gene in *Nipbl*$^{+/-}$ cells versus WT cells, P_Val = p-value from Mann-Whitney U test, Bon_P_Val = Bonferroni corrected p-value.

**Data S34. Differentially expressed genes between endoderms of LB-stage WT and *Nipbl*$^{+/-}$ embryos following projection of WT cells onto *Nipbl*$^{+/-}$ germ layers.**

Differential gene expression analysis was performed between endoderms of LB-stage WT and *Nipbl*$^{+/-}$ embryos using the Mann-Whitney U test. P-values were adjusted for multiple hypotheses by Bonferroni Correction. Gene = gene, Avg_Expr_Wildtype = average expression of indicated gene in Wildtype cells, Avg_Expr_Nipbl+/- = average expression of indicated gene in *Nipbl*$^{+/-}$ cells, SE_Wildtype = standard error of average expression in Wildtype cells, SE_Nipbl+/- = standard error of average expression in *Nipbl*$^{+/-}$ cells, Avg_FC = average fold change in expression of indicated gene in *Nipbl*$^{+/-}$ cells versus WT cells, P_Val = p-value from Mann-Whitney U test, Bon_P_Val = Bonferroni corrected p-value.

**Data S35. Network of gene interactions predicted by STRING for genes differentially expressed more than two-fold up or down in germ layers of LB-stage *Nipbl*$^{+/-}$ embryos than WT embryos.**

For each germ layer, genes differentially expressed more than two-fold up or down in Nipbl+/- embryos relative to WT embryos were inputted into STRING. STRING outputted networks of predicted gene interactions.

**Data S36. Expression of *Nanog* in mesodermal cell populations of LB-stage embryos.**

Genotype = genotype of embryo, Embryo = embryo identifier, Cell_Type = cell type in embryo, Nanog = normalized transcript counts for *Nanog* in cell type.

**Data S37. Expression of *Pou5f1* in mesodermal cell populations of LB-stage embryos.**

Genotype = genotype of embryo, Embryo = embryo identifier, Cell_Type = cell type in embryo, Pou5f1 = normalized transcript counts for *Pou5f1* in cell type.

**Data S38. qRT-PCR results for *Nipbl*, *Nanog*, and *Pou5f1* in $Nipbl^{FLEX/+}$ and $Nipbl^{Flrt/+}$ ES cells.**

RNA extracted from 9 clones of $Nipbl^{FLEX/+}$ (WT) and $Nipbl^{Flrt/+}$ ($Nipbl^{+/-}$) ES cells. cDNA was synthesized from extracted RNA. qRT-PCR was performed for *Nipbl*, *Nanog*, and *Pou5f1* for each clone in technical replicates of three, with *Rpl4* as the housekeeping gene.

**Data S39. Differentially expressed genes between whole LB-stage WT and $Nipbl^{+/-}$ embryos.**

Differential gene expression analysis was performed between whole LB-stage WT and $Nipbl^{+/-}$ embryos using the Mann-Whitney U test. P-values were adjusted for multiple hypotheses by Bonferroni Correction. Gene = gene, Avg_Expr_Wildtype = average expression of indicated gene in Wildtype cells, Avg_Expr_Nipbl+/- = average expression of indicated gene in $Nipbl^{+/-}$ cells, SE_Wildtype = standard error of average expression in Wildtype cells, SE_Nipbl+/- = standard error of average expression in $Nipbl^{+/-}$ cells, Avg_FC = average fold change in expression of indicated gene in $Nipbl^{+/-}$ cells versus WT cells, P_Val = p-value from Mann-Whitney U test, Bon_P_Val = Bonferroni corrected p-value, Nanog_Target = predicted by Fig. 8A to be a target of *Nanog*.

**Data S40. Differentially expressed genes between whole CC-stage WT and $Nipbl^{+/-}$ embryos.**

Differential gene expression analysis was performed between whole CC-stage WT and $Nipbl^{+/-}$ embryos using the Mann-Whitney U test. P-values were adjusted for multiple hypotheses by Bonferroni Correction. Gene = gene, Avg_Expr_Wildtype = average expression of indicated gene in Wildtype cells, Avg_Expr_Nipbl+/- = average expression of indicated gene in $Nipbl^{+/-}$ cells, SE_Wildtype = standard error of average expression in Wildtype cells, SE_Nipbl+/- = standard error of average expression in $Nipbl^{+/-}$ cells, Avg_FC = average fold change in expression of indicated gene in $Nipbl^{+/-}$ cells versus WT cells, P_Val = p-value from Mann-Whitney U test, Bon_P_Val = Bonferroni corrected p-value, Nanog_Target = predicted by Fig. 8C to be a target of *Nanog*.

**Data S41. Differentially expressed genes between mesoderms of CC-stage WT and $Nipbl^{+/-}$ embryos.**

Differential gene expression analysis was performed between mesoderms of CC-stage WT and $Nipbl^{+/-}$ embryos using the Mann-Whitney U test. P-values were adjusted for multiple hypotheses by Bonferroni Correction. Gene = gene, Avg_Expr_Wildtype = average expression of indicated gene in Wildtype cells, Avg_Expr_Nipbl+/- = average expression of indicated gene in $Nipbl^{+/-}$ cells, SE_Wildtype = standard error of average expression in Wildtype cells, SE_Nipbl+/- = standard error of average expression in $Nipbl^{+/-}$ cells, Avg_FC = average fold change in expression of indicated gene in $Nipbl^{+/-}$ cells versus WT cells, P_Val = p-value from Mann-Whitney U test, Bon_P_Val = Bonferroni corrected p-value, Nanog_Target = predicted by Fig. 8H to be a target of *Nanog*.

**Data S42. Differentially expressed genes between ectoderms of CC-stage WT and *Nipbl*$^{+/-}$ embryos.**

Differential gene expression analysis was performed between ectoderms of CC-stage WT and *Nipbl*$^{+/-}$ embryos using the Mann-Whitney U test. P-values were adjusted for multiple hypotheses by Bonferroni Correction. Gene = gene, Avg_Expr_Wildtype = average expression of indicated gene in Wildtype cells, Avg_Expr_Nipbl+/- = average expression of indicated gene in *Nipbl*$^{+/-}$ cells, SE_Wildtype = standard error of average expression in Wildtype cells, SE_Nipbl+/- = standard error of average expression in *Nipbl*$^{+/-}$ cells, Avg_FC = average fold change in expression of indicated gene in *Nipbl*$^{+/-}$ cells versus WT cells, P_Val = p-value from Mann-Whitney U test, Bon_P_Val = Bonferroni corrected p-value, Nanog_Target = predicted by Fig. 8H to be a target of *Nanog*.

**Data S43. Differentially expressed genes between endoderms of CC-stage WT and *Nipbl*$^{+/-}$ embryos.**

Differential gene expression analysis was performed between endoderms of CC-stage WT and *Nipbl*$^{+/-}$ embryos using the Mann-Whitney U test. P-values were adjusted for multiple hypotheses by Bonferroni Correction. Gene = gene, Avg_Expr_Wildtype = average expression of indicated gene in Wildtype cells, Avg_Expr_Nipbl+/- = average expression of indicated gene in *Nipbl*$^{+/-}$ cells, SE_Wildtype = standard error of average expression in Wildtype cells, SE_Nipbl+/- = standard error of average expression in *Nipbl*$^{+/-}$ cells, Avg_FC = average fold change in expression of indicated gene in *Nipbl*$^{+/-}$ cells versus WT cells, P_Val = p-value from Mann-Whitney U test, Bon_P_Val = Bonferroni corrected p-value, Nanog_Target = predicted by Fig. 8H to be a target of *Nanog*.

**Data S44. Expression of *Hox* genes in mesoderms of LB- and CC-stage embryos.**

Stage = stage of embryo, Genotype = genotype of embryo, Embryo = embryo identifier, Gene = *Hox* gene, Norm_Transcripts = normalized transcript counts for indicated *Hox* gene.

**Data S45. Expression of *Hox* genes in ectoderms of LB- and CC-stage WT embryos.**

Stage = stage of embryo, Genotype = genotype of embryo, Embryo = embryo identifier, Gene = *Hox* gene, Norm_Transcripts = normalized transcript counts for indicated *Hox* gene.
